# Discovery and mechanistic characterization of a probiotic-origin 3β-OH-Δ^5-6^-cholesterol-5β-reductase directly converting cholesterol to coprostanol

**DOI:** 10.1101/2024.06.04.597308

**Authors:** Urmila Netter, Vishakha Bisht, Amit Gaurav, Rekha Sharma, Avik Ghosh, Vinod Singh Bisht, Kiran Ambatipudi, Hanuman Prasad Sharma, Sujata Mohanty, Shubham Loat, Mihir Sarkar, Kapil Tahlan, Naveen K Navani

## Abstract

On the one hand, cholesterol is a foundational molecule for various structural and biochemical pathways, while elevated cholesterol levels are associated with cardiovascular diseases. Some selected strains of Lactobacilli are also known for modulating cholesterol levels. However, the molecular mechanism of cholesterol transformation by lactobacilli has remained challenging to reveal. This study reports the discovery and role of a microbial 3β-OH-Δ^5-6^-cholesterol-5β-reductase from *Limosilactobacillus fermentum* NKN51, which directly converts cholesterol to coprostanol, thereby resolving the enigma. Protein engineering of the reductase enzyme identified the cholesterol and NADP^+^ interacting amino acids, detailing the catalytic mechanism of 5βChR. Phylogenetic studies emphasize the abundance of 5βChRs in gut commensal lactobacilli, which shares a common ancestor with plant 5β reductases. Meta-analysis results of healthy participant microbiomes underline the significance of 5βChR homologs, while a cohort study reveals an association between higher 5βChR abundance and diabetes. The discovery of the 5βChR enzyme and its molecular mechanism in cholesterol metabolism pave the way for a better understanding of the gut-associated microbiome and the design of practical applications to ameliorate dyslipidemia.

## Introduction

Cholesterol is a vital animal sterol, serving essential structural and metabolic functions. However, high cholesterol levels are a critical risk factor for cardiovascular diseases (CVDs) and atherosclerosis, which can lead to heart failure, obesity, diabetes, fatty liver, and Alzheimer’s disease [1–3]. Gut microbes can transform dietary cholesterol into coprostanol by direct stereospecific reduction of the Δ5 double bond or via forming intermediates such as cholestenone and coprostanone **(Fig. S1)** [4]. The short-chain dehydrogenase (SDRs) enzyme family has recently been shown to contribute to gut cholesterol transformation [5]. The commensal gut microflora possesses SDRs that facilitate the conversion of hydroxyl groups in alcohols, carbohydrates, and xenobiotics from their enol to keto forms [6–8]. The SDRs contain conserved motifs spanning the Rossmann fold for NADP^+^/NAD^+^ cofactor binding and a catalytic tetrad comprising serine, tyrosine, lysine, and asparagine residues [9, 10]. Eight 5β reductases classified as SDRs have been thoroughly studied in plants; however, their role in bacteria has received considerably less attention[11, 12]. Only recently has the mechanism of cholesterol metabolism been described for the obligatory gut anaerobes *Eubacterium coprostanoligenes, Bacteroides thetaiotaomicron,* and *Oscillibacter* species [5, 13, 14]. The genome of *E. coprostanoligenes* encodes the cholesterol dehydrogenase enzyme ‘Intestinal steroid metabolism A’ (IsmA), which converts cholesterol into cholestenone and coprostanone into coprostanol [5]. However, information on the steps involved in transforming cholestenone into coprostanone is lacking. On the other hand, *B. thetaiotaomicron* exhibits unique sulfotransferase activity that converts cholesterol into cholesterol sulfate [13]. Furthermore, *Oscillibacter* species convert cholesterol to cholestenone, cholesterol glucoside, and hydroxycholesterol [14]. Overall, coprostanol, cholesterol sulfate, cholesterol glucoside, and hydroxycholesterol are excreted, thereby regulating cholesterol homeostasis [5, 13, 14]. These anaerobes do not metabolize cholesterol directly. Although the direct conversion of cholesterol to coprostanol in the gut was suggested as early as 1954, the underlying mechanism of this transformation remains a mystery [4, 15, 16]. Selected strains of probiotic lactobacilli display potential for cholesterol metabolism. However, the specific mechanisms involved in cholesterol reduction have not yet been elucidated [17–22].

We report a novel cholesterol-specific microbial 5β reductase (5βChR) from *Limosilactobacillus fermentum* NKN51 isolated from fermented Himalayan yak milk. Our study provides mechanistic evidence of the direct conversion of cholesterol to coprostanol by 5βChR through its 3β-OH-Δ^5-6^-cholesterol-5β-reductase activity. The biochemical and structural analysis of 5βChR identify and characterize cholesterol metabolism in lactobacilli. Furthermore, phylogenetic and metagenomic analysis suggests the prevalence of 5βChR in microbes and its association with human physiology. Our findings addressed a primarily unexplored frontier of microbial cholesterol metabolism in *L. fermentum* NKN51 by identifying a novel enzyme class from the SDR family, establishing a path for unveiling the metabolic capabilities of one of the most popular probiotic bacteria.

## Results

### *L. fermentum* NKN51 exhibit 3β-OH-Δ^5-6^-cholesterol-5β-reductase activity

To understand the mechanism of cholesterol metabolism in lactobacilli, we performed cholesterol depletion assays with a battery of Lactobacillus spp. isolated from an ethnic fermented yak milk cheese (churppi) prepared by the native population in the Western Indian Himalayas. Among other strains, *L. fermentum* NKN51 depleted the maximum (∼64%) cholesterol in the media **(Fig. S2)**. Thin layer chromatography (TLC) and Liquid chromatography-mass spectrophotometry (LC-MS) analysis of *L. fermentum* NKN51 whole-cell lysate confirmed the conversion of cholesterol to coprostanol **(Fig. S3, 1*a*)**. We could not detect cholestenone or coprostanone in these assays, suggesting that *L. fermentum* NKN 51 might directly convert cholesterol to coprostanol in NADP^+^-dependent but oxygen-independent conditions **(Fig. S4, S5)**. Furthermore, a known cholesterol SDR from *E. coprostanoligenes* HL ATCC51222[5] (WP_078769004.1) was used as a query against the *L. fermentum* NKN51 draft genome, which highlights the putative cholesterol dehydrogenase protein with ∼30% identity in NKN51. The putative cholesterol dehydrogenase was cloned, and the 29.5 kDa overexpressed protein was purified by immobilized affinity chromatography (IMAC) **(Fig. S6)**. The recombinant purified protein converted cholesterol to coprostanol using NADP^+^ as a cofactor instead of NAD^+^, validating the activity we initially observed with *L. fermentum* whole-cell lysate assays **(Fig. 1*a*)**. To determine the binding affinities of protein with its cofactor and substrates, we performed isothermal titration calorimetry (ITC) analysis. NADP^+^ had an affinity for protein with *K_d_* 6.88 µM and the protein-NADP^+^ complex bound with cholesterol with *K_d_* of 4.66 µM **(Fig. 1*b*, *c*)**. However, protein-NADP^+^ did not display an affinity for cholestenone or coprostanone, substantiating that neither of these compounds is cognate substrates of the enzyme **(Fig. S7)**. Unlike typical microbial SDRs, the cholesterol dehydrogenase remains primarily a dimeric protein in solution, confirmed by size exclusion chromatography **(Fig. S8)**. The substrate specificity experiments validated that protein can specifically catalyze the direct conversion of cholesterol to coprostanol by saturating Δ^5^ double bond with the OH group at the 3β position **(Fig. S9, 1*d*)**. Hence, these distinctive catalytic features of the protein ascribed it to a newly identified class within microbial SDRs, i.e., 3β-OH-Δ^5-6^-cholesterol-5β-reductase (5βChR). Enzyme-substrate kinetic studies confirmed that 5βChR obeys the Michaelis–Menten equation, with a *K_m_* of 19.09 µM and *V_max_* of 15.35x10^-6^ µmol sec^-1^ nmol^-1^ **(Fig. S10)**. At the genetic level, 5βChR transcript expression levels were ∼1.5-fold increased when the NKN51 strain was grown in the presence of cholesterol, signifying that the 5βChR genetic locus was induced upon cholesterol supplementation (*p*=0.0022) in the native host **(Fig. 1*e*)**. The biophysical characterization and *in-vitro* assay of 5βChR map the molecular basis of cholesterol metabolism in *L. fermentum* NKN51.

**Fig. 1.**
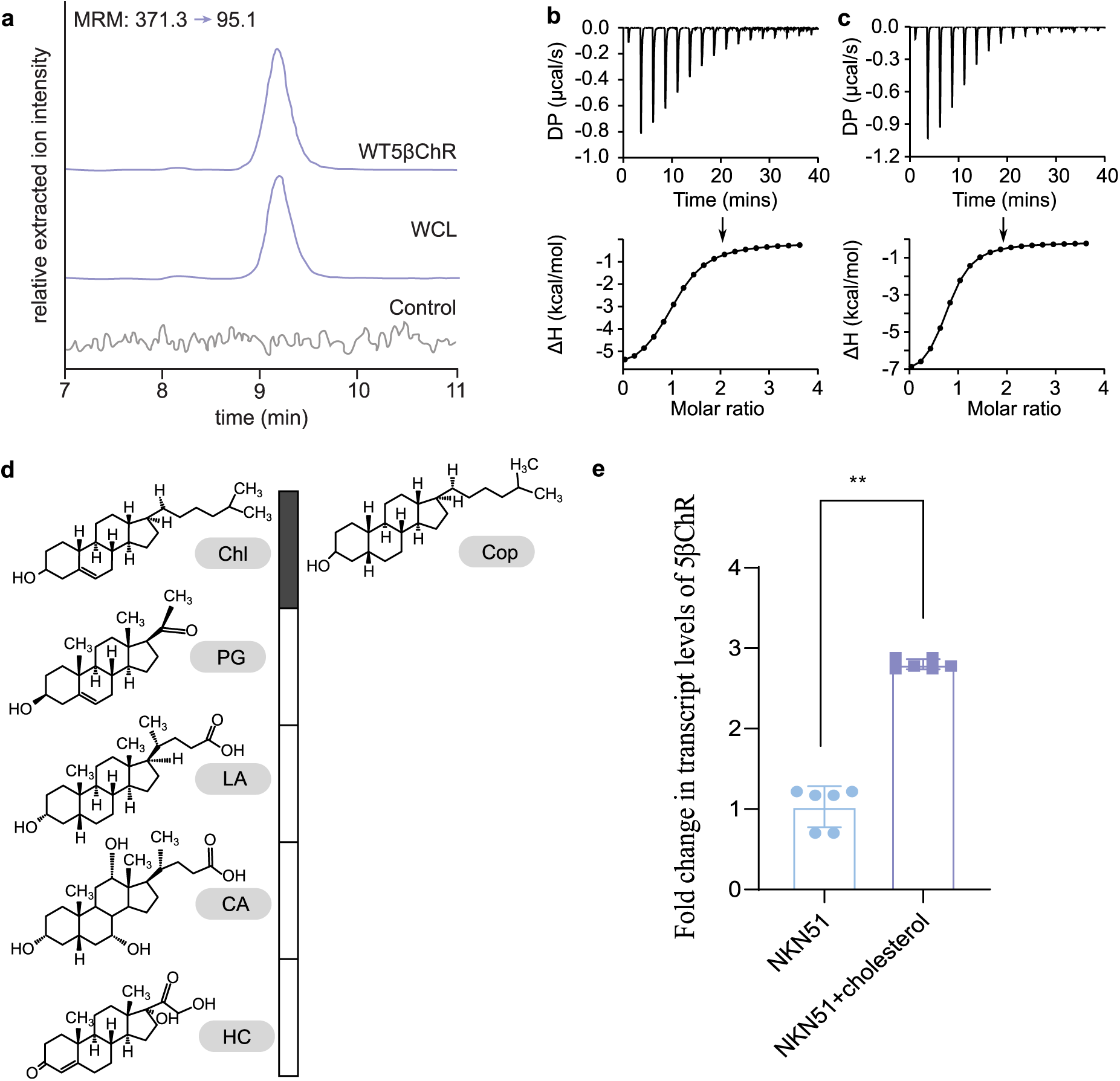
*L. fermentum* NKN51 harbors a cholesterol dehydrogenase that can convert cholesterol to coprostanol. **(a).** Extracted ion chromatograph shows the reaction product of *L. fermentum* NKN51 whole-cell lysate (WCL) and purified cholesterol-5β-reductase (5βChR) incubated with NADP^+^ (100 µM) and cholesterol (100 µM) for 12 h. **(b).** Isothermal Titration Calorimetry (ITC) data displaying binding affinity of the 5βChR (20 µM) for NADP^+^ (1 mM). **(c).** The titration curve provides the interaction between the 5βChR + NADP^+^ complex and cholesterol (500 µM), where DP stands for differential power, and ΔH represents the enthalpies (normalized with the control). **(d).** The bar graph indicates the conversion of cholesterol to coprostanol (grey block) by 5βChR under the same conditions. The other structurally similar compounds are not metabolized into their respective metabolites. Abbreviations: PG stands for pregnenolone, LA-lithocholic acid, CA-cholic acid, and HC-hydrocortisone. Spectra analysis of this substrate specificity assay is given in Supplementary Figure 8. **(e).** The transcript levels of 5βChR in a cholesterol-rich medium compared to a medium without cholesterol at 24 h under anaerobic conditions show ∼1.5-fold change. In the graph, each point represents a biological replicate, with the centre bar representing the mean and the error bars representing the standard deviation. ** denotes the *p-*value-0.0022 calculated by the Mann-Whitney test.

**Fig. 2.**
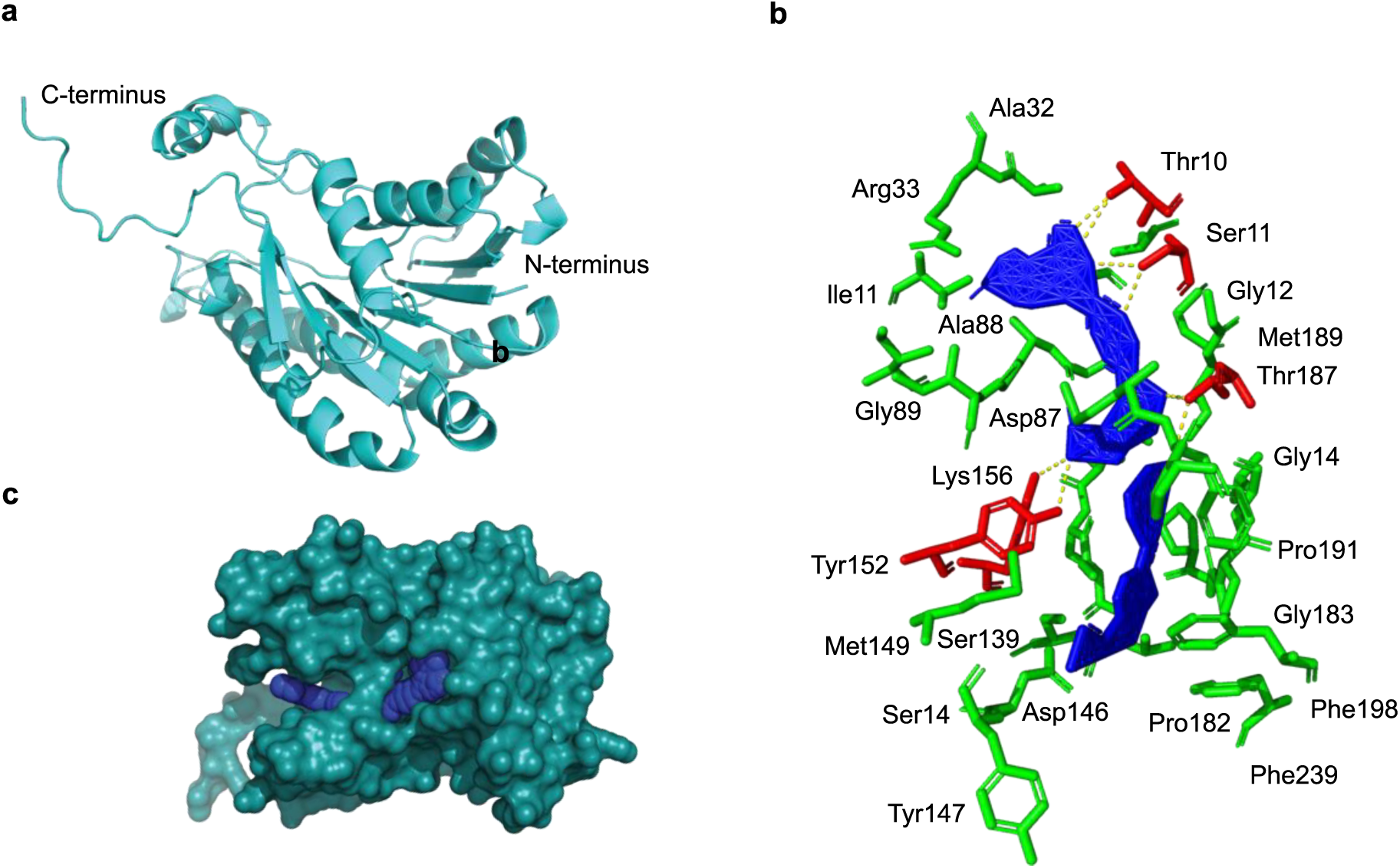
Structure and mutation analysis of 5βChR for ligand binding and comparison study with the plant 5β reductase. **(a).** Alphafold2 predicted a top-ranked 5βChR structure with high confidence (>90%), a high preresidue confidence score (>90%), and a low domain position confidence, and visualized by PyMOL. **(b).** The surface representation of 5βChR shows the active binding pocket of cholesterol and NADP^+^ (blue). **(c).** The molecular docking results of 5βChR-NADP^+^-cholesterol depict the interacting amino acids. The Cofactor (NADP^+^) binding pocket highlights the ionic interaction (red) (indicated by dashed lines). The amino acids involved in hydrogen bonding are Thr10, Ser11, and Tyr152, while Lys156, Ala32, Arg33, Gly89, Ile89, Ser139, Asp146, Met149, Ser11, Ile90, and Ala9 showcase involvement in van der Waals bonds (green). Protein-substrate (5βChR-cholesterol) complex shows hydrophilic interaction with Thr187, whereas the hydrophobic forces are present between Pro182, Pro184, Met189, Phe190, Phe198, and Phe238.

### 5βChR interacts with cholesterol via C-terminus and NADP+ by N-terminus

The 5βChR AlphaFold tertiary structure depicts conserved motifs of the SDR superfamily, including the N-terminus dinucleotide binding Rossmann fold consisting of seven central β-strands flanked by six α-helices and a catalytic tetrad (NSYK) **(Fig. 1*a-b*)**. Molecular docking results predicted that the cholesterol was bound in the C-terminus pocket of the protein. Cholesterol steroid ring-1 interacts with methionine (Met189) and phenylalanine (Phe190) residues of 5βChR through hydrophobic forces. An electrostatic interaction was seen between the threonine (Thr187) and the hydroxyl group of cholesterol. NADP^+^ was linked to the N-terminus of 5βChR via the conserved glycine-rich (Gly 8, 9, 12, and 14) motif. The tyrosine (Tyr152), lysine (Lys156), serine (Ser11), and threonine (Thr10 and Thr187) amino acid residues contributed to a total of eight polar interactions with NADP^+^. Furthermore, 5βChR is directed towards the nicotinamide oxygen (O=C-NH) by Tyr152 and Lys156 amino acid residues. The N-terminus Ser11 amino acid residue interacts with the diphosphate group of NADP^+^. On the other hand, the ribose phosphate group of the NADP^+^ coordinates with the Thr10 amino acid residue of 5βChR **(Fig. 1*c*)**. The superimposition of 5βChR and its structural homologs (SDR reductase, sorbitol dehydrogenase, and putative oxidoreductase RV2002) revealed a shared topological arrangement with root square mean (r.m.s) deviations of 1.3, 1.3, and 1.2 Å, respectively and confirmed that 5βChR belongs to SDRs **(Fig. S11)**.

### Identification of catalytic residues for NADP^+^ and cholesterol binding in 5βChR

Using site-directed mutagenesis, we elucidated the role of the catalytic tetrad (NSYK) and other conserved residues of 5βChR. Since the C-terminus of SDRs was reportedly involved in substrate binding,[6] we mutated the C-terminus of the protein to investigate its role in cholesterol binding **(Fig. S12)**. TLC and LC-MS experiments of 5βChR mutants with NADP^+^ and cholesterol revealed the importance of Ser139, Tyr152, Thr187, Met189, and Phe190 amino acid residues in cholesterol conversion to coprostanol **(Fig. S13, 3*a*)**. Furthermore, ITC data revealed that the S139A and Y152A mutants lost NADP^+^ binding but retained the cholesterol binding, with *K_d_* values of 24.8 µM and 4.44 µM, respectively. On the other hand, T187A, M189A, and F190A mutants retained NADP^+^ binding (*K_d_* 8.77 µM, 7.49 µM, and 5.2 µM, respectively) **(Fig. 3*b*)** but lost the binding ability to cholesterol. Thus, the C-terminus of 5βChR is important for substrate interaction, and the C-terminus of SDRs is indeed necessary for substrate interaction[6]. Other mutations, such as N136A, K156A, H131A, Y170A, E172A, and T187A, did not affect the ability to convert cholesterol to coprostanol, thus indicating their nonsignificant role in enzyme activity **(Fig. 3*a*)**. Circular dichroism spectral analysis of the wild-type and mutant 5βChR enzymes did not predict any change in protein structure. Thus indicating that the changes in cofactor and substrate binding were not due to perturbations in the secondary structure of the 5βChR enzymes. **(Fig. S14)**. Since plant and microbial 5β reductases belong to the SDR superfamily, we performed an in-depth analysis of the structural attributes of plant and microbial 5β reductases. A systematic comparison of the NADP^+^ binding pocket of microbial 5βChR with the plant origin 5β-progesterone reductases (5β-POR) (PDB: 2V6G) was accomplished [11]. It revealed that the NADP^+^ was bound to the Rossmann fold in plant and bacterial 5β reductase. However, the lysine residue of a conserved catalytic tetrad of SDRs does not interact with the O-2’ or O-3’ of nicotinamide ribose in either case **(Fig. 4)** [9, 23].

**Fig. 3.**
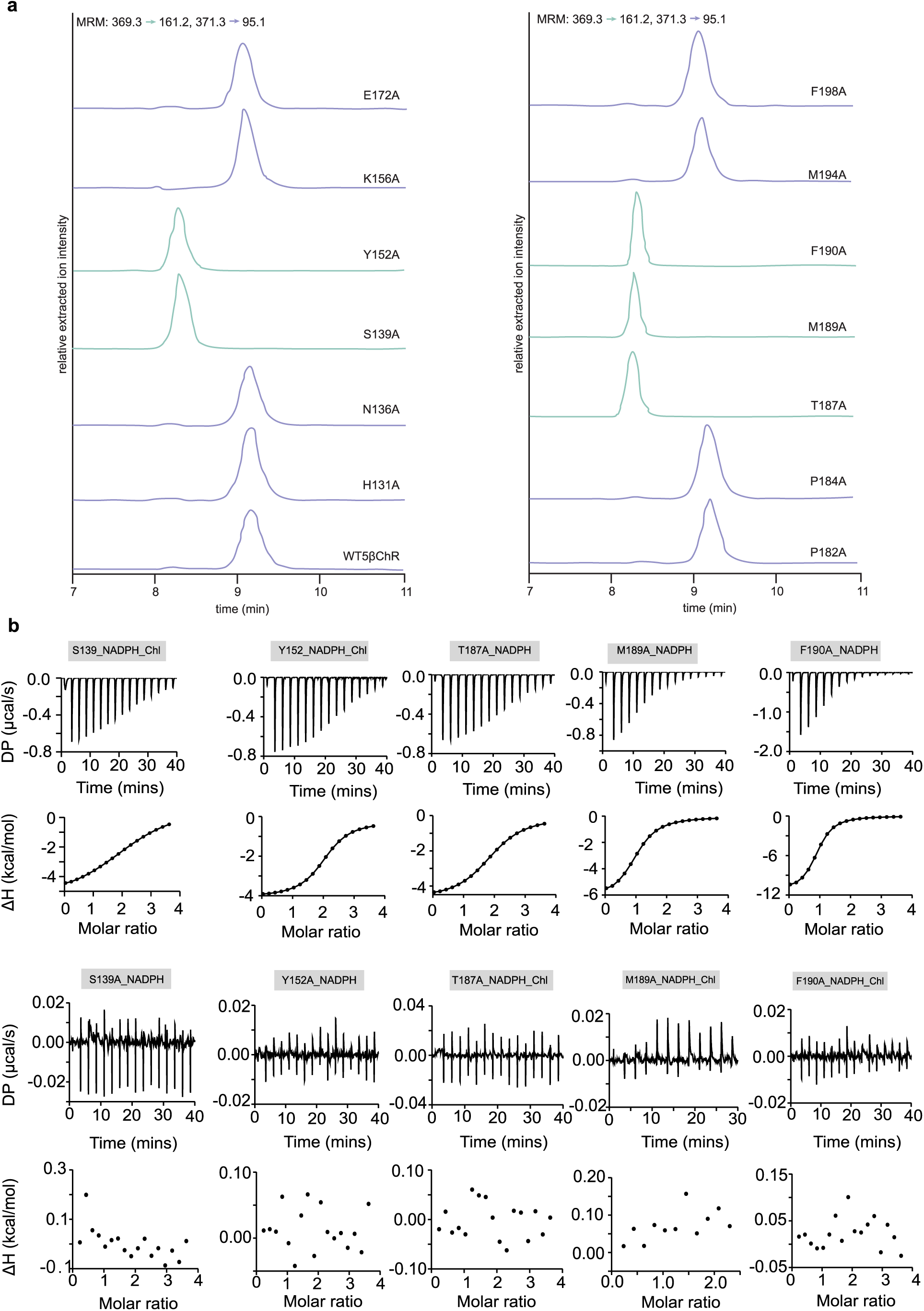
Characterization of 5βChR mutants. **(a).** The LC-MS data is represented by the extracted ion chromatographs of 5βChR mutants supplemented with NADP^+^ (100 µM) and cholesterol (100 µM) under similar conditions. **(b).** The normalized enthalpy curves represent the interaction profile of 5βChR mutants with their ligands. S139A and Y152A mutants have an affinity for cholesterol and lost binding for NADP^+^. T187A, M189A, and F190A mutants do not bind to cholesterol. In contrast, they retained an affinity for NADP^+^.

**Fig. 4.**
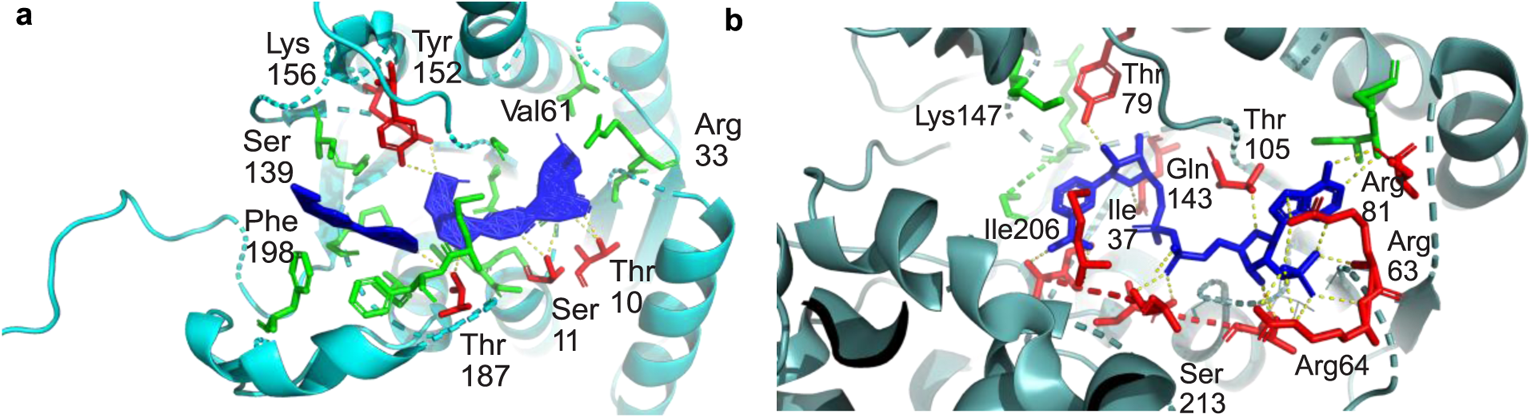
Comparison of the active sites of 5βChR and 5β-POR with their respective substrates. The comparison study of the active site of 5βChR and plant 5β reductase (5β-POR) with their respective substrates. The stereo view of 5βChR complexed with NADP^+^ is mediated by tyrosine (Tyr152) and serine (Ser139) of the conserved tetrad. In 5β-POR - two arginine residues (Arg63 & Arg64) stabilise NADP^+^ interaction. 5β-POR has two Rossmann folds (glycine residues) (GXXGXXG), whereas 5βChR harbors a single N-terminus fold, contributing to NADP^+^ binding.

### 5βChRs is a prevalent enzyme class in microbes and shows its association with human physiology

Since 5βChRs are a novel class of enzymes discovered in bacteria, we focused on assessing the diversity and abundance of these enzymes within Lactobacillaceae and overall in eubacteria. A local database of 256 lactobacilli was screened to determine the prevalence of 5βChRs in the genome. The maximum likelihood method confirmed the presence of the 5βChR sequence in 84 lactobacilli genomes, which were further divided into six major clades. Most of these representatives (*Lactiplantibacillus plantarum, Limosilactobacillus mucosae, Levilactobacillus parabrevis*, *Latilactobacillus sakei*, *Latilactobacillus curvatus*, *Ligilactobacillus salivarius,* and *Levilactobacillus brevis*) were associated with a healthy commensal gut community **(Fig. 5)** [24, 25]. We traced the evolutionary relationship of microbial 5βChRs with plant 5β reductases using human homologs as an outgroup, which revealed that plant and microbial 5β reductases originated from a common ancient ancestor **(Fig. S15)**. To assess the distribution of 5βChRs in different microbial taxa, we retrieved 5000 5βChRs and interpreted the data using Cytoscape. 5βChR homologs were grouped into 33, 138, and 269 clusters with threshold values for sequence identity (e-value) of 10^−60^, 10^-70^, and 10^-80^, respectively **(Fig. S16, S17*a*)**. A threshold value of 10^−80^ was chosen to map the 5βChR homologs to ensure similar functional and structural aspects, which led to the grouping of proteins into 269 distinct clusters. The phylum-level taxonomic classification of the respective clusters was considered to assess the prevalence of microbial 5βChRs. Across the different clusters, 5βChRs were predominantly present in Pseudomonadota, Actinomycetota, Planctomycetota, Chloroflexota, Cyanobacteria, Acidobacteriota, Bacillota, and Candidatus omnitrophota **(Fig. S17*b*, S18).** The SSN analysis indicated that inhabiting different niches belonging to different phyla harbored 5βChR homologs, possibly providing some benefit to their respective hosts. To understand the importance of the aforementioned 5βChR clusters, we assessed the data through ShortBRED, which consists of protein classes in 380 high-quality, healthy human metagenome sequences (from the human microbiome project), covering samples from the different regions of the body. Using the EFI-Computationally Guided Functional Profiling (EFI-CGFP) tool, we identified 20 clusters of 5βChR homologs in the Human Microbiome Project data **(Fig. S19)**. Clusters 3, 9, and 11 were abundant in the oral microbiome (supragingival plaque, buccal cavity, and tongue dorsum). In contrast, clusters 2, 3, 6, 9, 11, 52, 83, 104, 119, 122, 128, 140, 156, 172, 185, and 24 were the dominant clusters in the fecal samples, indicating the presence of 5βChR homologs in anaerobes. Among the bacteria-dominated clusters, 16 and 139 had more significant distributions in the posterior fornix (vaginal microbiome) and anterior nares, respectively **(Fig. 7, S20)**. The comprehensive analysis of the 5βChR family in the human gut inhabitants highlights this enzyme’s prevalence and potential role in maintaining cholesterol homeostasis.

**Fig. 5.**
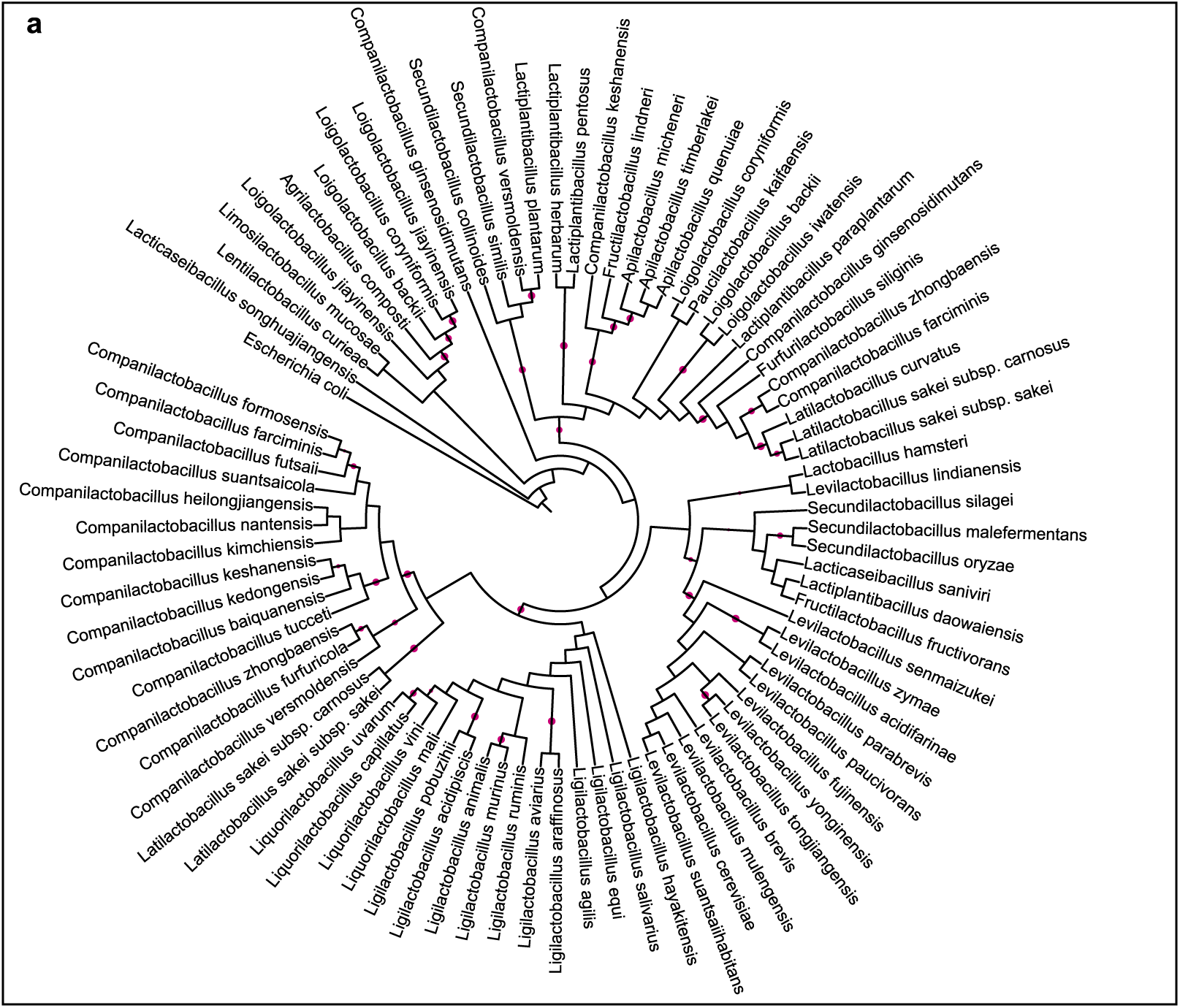
Mapping the prevalence of 5βChR in lactobacilli in gut lactobacilli. The cladogram of 5βChR protein across 256 lactobacilli depicts its abundance in 84 strains. This study used the PROTCATWAG model in RAxML and rooted using the branch leading to *Lacticaseibacillus songhuajiangensis* and *Escherichia coli* as the outgroups.

### 5βChR has an inverse association with diabetes

We explore the correlation of 5βChR with diabetes because diabetes can induce changes in lipid levels, leading to diabetic dyslipidemia, which exacerbates risk factors for CVDs [26–28]. Analyzing the relationship between 5βChR and diabetes using Chinese-type 2 diabetes shotgun data (PRJNA855128). The two independent metagenomic cohorts included diabetic individuals (n=10) and healthy controls (n=10). The interpretation of the data confirmed that 5βChRs exhibited a significantly greater prevalence in healthy individuals than in those with diabetes (*p*=0.0213) **(Fig. 6*b*)**. These findings corroborate the inverse relationship between 5βChR and diabetes in this cohort study, indicating that 5βChR expression might reduce the risk of premature coronary heart disease and stroke in patients with diabetic dyslipidemia [29].

**Fig. 6.**
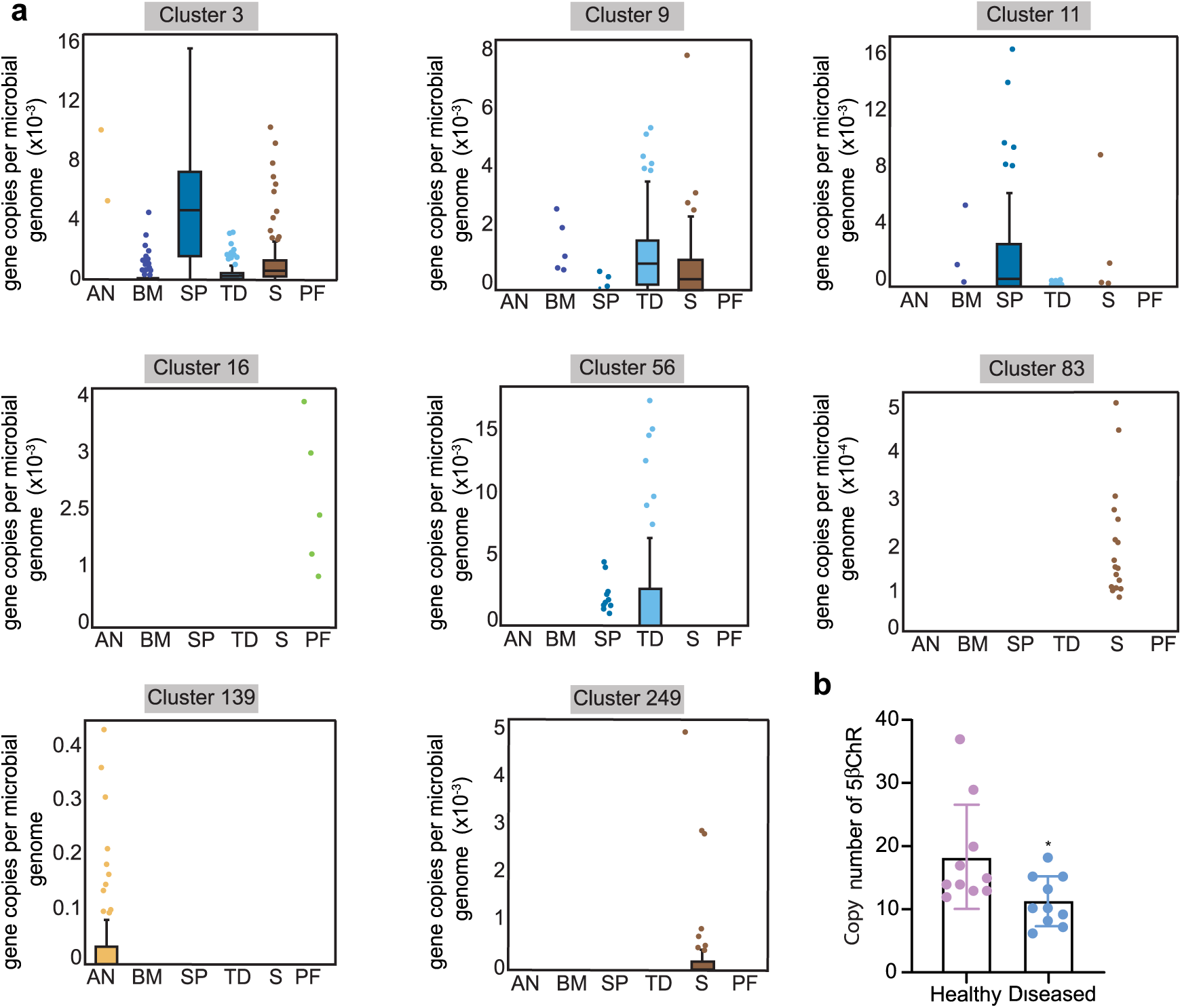
Meta-analysis depicting the abundance of 5βChR homologs in healthy microbiomes and comparative analysis in a human diabetic cohort study. **(a).** The box-plot graphs highlight the presence of 5βChR homologs in various body sites. Clusters 3, 9, 11, and 56 dominate the oral microbiome (supragingival plaque, buccal cavity, and tongue dorsum) and stool samples. Cluster 16 is predominant in vaginal samples. The enzymes from other clusters 139 showed higher distribution in anterior nares samples of human microbiomes. Further, clusters 83 and 249 are abundant in stool samples, indicating their abundance in the gut. **(b).** 5βChR has a lower abundance among diabetic patients than healthy individuals in the shotgun metagenomic data of a Chinese type 2 diabetes cohort study. Individuals with 5βChR high copy number are predominantly reported as healthy. The points in the graph represent the number of individuals in each category. \**p* value-0.0213 is determined by the Mann-Whitney test.

**Fig. 7.**
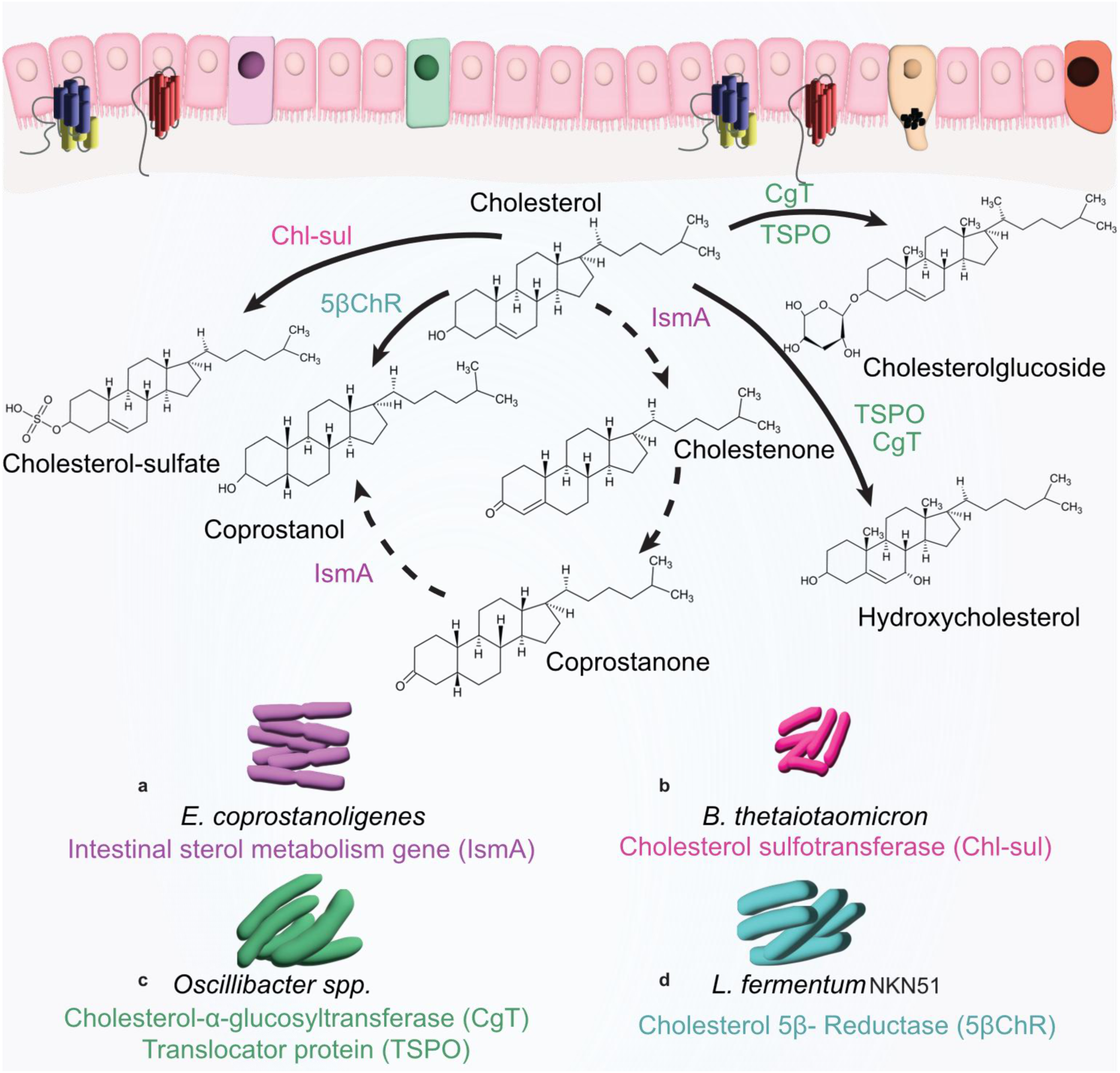
A comprehensive representation of microbial cholesterol transformation in the human gut. Various gut microbes transform cholesterol in the gut via direct and indirect pathways. **(a).** A cholesterol dehydrogenase (ismA) of *Eubacterium coprostanoligenes* is reported to catalyze the first and last step of the indirect pathway of cholesterol metabolism. **(b).** The gut cholesterol is also converted to cholesterol-sulphate with the help of a cholesterol sulfotransferase encoded by *Bacteroidetes thetaiotaomicron.* **(c).** A group of enzymes, cholesterol-α-glucosyltransferase, and translocator protein expressed by *Oscillibacter* species transformed cholesterol into cholesterolglucoside and hydroxycholesterol while **(d).** A comprehensive study demonstrates that 5βChR expressed by *Limosilactobacillus fermentum* NKN51 can directly convert cholesterol to coprostanol by a cholesterol-5β-reductase.

## Discussion

The gastrointestinal tract is a site for cholesterol metabolism and regulation of blood cholesterol levels. Although cholesterol metabolism by lactobacilli has been known for almost 50 years, the exact mechanism has remained elusive [16]. Recently, three independent studies have reported the mechanism of cholesterol metabolism by different bacterial gut residents. Still, the role of lactobacilli has not been addressed in either of the reports[5, 13, 14]. Here, we discovered and characterized a novel microbial 5β-reductase under the SDR superfamily, which catalyzes the direct transformation of cholesterol to coprostanol **(Fig. 7)**. Molecular docking observations of the cofactor and substrate with the wild-type and mutant 5βChRs identified the critical residues for NADP^+^ binding and revealed novel residues important for cholesterol binding to the protein. The structural and mutational analysis of 5βChR represents its similarity with microbial SDRs and plant 5β reductase, categorizing it into a new class of SDRs. Furthermore, evolutionary analysis of plant and microbial 5βChR enzymes suggested that both are paraphyletic groups and confirmed the presence of 5βChR in gut lactobacilli. Shotgun data analysis highlights the low levels of 5βChR transcripts in diabetic patients. Metagenomic mining revealed that the representatives of 5βChRs are present in *Clostridia, Eubacteria, Oscillopsia, Peptostreptococcus, Fusobacteria Bifidobacteria, and Lactobacillus*. Our findings suggest that 5βChR homologs are also acquired by uncharacterized bacterial classes, which may be explored further. The prevalence of 5βChR in the microbiomes of healthy individuals indicates the critical role this enzyme can play in designing strategies to regulate lipid homeostasis. We believe this study can pave the way for developing next-generation probiotics that can be explored to formulate functional foods to ameliorate dyslipidemia.

## Materials and methods

### In vitro cholesterol reduction assay

*Limosilactobacillus fermentum* NKN51,[30] *Lacticaseibacillus casei* NKN344, and *Lacticaseibacillus casei* NKN347 were fermented food isolates from our laboratory. For the cholesterol estimation assay, the secondary culture was inoculated in sterile anaerobic Man, Rogosa, and Sharpe (MRS) media (Difco Laboratories, USA) supplemented with cholesterol (Sigma Aldrich, USA) (100 µg/ml) and sodium thioglycolate (STG) (0.1%) (Tokyo Chemical Industry, China) and incubated at 37°C for 24 h. After incubation, the samples were centrifuged at 13,000 rpm for 5 min. The supernatant was subjected to a cholesterol estimation assay using the Rudel and Morris methods [31]. This procedure was performed in triplicate, and the average values were plotted. For whole-cell lysate assay, a 12 h *L. fermentum* NKN51 culture was pelleted by centrifugation at 4500 rpm for 20 min at 4°C, resuspended in phosphate-buffered saline (PBS) (1x) (Sigma Aldrich, USA), and lysed with a Branson probe sonicator 102C (CE) (Thermo Fisher Scientific, USA) and centrifuged at 13,000 rpm for 30 min at 4°C. The 500 µL supernatant (whole cell lysate) with cholesterol (100 µM) and NADP^+^ (100 µM) (Sigma Aldrich, USA) was incubated at 37°C for 12 h. Further, the sterols were extracted by adding double the volume of hexane (Merck, USA), followed by vigorous mixing. The microcentrifuge tube was left for phase separation for an hour at room temperature. After the phase separation, the upper organic layer was transferred into a fresh microcentrifuge tube and evaporated by a refrigerated centriVape concentrator (Labconco, USA). The sterols were resuspended in 10µl of methanol (Merck, USA) and separated by TLC (explained later). The reaction products were also checked via LC-MS (details are provided in the LC-MS instrument section).

### Cloning of the putative cholesterol dehydrogenase gene

The genomic DNA of *L. fermentum* NKN51 was isolated using the HiPurA® Bacterial Genomic DNA Purification Kit (Himedia, India). The 5βChR gene was amplified using Phusion High Fidelity DNA polymerase (Thermo Fisher Scientific, USA) with the primers listed in **Table S1**. Furthermore, the PCR product and pET28(a) were double digested with NdeI (Thermo Fisher Scientific, USA) and XhoI (Thermo Fisher Scientific, USA). The digested products were ligated using T4 DNA ligase (Thermo Fisher Scientific, USA) and transformed into chemically competent *E. coli* cells. The transformants were checked via double digestion of the plasmid, followed by sequencing.

### Site-directed mutagenesis of the putative cholesterol dehydrogenase gene

WT5βChR carrying plasmid was used to construct its mutants with the Q5® Site-Directed Mutagenesis Kit (New England Biolabs, USA). Primers containing the desired mutation were designed with the NEBase Changer webpage tool (https://nebasechanger.neb.com/). The primers used are listed in **Table S1**, and the transformants were confirmed by sequencing.

### Overexpression and purification of the N-His terminal tagged WT and mutant 5βChR proteins

Confirmed clones containing WT and mutant 5βChR were transformed into ultracompetent BL21AI (arabinose induction) (Thermo Fisher Scientific, USA) cells for protein overexpression. 5βChR protein (29.5 kDa) overexpression was optimized with IPTG (0.05 mM) (Tokyo Chemical Industry, China) and arabinose (0.2%) (Himedia, India). For protein purification, the cells were grown in Luria–Bertani broth (Difco Laboratories, USA) supplemented with kanamycin (50 µg/ml) (Sigma Aldrich, USA) overnight at 37°C with shaking at 180 rpm. The secondary culture was inoculated with a primary culture of WT or the 5βChR mutant (1%) and incubated at 37°C with shaking at 180 rpm until the optical density (OD_600_) reached ∼0.8. The bacterial cultures were induced with IPTG (0.05 mM) and 0.2% arabinose (0.2%) and incubated at 18°C with shaking at 180 rpm for 18 h. The cells were harvested at 4500 rpm for 20 min at 4°C and resuspended in 20 mL of buffer A (50 mM Tris-Cl, 500 mM NaCl 5% glycerol) (pH 7.8). The resuspended cells were lysed with a Branson probe sonicator 102C (CE) and then centrifuged at 4500 rpm for 30 min at 4°C. The clarified supernatant was loaded into the buffer A saturated HisTrap column (AKTA Pure-50 L; GE Healthcare). Purified protein fractions were eluted with Buffer B (50 mM Tris-Cl, 500 mM NaCl, 500 mM imidazole 5% glycerol) (pH 7.8) and analyzed via SDS‒PAGE. Furthermore, the protein was dialyzed in dialysis buffer (20 mM Tris-Cl, 300 mM NaCl 5% glycerol) with a 10 kDa membrane (Pall Corporation Centrifugal Devices, USA) for 8h intervals. The protein concentration was checked via SDS‒PAGE, and the proteins were stored in aliquots at -80°C. Purified protein fractions were subjected to further assays, which were discussed later.

### Thin-Layer Chromatography of Sterols

We checked the enzyme activity by TLC in the presence of cofactor and substrate. The reaction mixture was prepared with an enzyme (WT and mutants) (10 nm), NADP^+^ (100 µM), and substrate (100 µM) (cholesterol, cholestenone, and coprostanone) in dialysis buffer was incubated at 37°C for 12 h. These reaction mixtures were subjected to sterol extraction (as explained above) and analyzed via TLC. The resuspended sterol was spotted on a pretreated TLC plate (silica gel IB2-F flexible 20 × 20 cm) with methanol at 100°C for 30 min. The sterols were separated with a mobile phase (benzene: chloroform) (90:10) (Merck, USA). After being treated with 15% sulfuric acid in methanol, the plate was visualized and incubated at 110°C for 10 min.

### LC-MS instrumentation conditions for the measurement of sterols

The enzyme-cofactor-substrate analysis was conducted on ultra-high performance liquid chromatography (UHPLC) coupled with tandem mass spectrometry. The Exion LC UHPLC was equipped with an online degasser, binary pump, computer-controlled sample cooler, and column heater. Mass spectrometric analysis was on QTRAP 6500+ (Sciex, MA). UHPLC and mass spectrometer parameters were controlled using Analyst (ver 1.7.1; Sciex, MA, USA). The chromatographic separation was achieved on the Discovery C8 column (150X4.6 mm, 5µm; Supelco, Sigma-Aldrich, USA). An isocratic mobile phase consisting of ultra-pure water (A) and acetonitrile (B) at a ratio of 10:90 (A: B) was used for the separation. The flow rate was maintained at 1 mL/min, and the column oven was kept at ambient. A 20 µL of sample was injected into the column. The mass spectrometric analysis was done by an atmospheric pressure chemical ionization (APCI) probe operated at positive ion mode. The multiple reaction monitoring (MRM) mode was used for the quantification of sterols with the following mass transitions: cholesterol (369.3 →161.2), and coprostanol (371.3 →95.1).

### qPCR of the 5βChR transcript

The chloroform-phenol (VWR, USA) extraction method was used for total RNA purification from triplicate *L. fermentum* NKN51 cell pellets grown with or without cholesterol for 24 h. DNase-treated RNA samples were used to prepare cDNA using the PrimeScript 1st strand cDNA Synthesis Kit (Takara, Japan). The 5βChR transcript was quantified by real-time PCR with PowerTrack SYBR Green Master Mix (Applied Biosystems, USA). The expression was normalized to the 16S rRNA gene expression. The primers used in the analysis are listed in **Table S1**.

### Gel filtration chromatography of 5βChR

5βChR gel filtration chromatography (column: Hiload^TM^ 16/600 Superdex^TM^ 200 pg) was performed on an AKTA pure-50 L chromatography system (GE Healthcare) at 4°C. The concentrated, purified protein (10 mg/mL) was loaded onto an analytical column equilibrated with Tris-Cl (20 mM) and NaCl buffer (200 mM) at pH 7.8 and eluted at a flow rate of 1 ml/min. A protein standard consisting of thyroglobulin bovine, γ-globulins, ovalbumin, ribonuclease A type I, and p-aminobenzoic acid (pABA) (Merck, India) was used to determine the native molecular mass of 5βChR.

### Enzyme kinetic assay

The enzyme activity of the purified protein was measured spectrophotometrically by monitoring NADP^+^ levels at 340 nm. The standard reaction mixture comprised sodium acetate trihydrate buffer (20 mM) (pH 7.8), NADP^+^ (100 µM), and the enzyme (20 nm), and the substrate concentration varied from 10 to 300 µM. GraphPad Prism was used to plot the data using the Michaelis–Menton saturation curve and Lineweaver–Burk plots to calculate the kinetic parameters.

### Substrate-specificity assay

Cholic acid, lithocholic acid, and hydrocortisone (100 µM) (Tokyo Chemical Industry, China) substrate analogs of cholesterol were separately added to the reaction, and cholesterol (100 µM) as a control. The reaction mixture was incubated at 37°C for 12 h after the sterols were extracted; after that, the samples were subjected to LC-MS to determine the enzymatic activity of 5βChR. The flow injection analysis (FIA) was performed using an electrospray ionization (ESA) source working in positive mode for hydrocortisone and lithocholic acid while operating in negative mode for cholic acid. A gradient mobile phase containing pure acetonitrile was pumped into the mass spectrometer without a column. All injection volumes were kept at 20 µL.

### Circular dichroism spectroscopy of WT 5βChR and its mutants

CD spectroscopy of 5βChR (WT) or 5βChR-mutant strains was conducted using a J-815 CD spectropolarimeter (Jasco, Tokyo, Japan). The protein samples (20 µM) were prepared in KH_2_PO_4_ buffer (10 mM) (pH 7.8). A quartz cell 0.1 cm in length was used for the CD scan, and scans were carried out at 25°C from 190 to 260 nm at a 50 nm/min speed with a wavelength pitch and three accumulation data points. The data were analyzed using GraphPad Prism.

### Isothermal titration calorimetry assay for binding analysis

ITC analysis was performed using a MicroCal PEAQ-ITC (Malvern Panalytical, United Kingdom) with 300 cell volumes. The NADP^+^ cofactor was dissolved in dialysis buffer, and the substrate was dissolved in methanol. Purified 5βChR (20 µM) was injected with 18 successive 2 µl aliquots of NADP^+^ (1 mM) at 150 s intervals. Furthermore, protein-NADP^+^ was injected similarly to the substrate (500 µM). The data were fitted to a nonlinear regression model using a single binding site with MicroCal Origin software. The thermodynamic parameters were calculated using the Gibbs free energy equation (ΔG= ΔH-TΔS and ΔG= -RTln (K_a_)).

### 5βChR structure determination

5βChR structural modeling was performed using the ColabFold (v.1.5.5) implementation of Alphafold (https://github.com/sokrypton/ColabFold) [32, 33].

Multiple sequence alignment was performed using the MMseq2 algorithm [34]. The models were ranked through the pTM score, and the highest-scoring model was used for analysis. PyMOL [35] (v2.5.7 Schrödinger, LLC) was used for protein visualization and structural analysis of 5βChR. The DALI server [36] was used for structural similarity searches.

### Molecular docking of multiple ligands with 5βChR

The three-dimensional structure of 5βChR was predicted using ColabFold (v.1.5.5), which uses MMseqs2 along with AlphaFold2 in its background [32, 33]. The potential ligands of the 5βChR-NADP^+^-cholesterol dataset were downloaded from the ZINC drug database in SD format (https://zinc.docking.org/). AutoDockTools-1.5.7 was used to convert the 5βChR PDB files and ligand structures into PDBQT format [37]. Autodock Vina (v.1.2.3) performed molecular docking with multiple ligands [38]. The protein‒ligand interactions were computed and visualized with PyMOL [35] (v2.5.7 Schrödinger, LLC).

### Phylogenetic analysis of 5βChR within lactobacilli and plants

The prevalence of 5βChR was examined by creating a protein-based custom-made database for available lactobacilli from RefSeq using BLAST (v2.15.0) **(Table S2)**. 5βChR was used as a query search in the database, and the homologous proteins were retrieved for evolutionary studies. To determine the ancestral relationship of 5βChR in plants, 100 related sequences **(Table S3)** were removed from UniProt (https://www.uniprot.org/) [39]. MUSCLE (v3.8.31)[40] was used to align all the respective sequences, and phylogenetic analysis was performed using RAxML (v8) via the maximum likelihood Protcat blossom62 model [41].

### 5βChR sequence similarity network data analysis

For 5βChR sequence similarity network (SSN)[42] analysis, the amino acid sequence of 5βChR was used as a query in the UniProt database via BLASTp with a series of cutoff e-values of 10^−60^, 10^-70^, and 10^-8^ and protein identity values >50%, >60% and >70%, respectively. The EFI-EST tool was used to construct the protein SSNs of these sequences with various e-values, which were visualized by Cytoscape 3.3 [43].

### Determination of 5βChR abundance using healthy human metagenomic data

The abundance of 5βChR was checked in HMP[44, 45] as previously described using the ShortBRED **(Table S4)** [46]^-^[47]. This was performed using the EFI-chemically guided functional profiling (EFI-CGFP) platform [48–50]. Using UniProt, 5βChR sequence representatives were filtered at 80% amino acid similarity thresholds to identify nonredundant peptide markers. ShortBRED-quantify was used with default parameters to quantify the marker abundances in the 380 metagenomic database. The relative abundance was calculated based on reads per kilobase million (RPKM) values and converted into ‘copies per microbial genome’, as explained earlier [50, 51].

### Abundance analysis of the 5βChR gene in human gut metagenomes under disease conditions

The shotgun metagenomic sequencing datasets of the human gut microbiomes describing diabetes were downloaded from the Sequence Read Archive of the NCBI (https://www.ncbi.nlm.nih.gov/sra). The dataset was divided into healthy and diseased based on the metadata provided with the original data. The read quality was checked using FastQC (v0.11.9) (https://www.bioinformatics.babraham.ac.uk/projects/fastqc/), followed by MultiQC (v1.11) [52]. Only reads with a phread score higher than 30 were included in the study. The high-quality sequencing reads were assembled using MEGAHIT (v1.2.9) with a minimum contig length of 500 bp [53]. The assembled sequences were matched to the nucleotide sequence of 5βChR using DIAMOND (v2.0.15.1531) with the following parameters: e value – 0.00001, query coverage 30, and percent identity 30 [54].

## Authors contribution

U.N. designed the study, curated the data, developed the methodology, conceptualized the project, and wrote the original draft. V.B. methodology and writing.

A.G. Methodology, Data curation, Visualization, and writing. RS methodology. A.G. methodology. V.S.B. methodology and data curation. KA. Formal analysis. HPS. LC-MS methodology and data analysis. SM. LC-MS methodology and data analysis. SL. Methodology. MS. Methodology. K.T. Formal analysis and writing. NKN Formal analysis, conceptualization, writing, resources, and project administration.

## Acknowledgments

This work was supported by the National Priority Project (NPP) under the Corpus Fund Head (CFH) of the Indian Council of Agricultural Research. Urmila Netter thanks the Council of Scientific and Industrial Research (CSIR), India, for providing a research fellowship. For financial support, Vishakha Bisht and Amit Gaurav thank the Ministry of Human Resource and Development and the Department of Biotechnology (DBT) Govt. of India.

## Supplementary information

### 1. Supplementary Figures

**Fig. S1.**
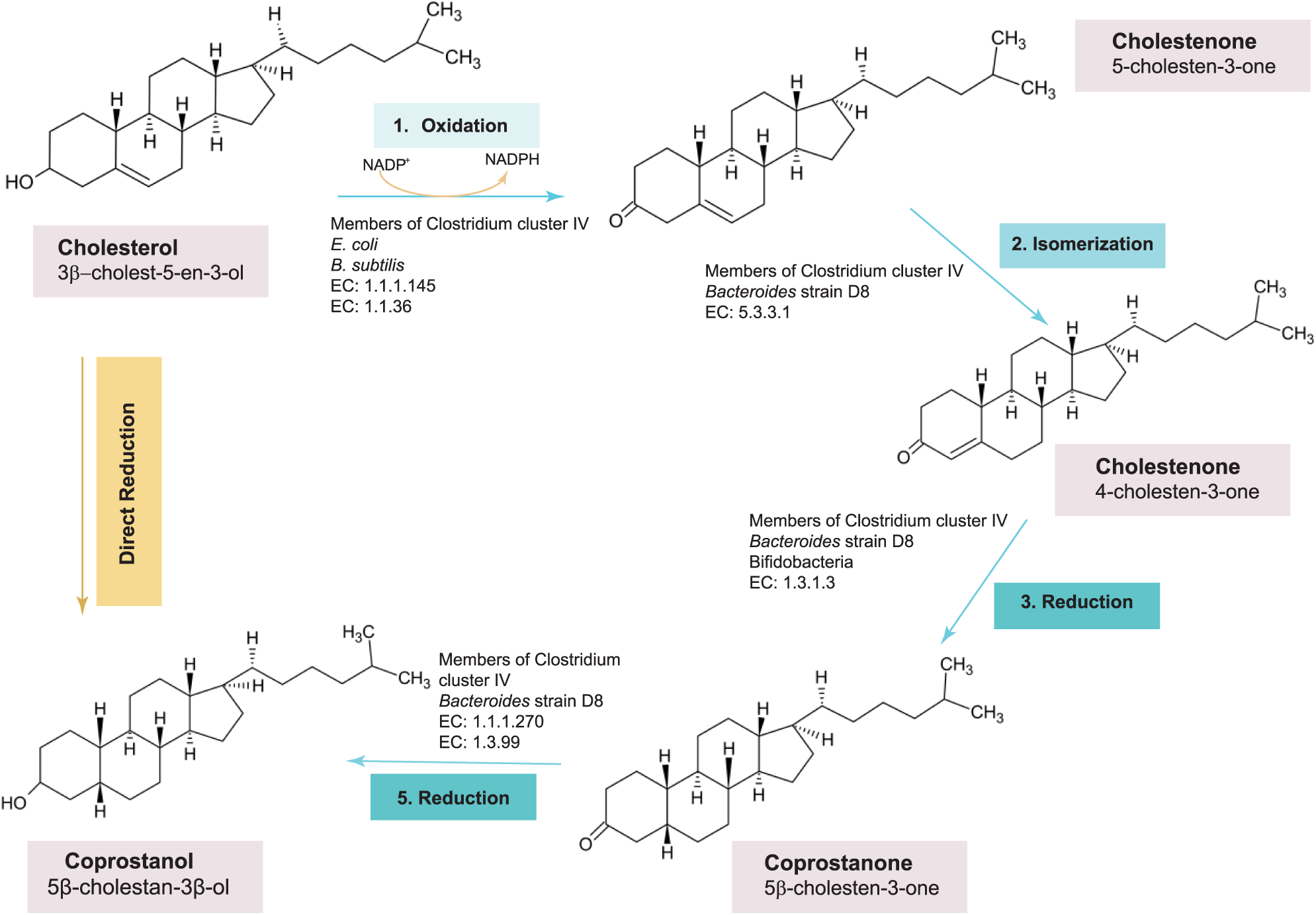
Biotransformation pathways of cholesterol to coprostanol by gut microbes.

**Fig. S2.**
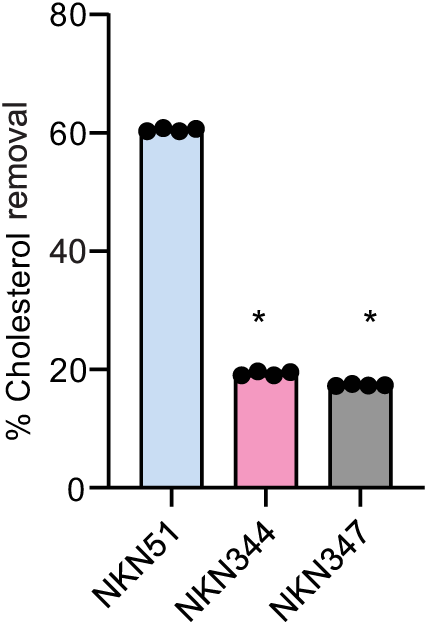
Rudel & Morris cholesterol depletion assay representing in-vitro cholesterol removal by different *Lactobacillus* strains. Each point represents the biological replicate and *p-value* calculated by the Mann-Whitney test (\**p*-0.0286).

**Fig. S3.**
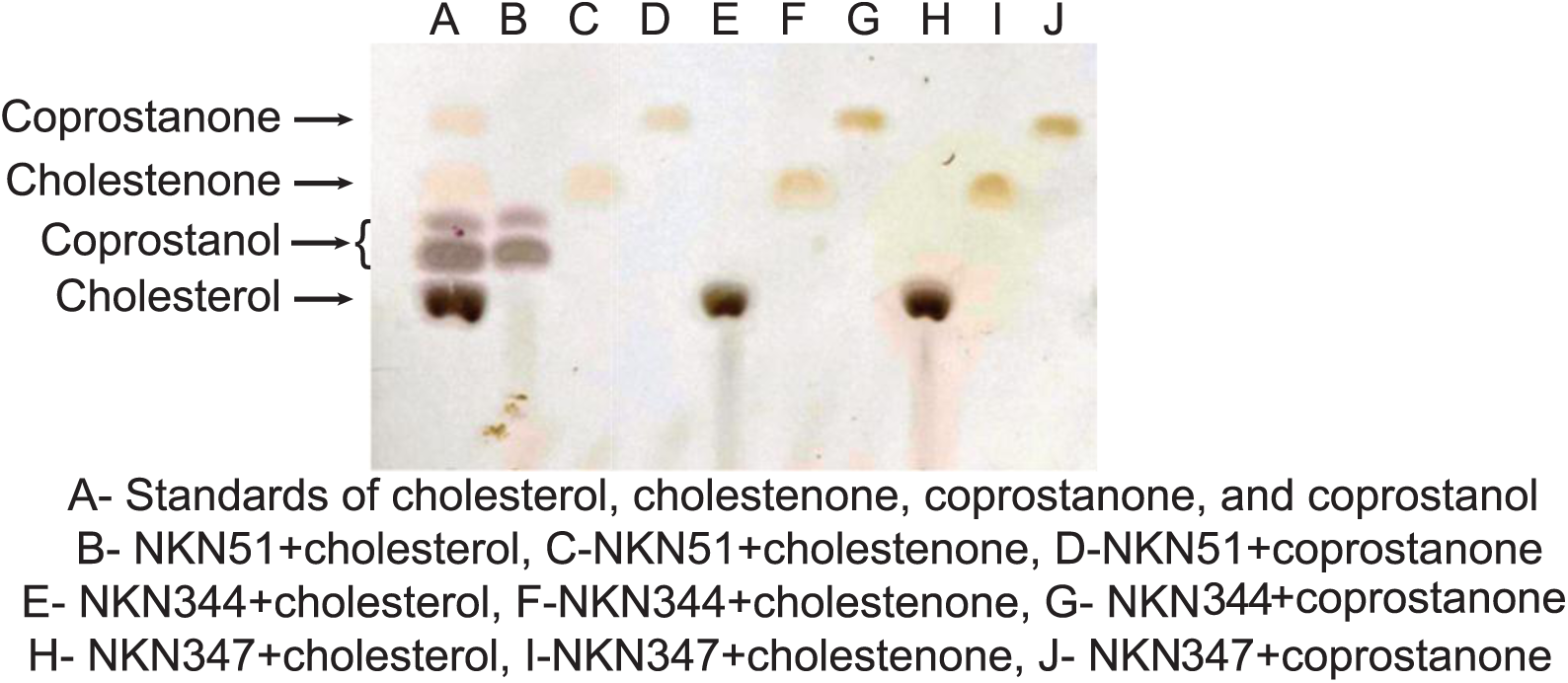
Thin-layer chromatography (TLC) screening of cholesterol transformation to coprostanol by lactobacilli. The whole-cell lysate of different lactobacilli strains supplemented with NADP^+^ (100 µM) and either of substrates, cholesterol (100 µM), cholestenone (100 µM), or coprostanone (100 µM) for 12 h in anaerobic conditions.

**Fig. S4.**
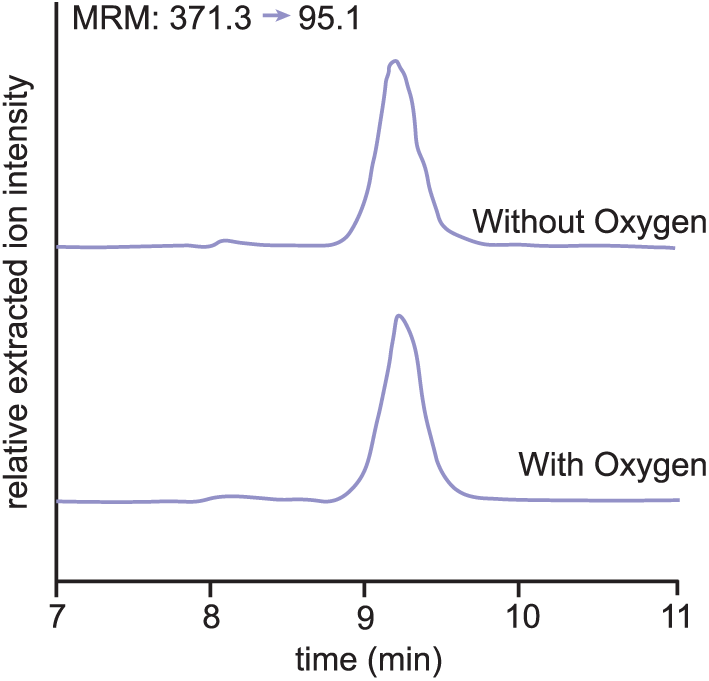
Analysis of *L. fermentum* NKN51 lysate activity in microaerophilic conditions by Liquid chromatography Mass Spectrometry (LC-MS). The provided extracted ion chromatogram (EIC) represents the product of the whole-cell lysate assay of *L. fermentum* NKN51 supplemented with NADP^+^ (100 µM) and cholesterol (100 µM) over 12 h in microaerophilic and anaerobic conditions as a control.

**Fig. S5.**
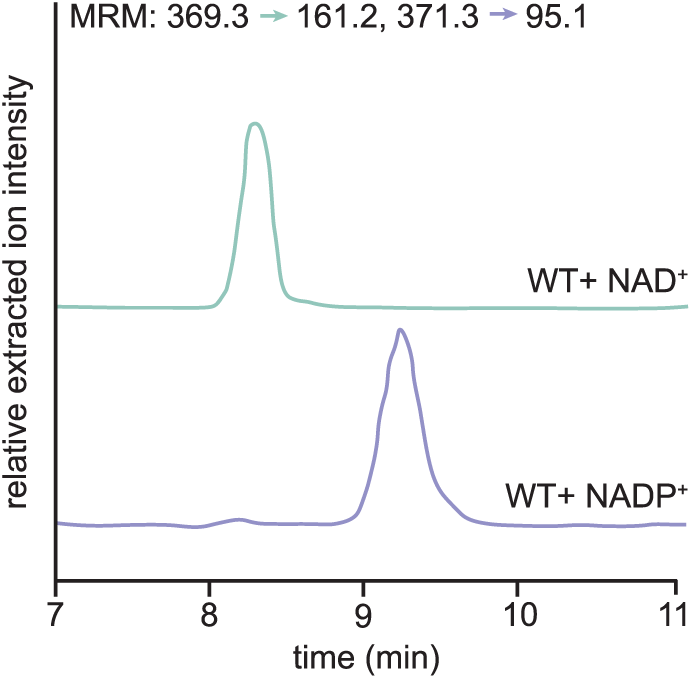
Investigating the role of NAD^+^ as a cofactor for cholesterol metabolism by LC-MS. The reaction mixture of *L. fermentum* NKN51 whole-cell lysate with cholesterol (100 µM) and NAD^+^ (100 µM) or NADP^+^ (100 µM) for 12 h. No reaction was observed with NAD^+^ as a cofactor.

**Fig. S6.**
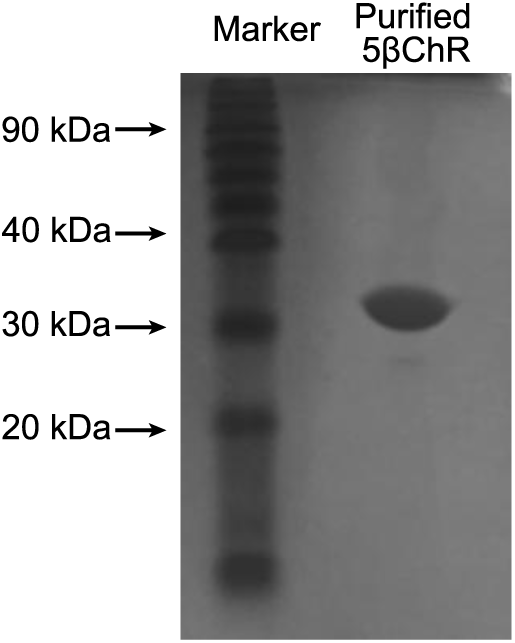
Recombinant putative cholesterol dehydrogenase protein purification by immobilized affinity chromatography (IMAC). The SDS-PAGE depicts purified 5βChR protein by Ni-NTA-based affinity chromatography, with a theoretical molecular weight of 29.5 kDa.

**Fig. S7.**
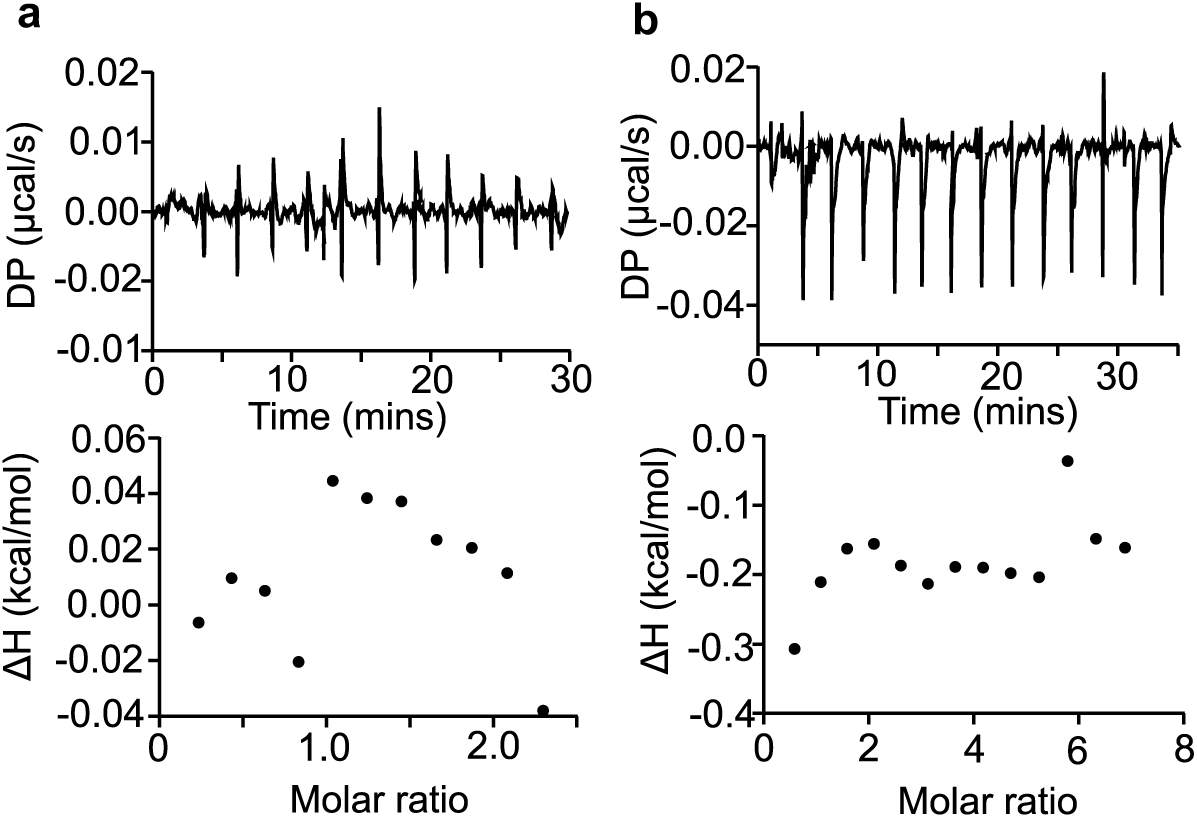
Binding studies of 5βChR protein with substrates. The Isothermal Titration Calorimetry (ITC) of WT5βChR with (a) cholestenone and (b) coprostanone, wherein the points represent the normalized enthalpy change of the reaction at the different time intervals.

**Fig. S8.**
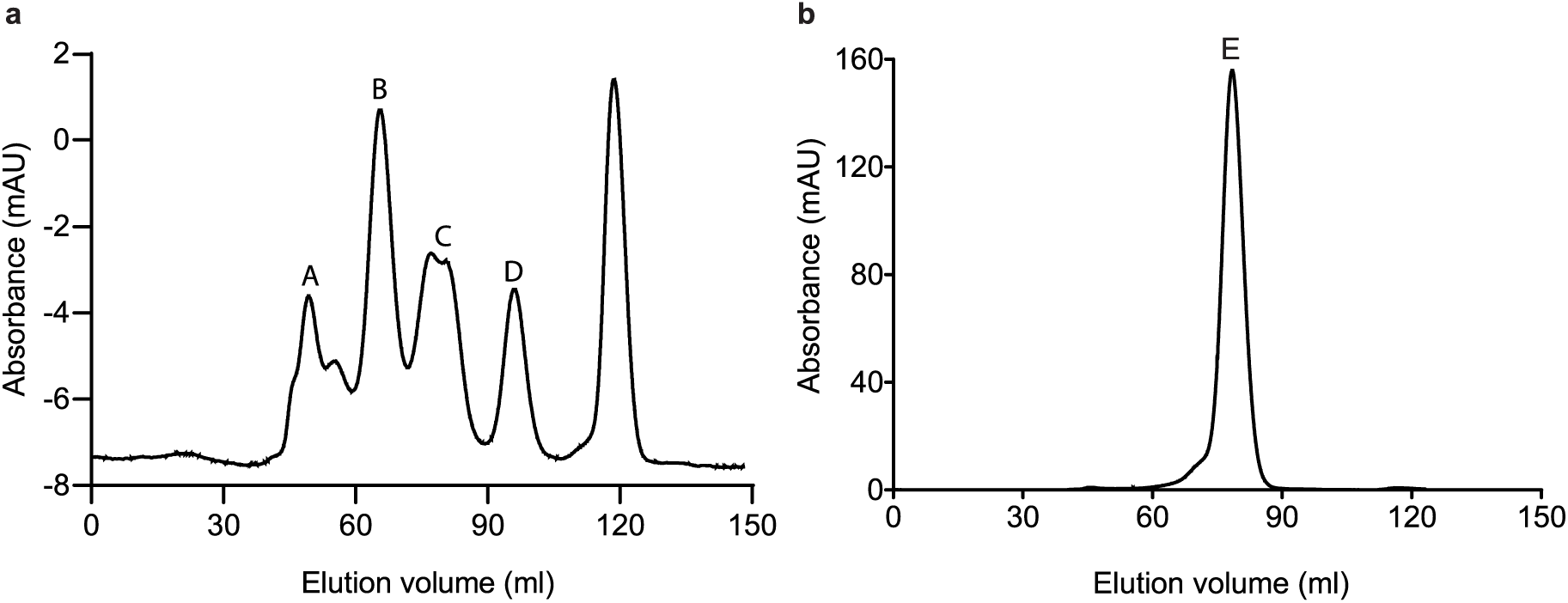
Calculation of native molecular size of recombinant 5βChR. **(a).** The gel filtration graph of the standard mixture consists of various proteins represented by a distinct peak and their respective molecular weight along with elution volumes, which are as follows: A-Thyroglobulin (670kDa) 49.33ml, B-Gamma globulin (150 kDa) 65.63ml, C-Ovalbumin (44.5 kDa) 80.02 ml and, D-Ribonuclease A (13.7 kDa) 96.97 ml. **(b).** A 29.5 kDa recombinant cholesterol dehydrogenase (E) eluted at 78.43 ml, depicts its dimeric form.

**Fig. S9.**
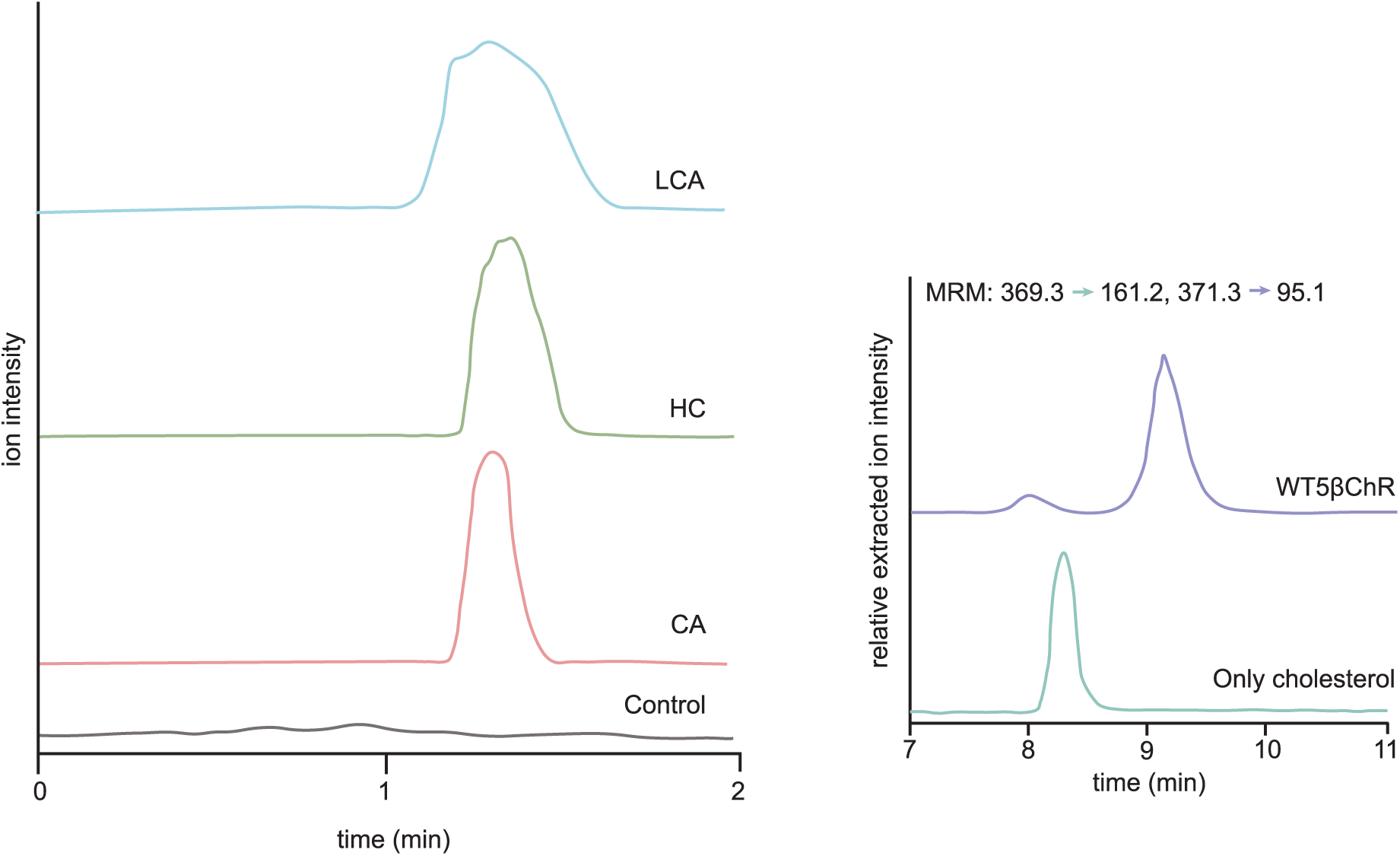
5βChR enzyme activity with a similar structure compound like cholesterol. The total ion chromatograph demonstrates no reaction observed by 5βChR incubated with either 100 µM cholic acid (CA), hydrocortisone (HC), or lithocholic acid (LCA). EIC shows the coprostanol detection from the reaction of 5βChR incubated with cholesterol (100 µM) and NADP^+^ (100 µM).

**Fig. S10.**
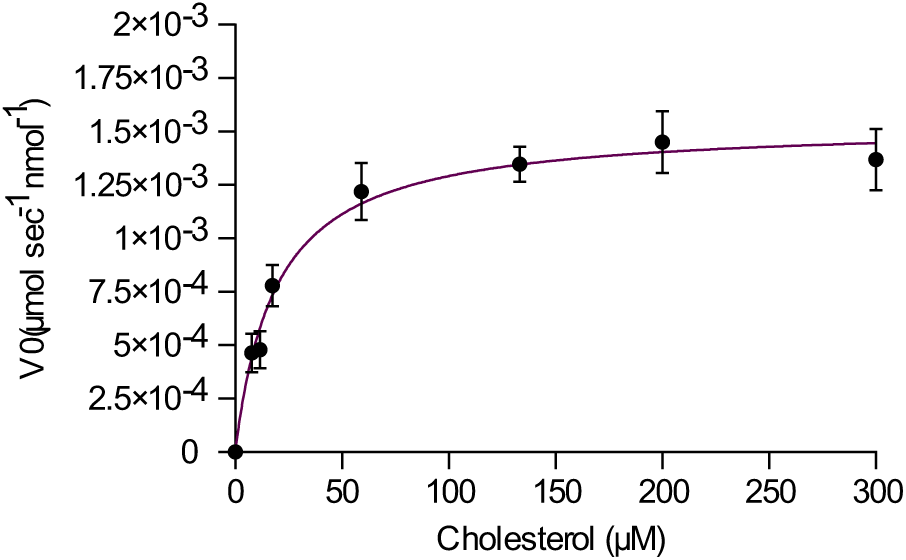
Enzyme kinetics of WT5βChR. The Michalis-Menton curve is determined from the reaction of cholesterol with 5βChR. The saturation curve predicts the kinetics of cholesterol reduction by recombinant 5βChR enzyme with *Km*-19.09 µM and *Vmax*-15.35x10^-6^ µmol sec^-1^ nmol^-1^.

**Fig. S11.**
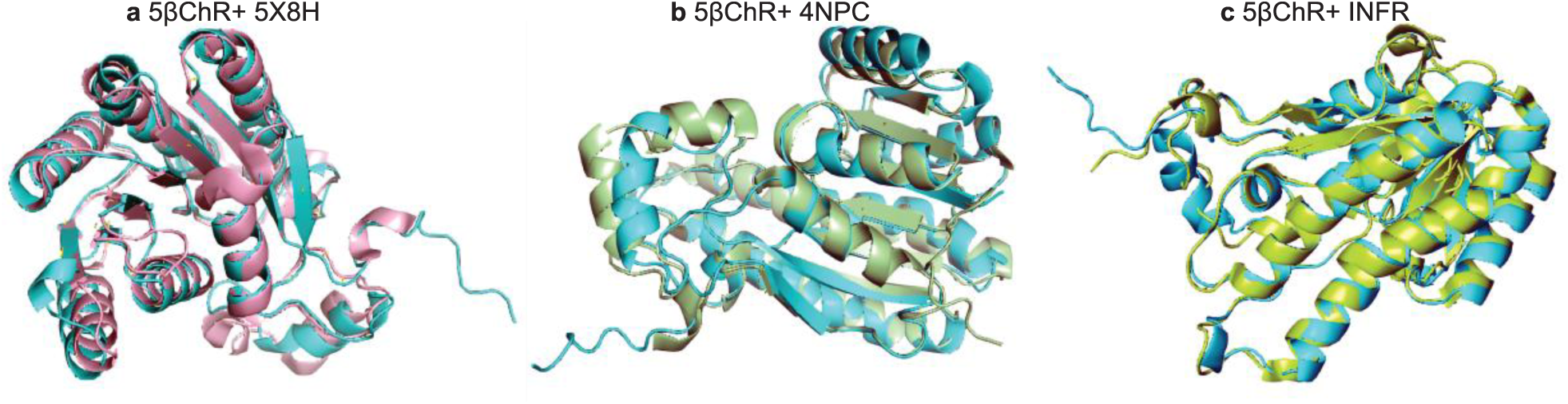
Structural comparison of 5βChR with its predicted homologs. According to the DALI server, the closest structural homologs of 5βChR are SDR reductase (PDB ID: 5X8H), Sorbitol dehydrogenase (PDB ID: 4NPC), and Putative oxidoreductase RV2002 (PDB ID: 1NFR). These three enzymes are superimposed onto 5βChR with root square mean value (r.m.s) deviation of 1.3, 1.3, and 1.2 Å, respectively.

**Fig. S12.**
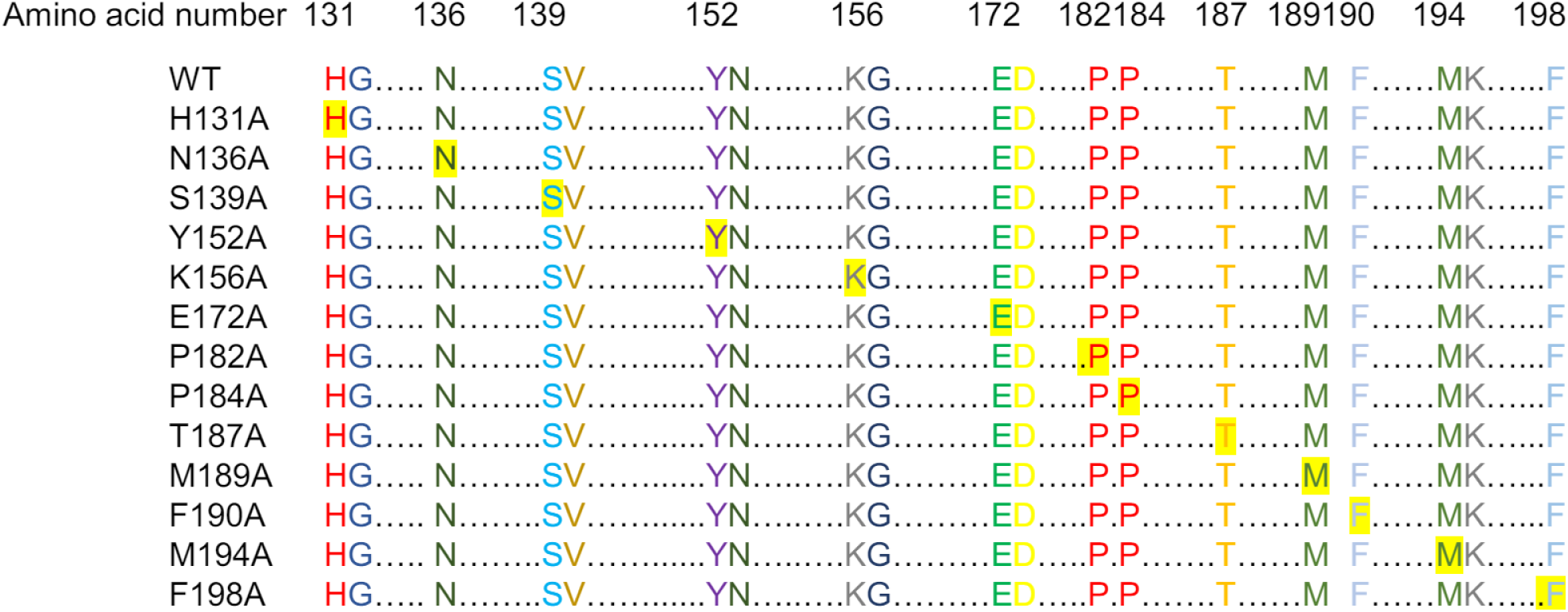
Site-directed mutagenesis of WT5βChR. The highlighted amino acids represent the mutation in conserved residues of WT5βChR. N-terminus residues, H131, N136, S139, Y152, K156 and E172 mutated to alanine. C-terminus residues, P182, P184, T187, M189, F190, M194, and F198, are converted to alanine for cholesterol binding analysis.

**Fig. S13.**
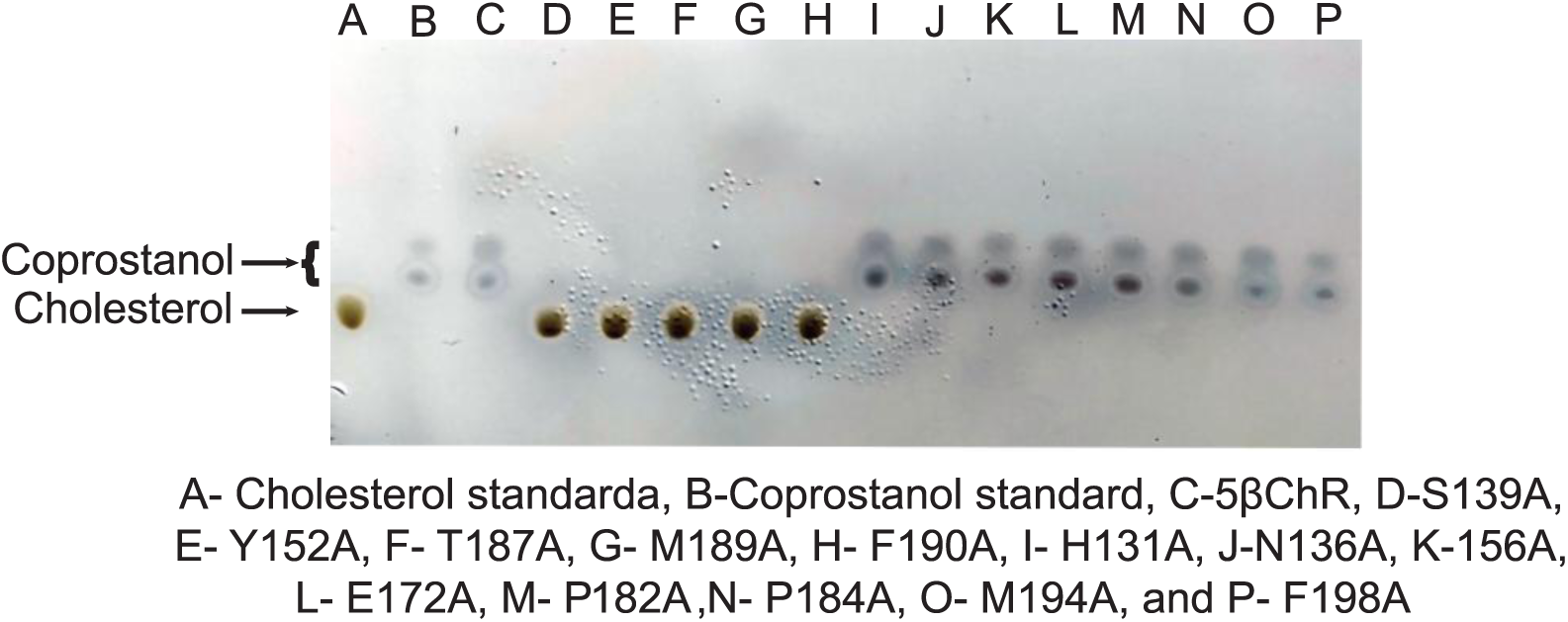
Cholesterol transformation profile of WT and mutant 5βChR. The reaction mixture of 5βChR WT and mutants with cholesterol (100 µM) and NADP^+^ (100 µM) are analyzed through TLC. S139A, Y152A, T187A, M189A, and F190A show the spot equivalent to cholesterol. H131A, N136A, 156A, E172A, P182A, P184A, M194A, and F198A highlighting spots corresponding to coprostanol.

**Fig. S14.**
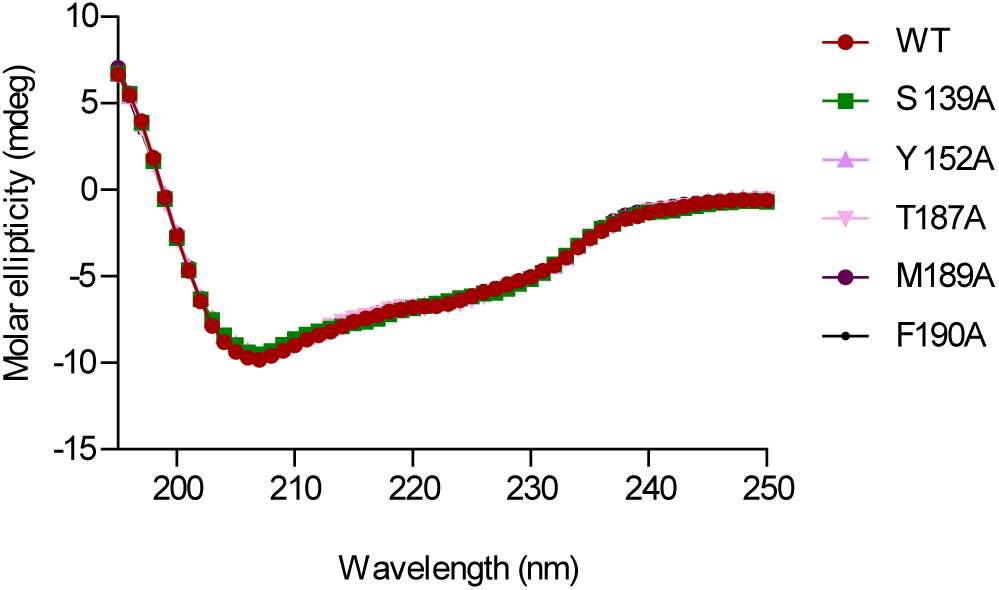
Circular Dichroism spectra of 5βChR and its mutants. The graph represents the secondary structure profiles of WT5βChR, S139A, Y152A, T187A, M189A, and F190A.

**Fig. S15.**
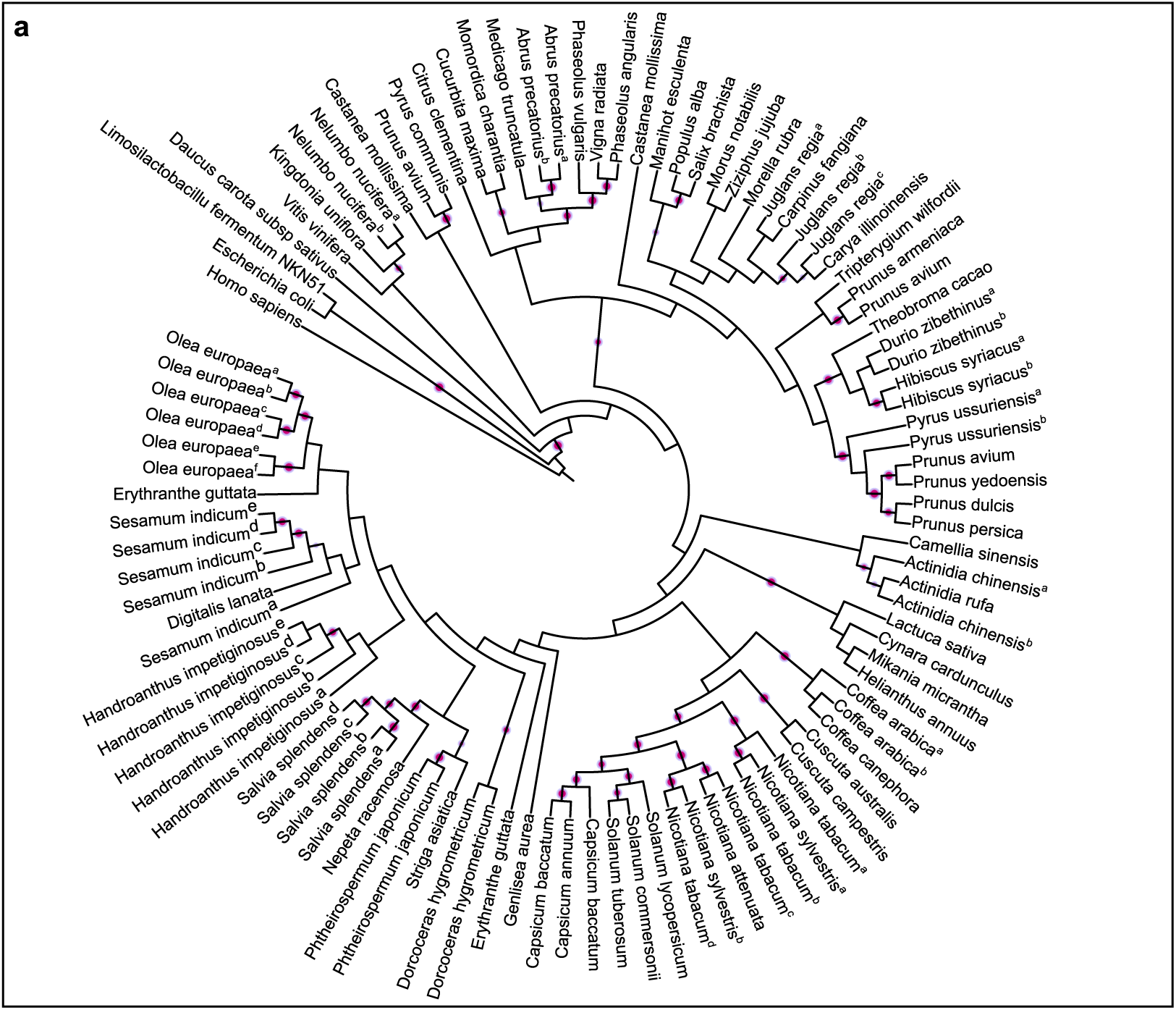
Mapping the prevalence of 5βChR within plants by phylogenetic profiling. **(a).** The evolution analysis of 5βChR homologs in plants (∼100) is acquired from UniProt in RAxML using the PROTCATWAG model with human 5β reductase as an outgroup. 5βChR and plant 5β reductase evolved from a common ancestor. A pink circle indicates a bootstrap value of more than 70.

**Fig. S16.**
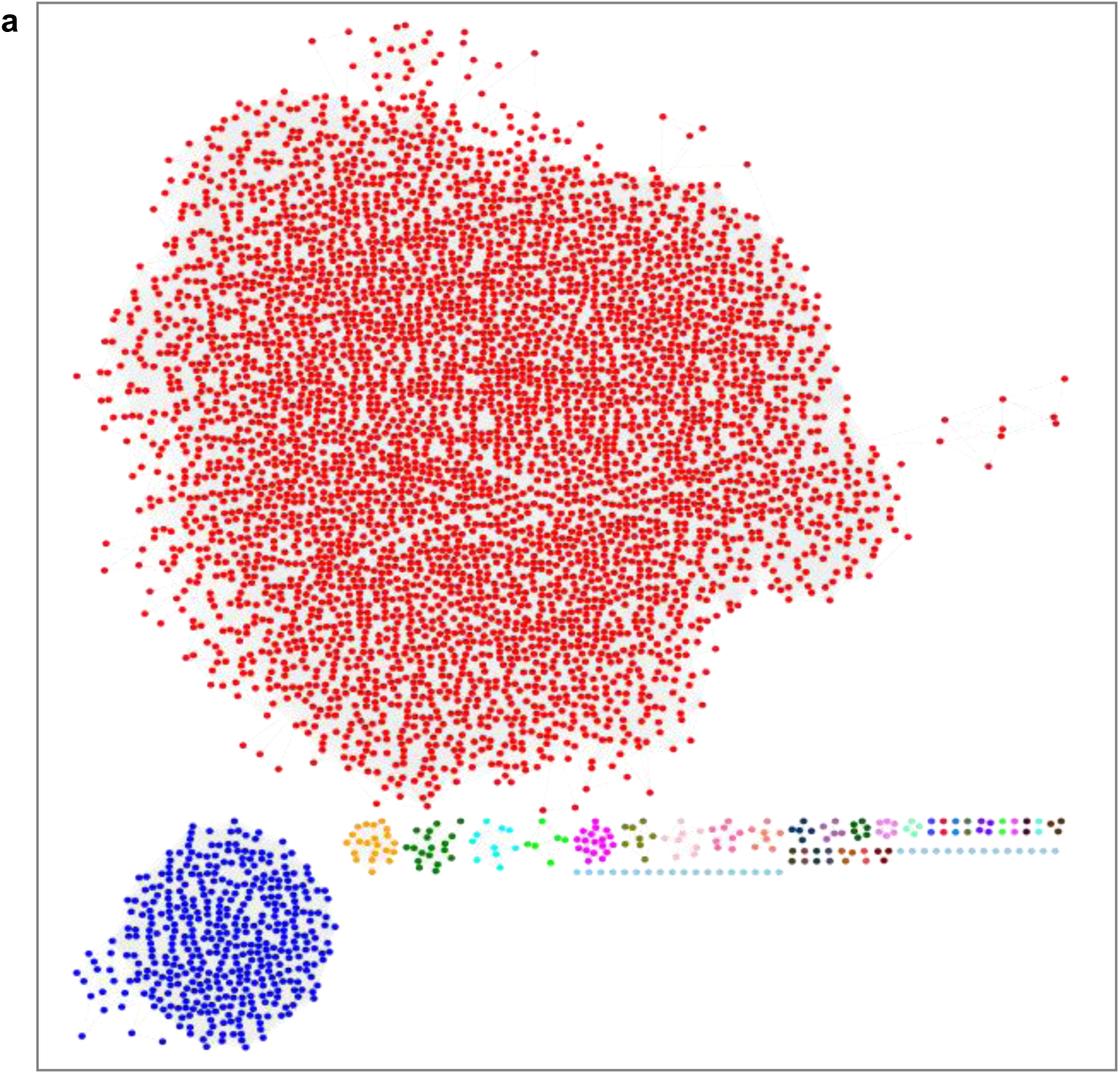
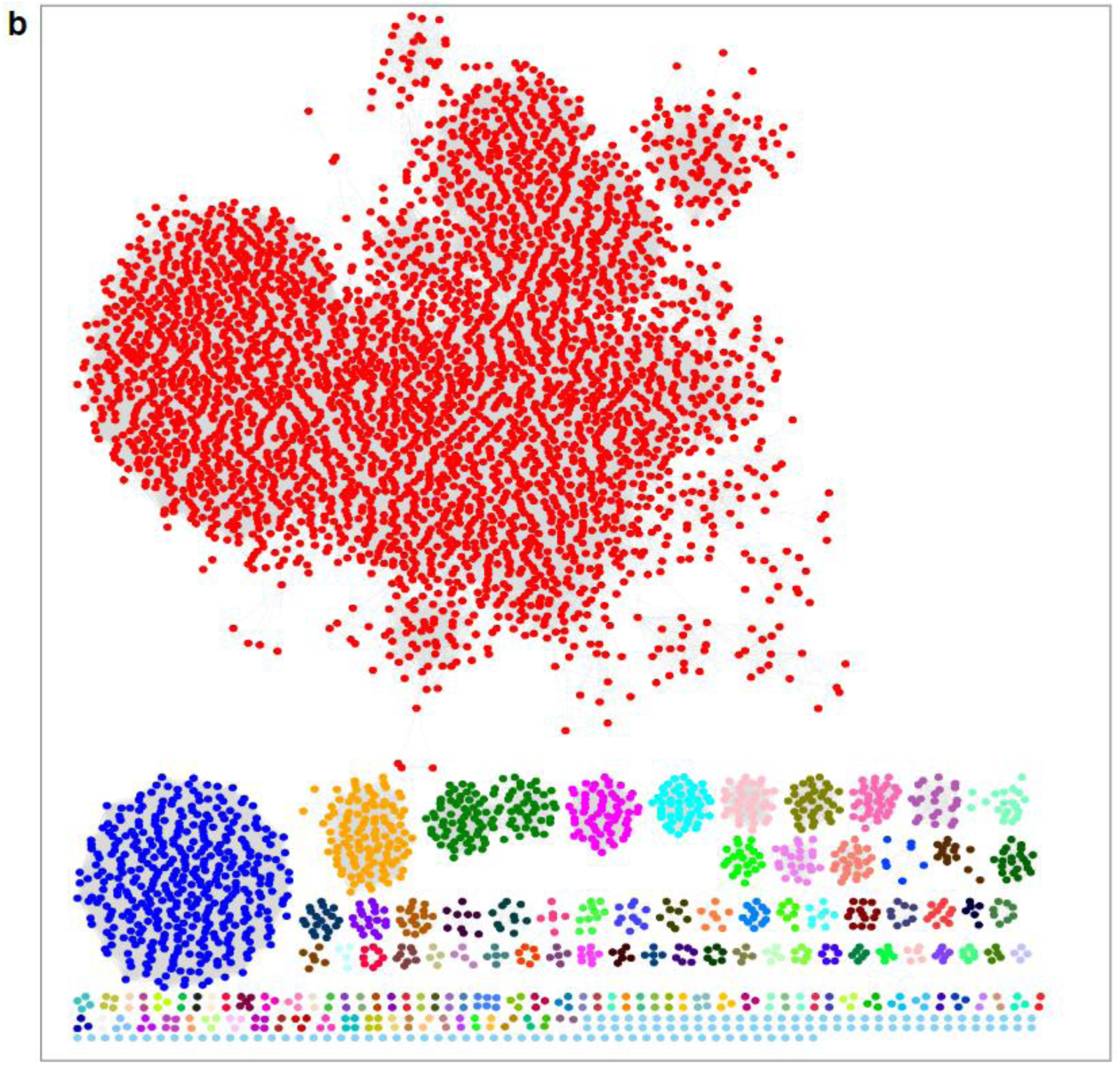
5βChR sequence similarity network. **(a)** A sequence similarity Network (SSN) is built using 5000 proteins from the UniProt with 60 sequence coverage and >50 sequence identity to 5βChR. 4383 proteins out of 5000 are combined into a single cluster, and the rest of the protein sequences are divided into 32 clusters. **(b)** With a 70 threshold and >60 sequence identity to 5βChR, the 5000 proteins are divided into 138 clusters based on functional similarity and increasing sequence identity.

**Fig. S17.**
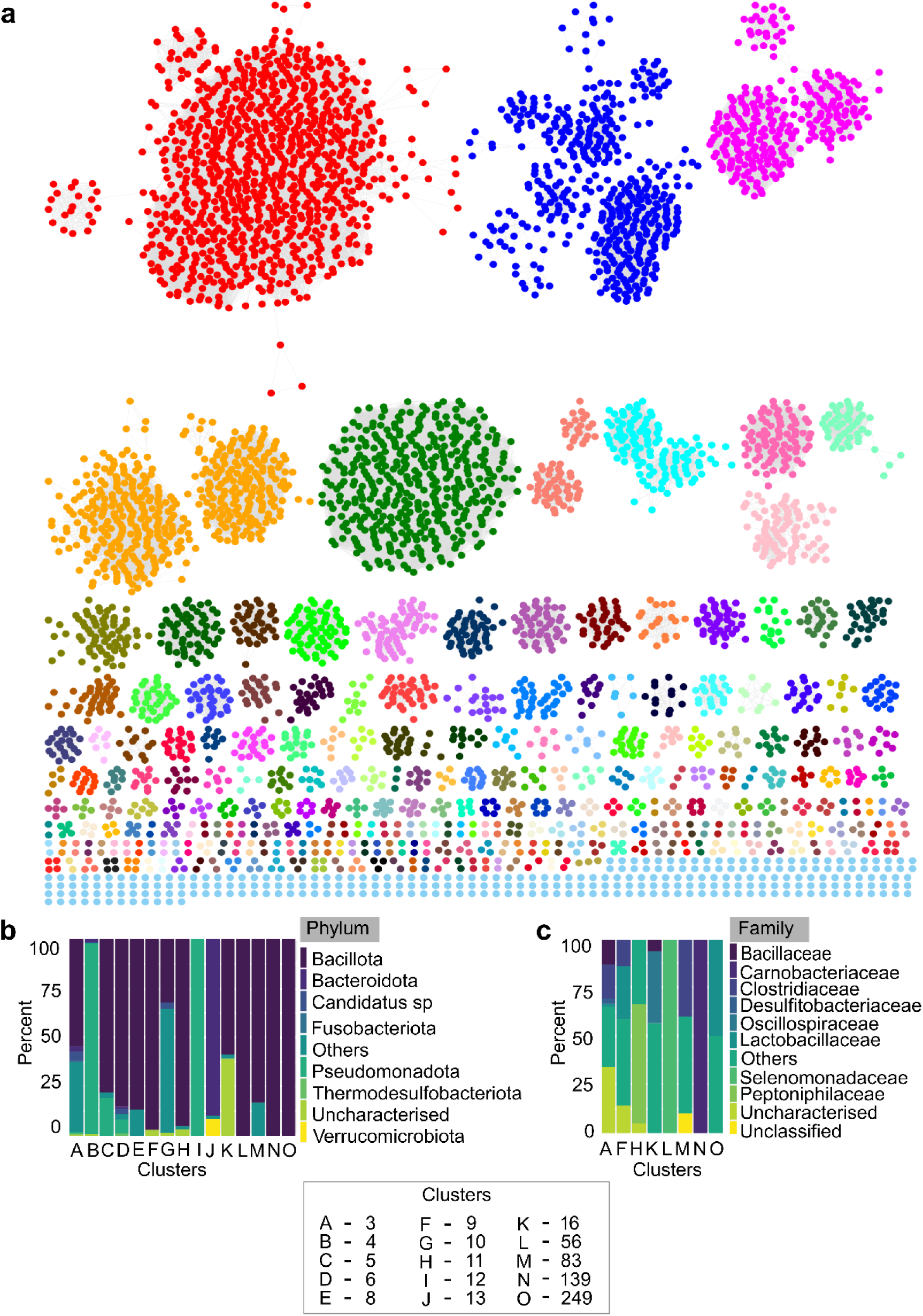
Network maps illustrating the abundance and distribution of the 5βChR gene family across the different microbes. **(a)**. Microbial origin 5βChR protein homologs are retrieved from UniProt for sequence similarity network (SSN) analysis. These networks are visualized by the Enzyme Function Initiative-Enzyme Similarity Tool (EFI-EST) with threshold values of 10^-80^. The proteins are divided into 269 clusters based on sequence identity, structural similarities, and functional profile of 5βChR proteins, highlighting their presence in various microbes. **(b-c)** The graphs depict the taxonomy of microbial 5βChR homologs from the selected clusters, which are prevalent in human microbiome data. The representative sequences of clusters belong to different bacteria belonging to diverse phyla and families.

**Fig. S18.**
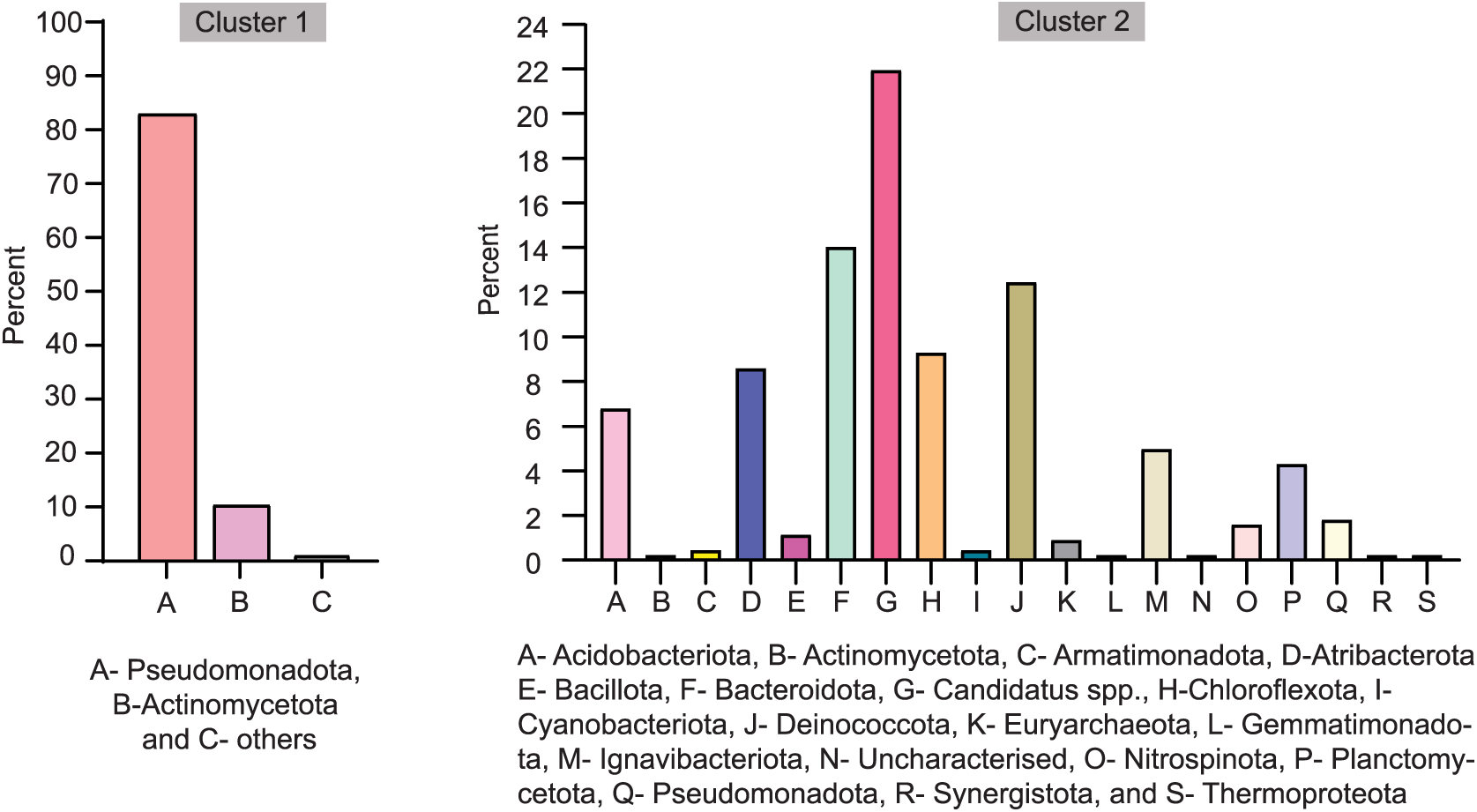
Assessing the taxonomy of 5βChR representatives. The phyla-based classification of clusters 1 and 2 of 5βChR homologs with >80 sequence coverage and >70 sequence identity. The bar graphs show the prevalence of 5βChR protein in various phyla.

**Fig. S19.**
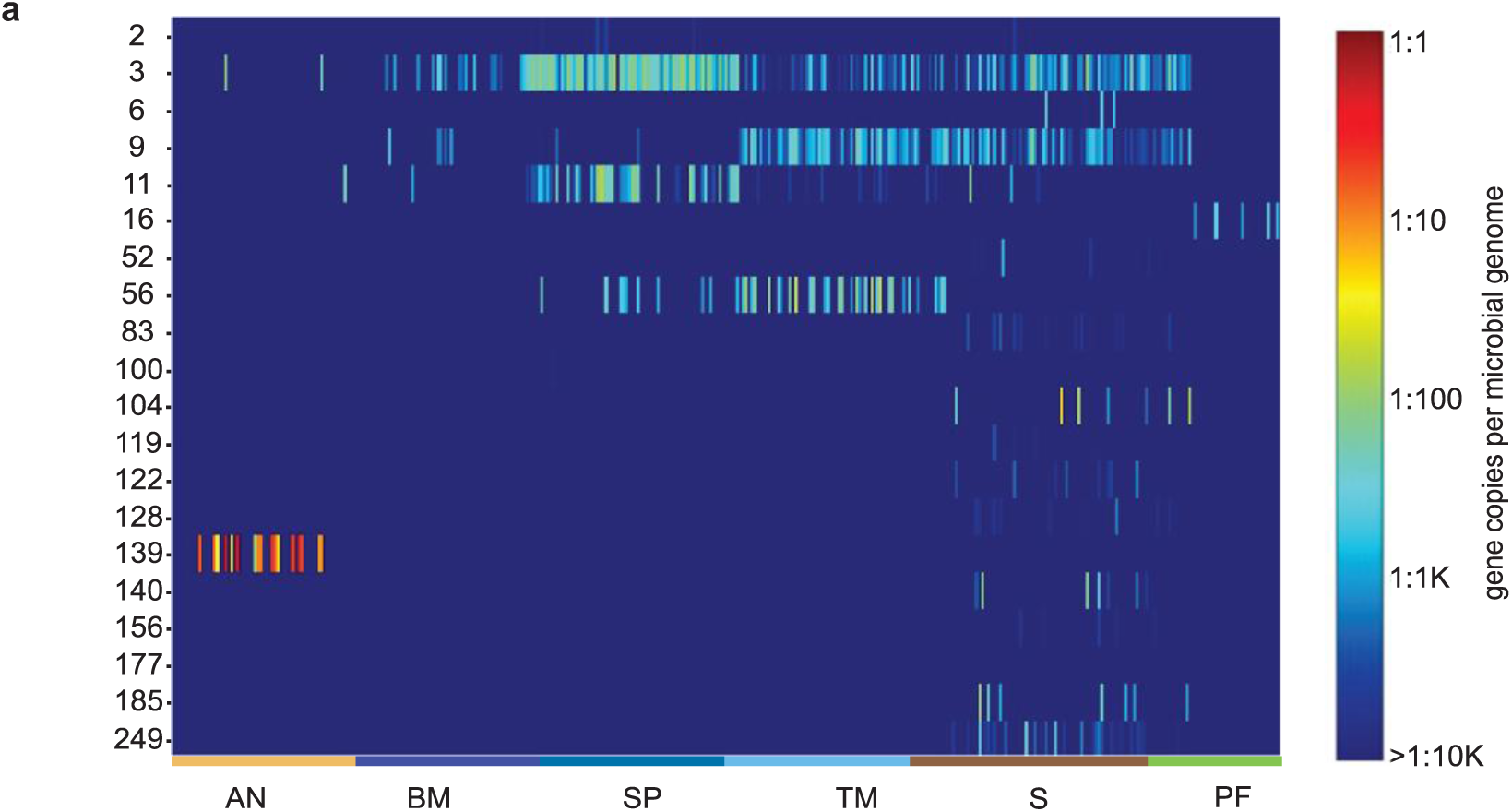
Meta-analysis depicts the abundance of 5βChR homologs in a healthy microbiome. The heat map illustrates the prevalence of 5βChR homologs in the human metagenomic data of 380 healthy individuals. Analysis validates the presence of 20 clusters in the participants’ metagenome data.

**Fig. S20.**
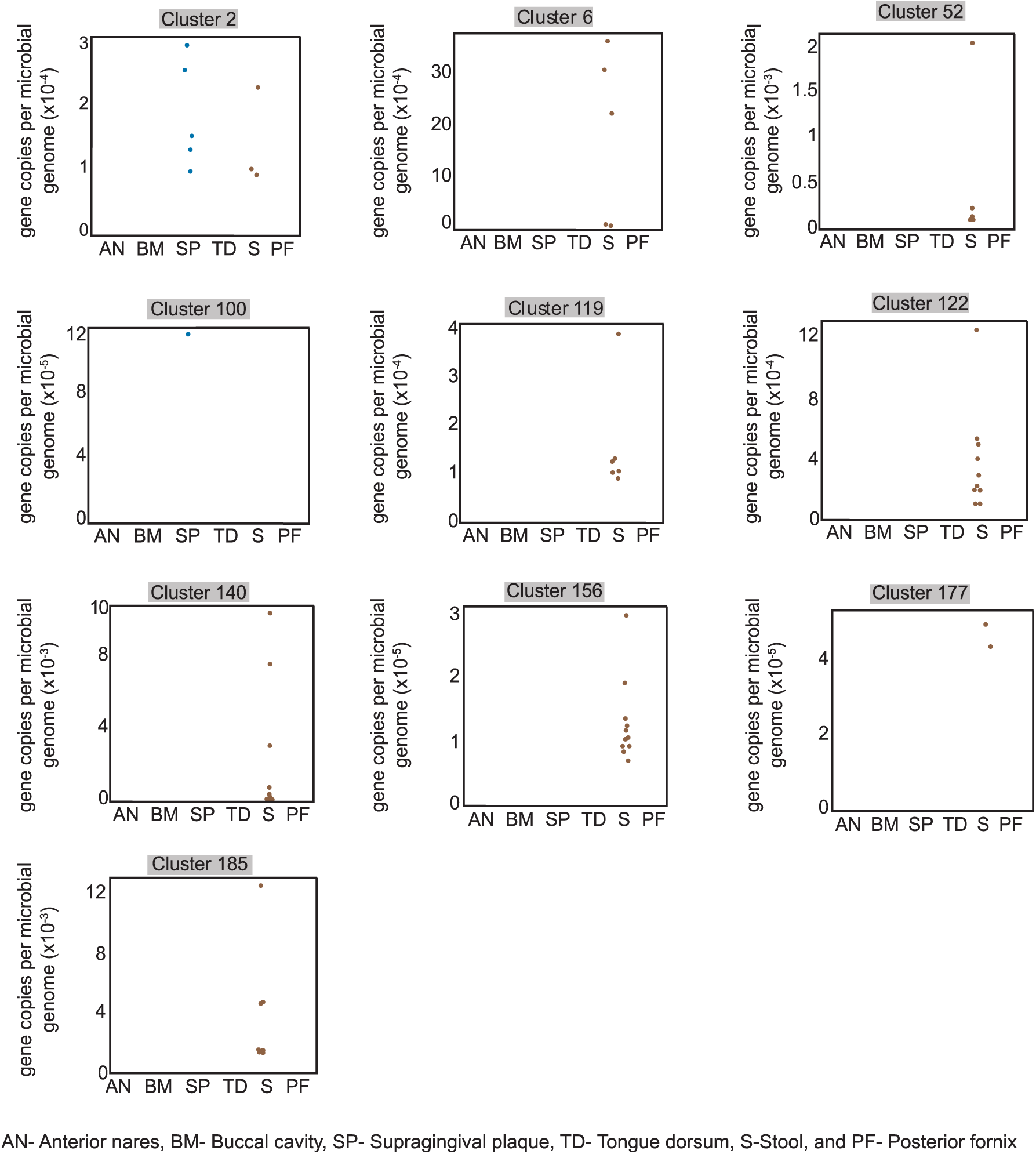
The distribution of different 5βChR homolog clusters in the microbiome data of healthy human participants. The box plot displays the various clusters of 5βChR homologs primarily present in stool samples.

### 2. Supplementary Tables

**Table S1.**
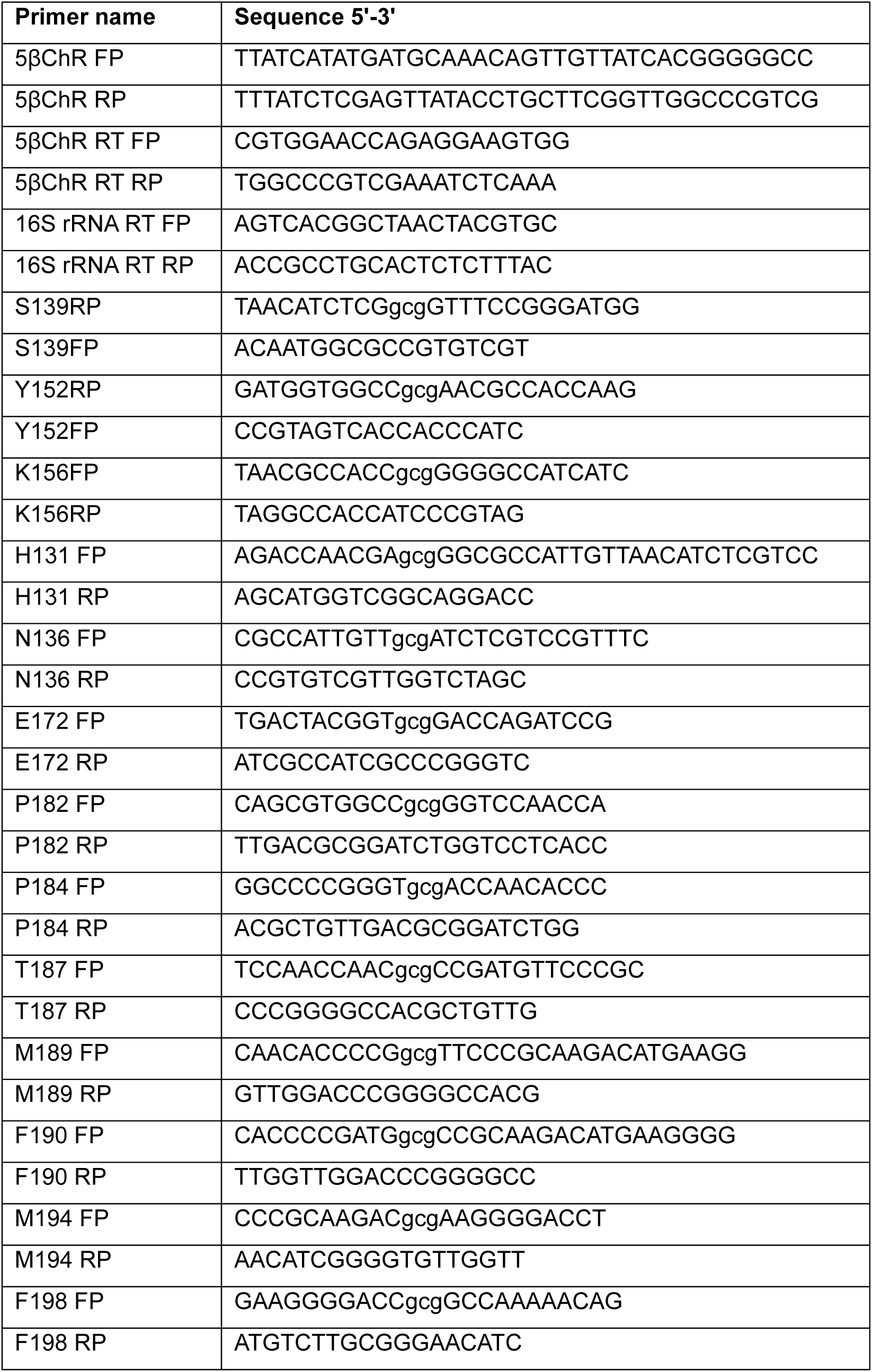
Primer details used in this study.

**Table S2.**
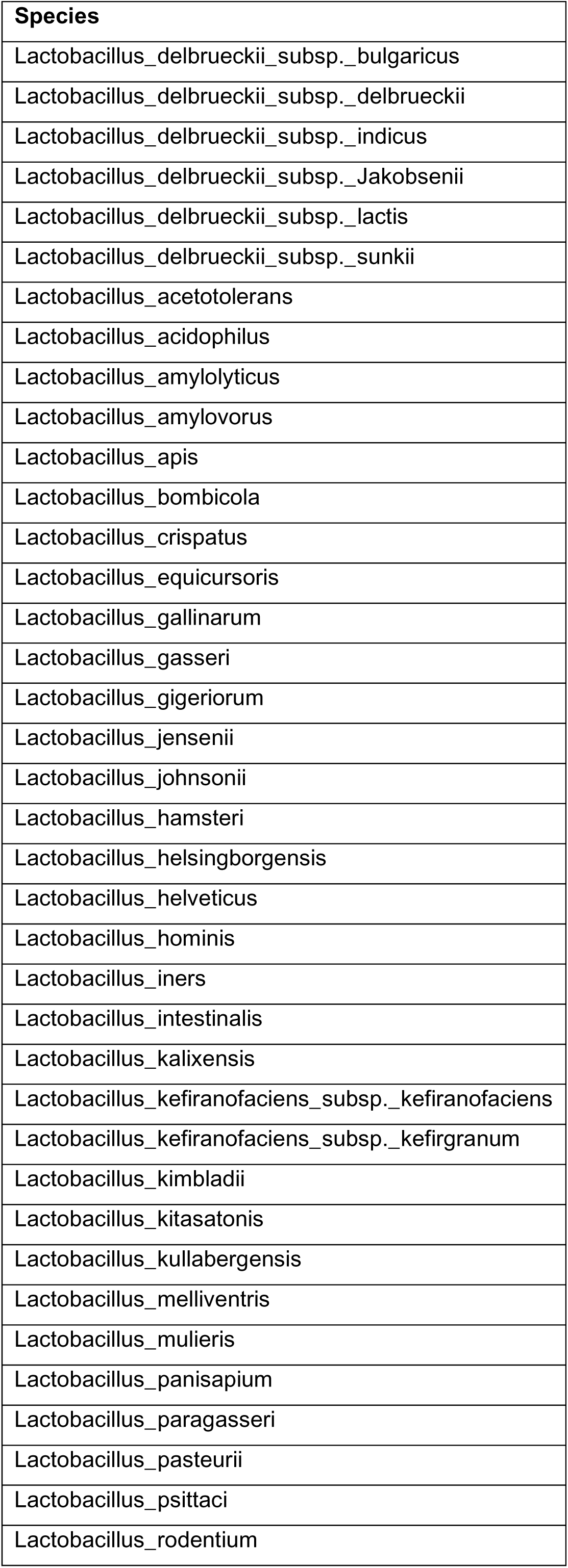
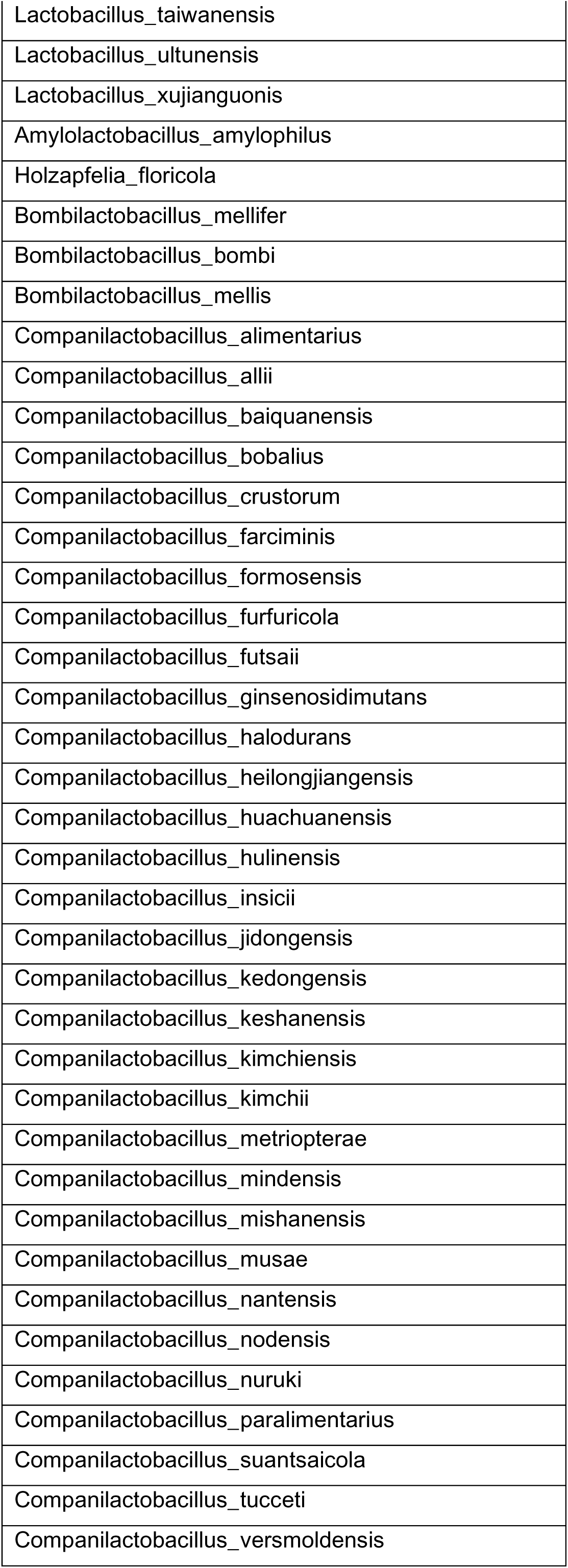
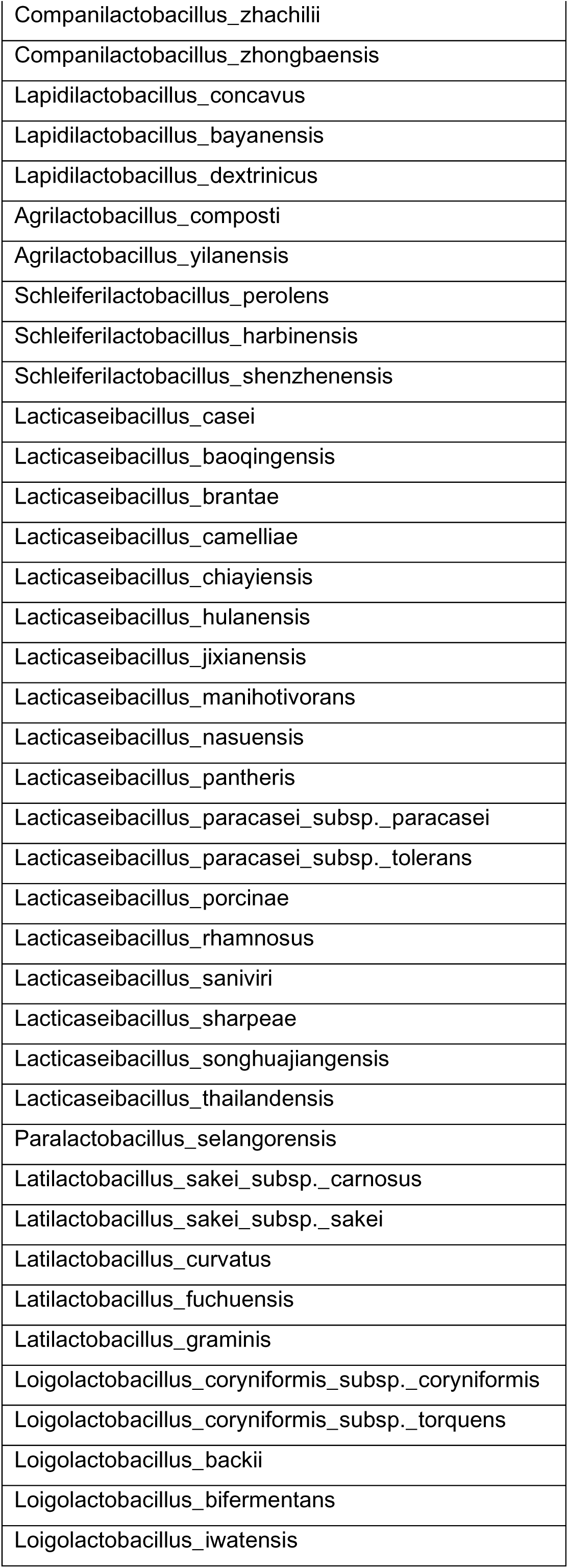
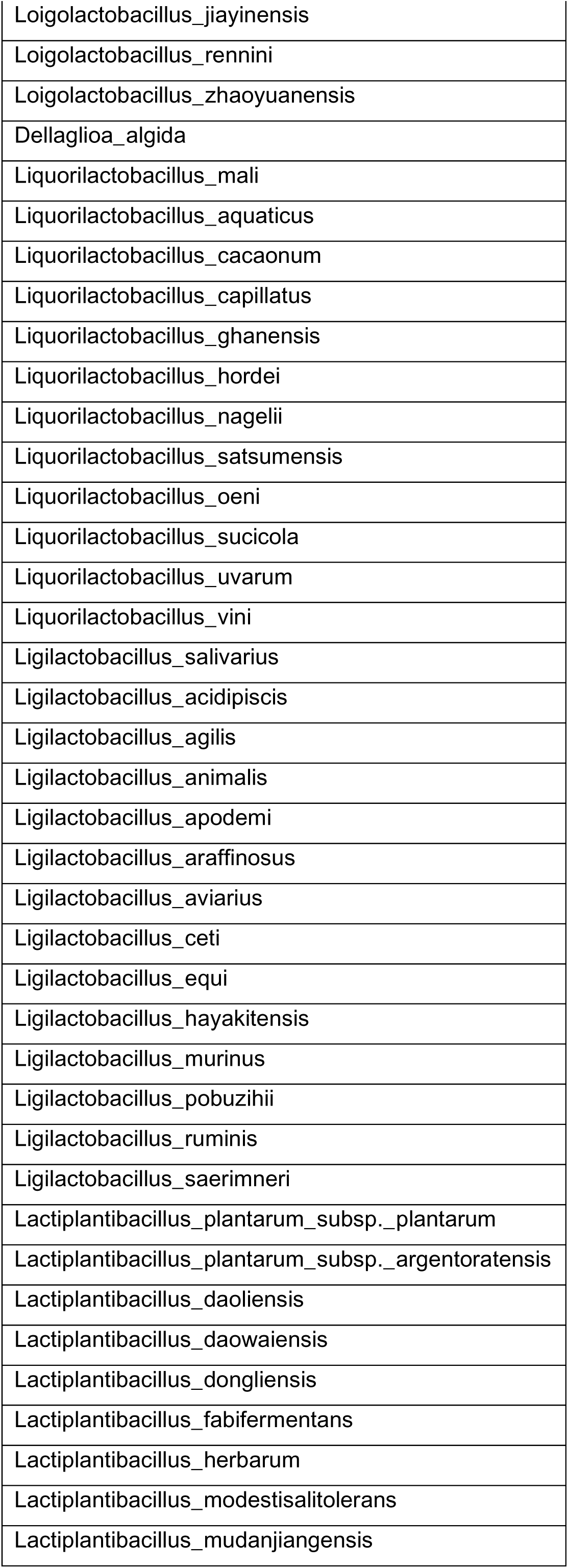
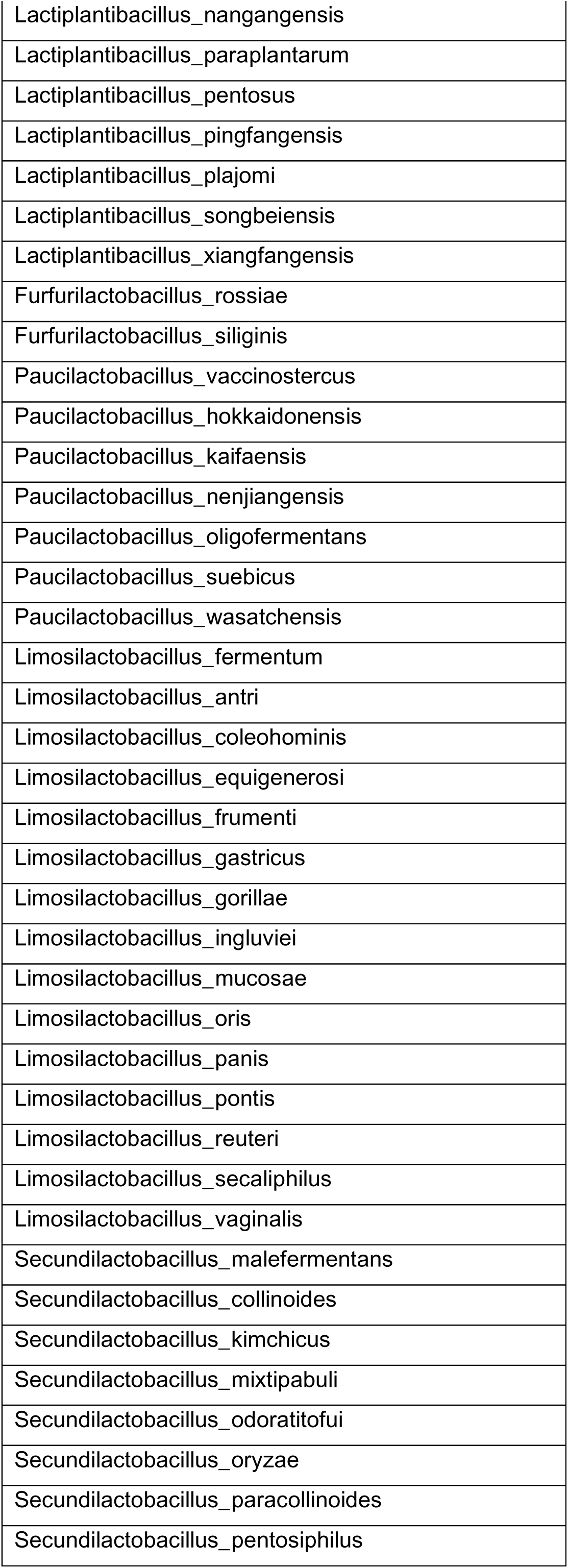
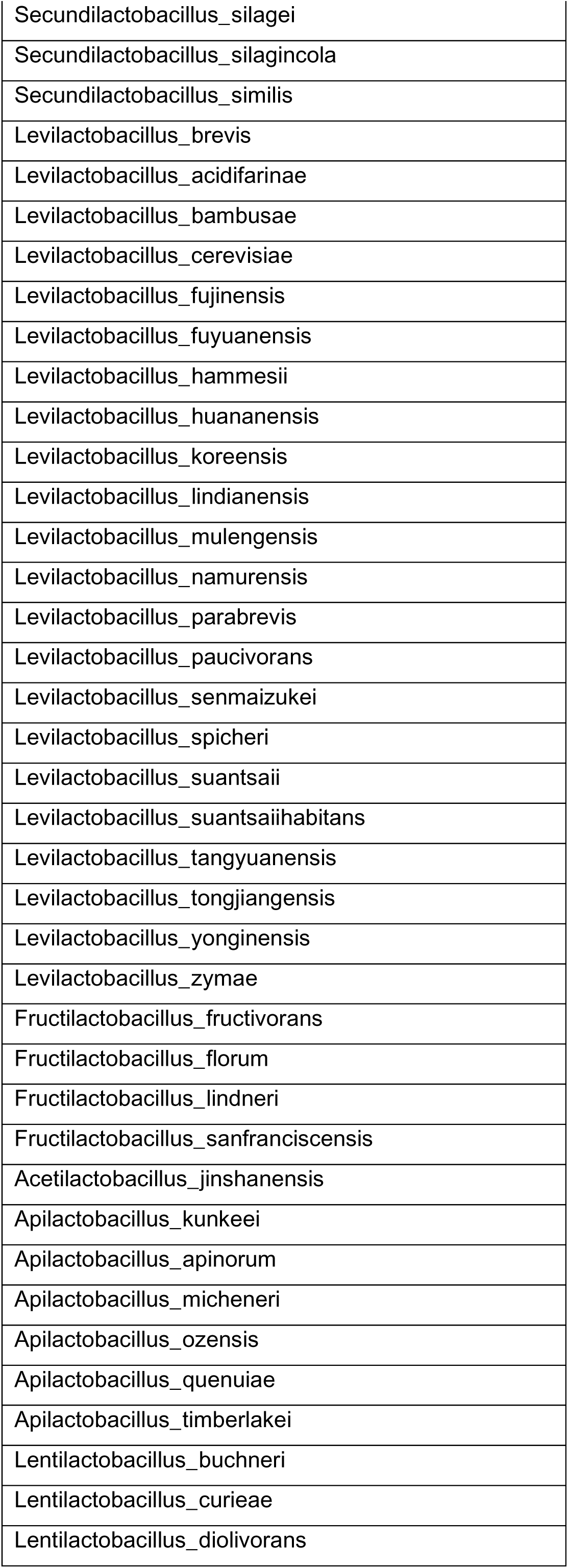
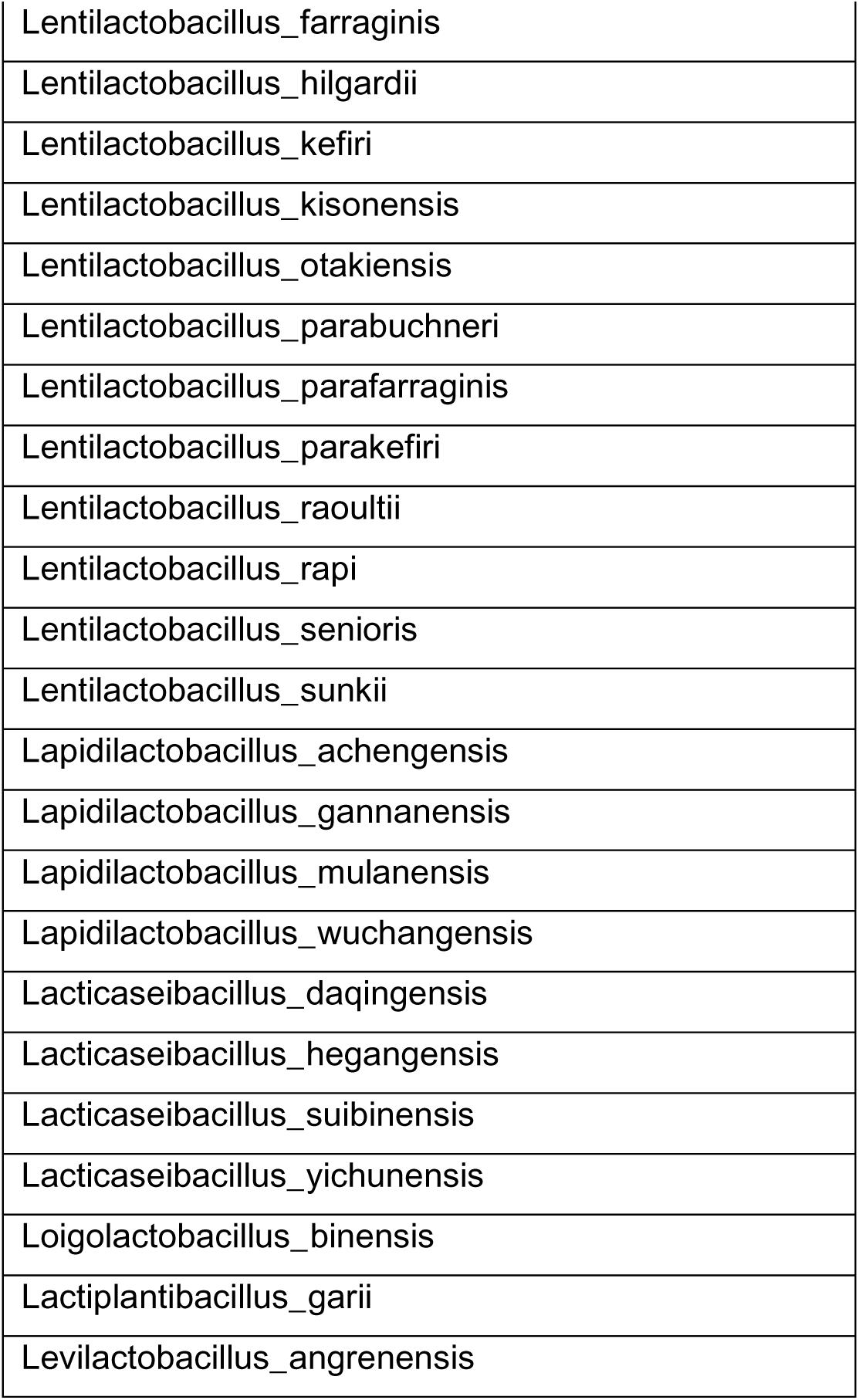
Lactobacillus strains used for phylogenetic analysis.

**Table S3.**
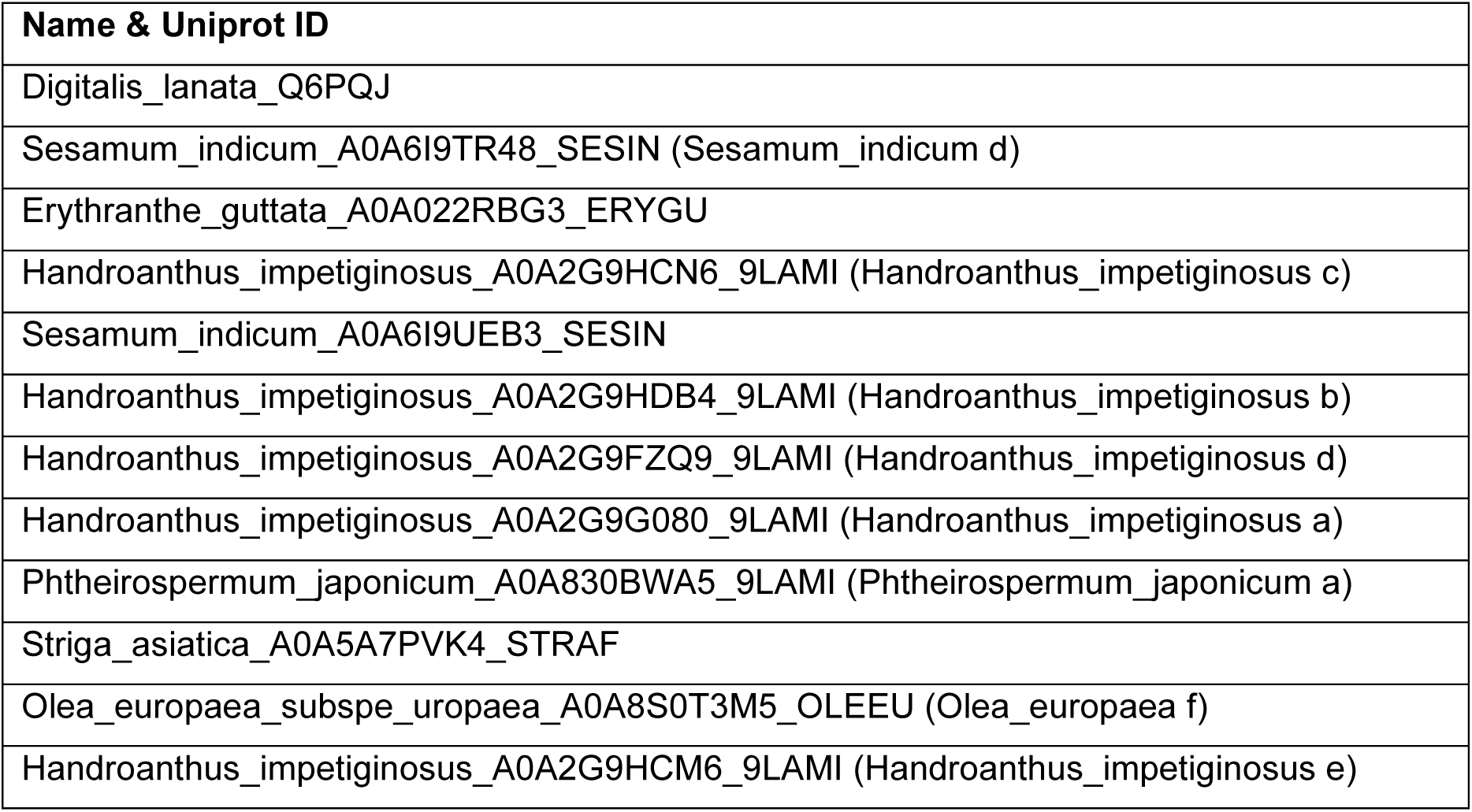
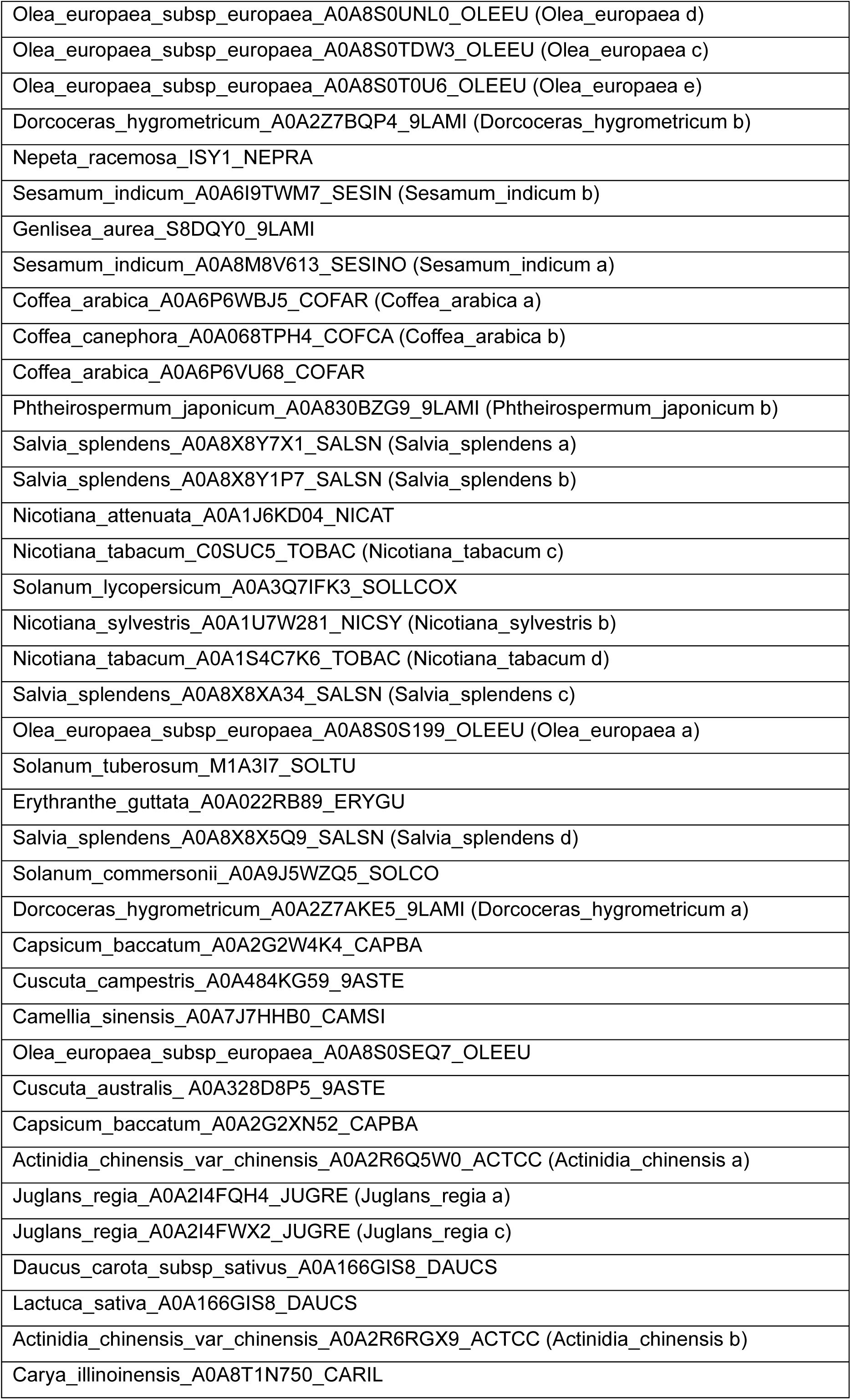
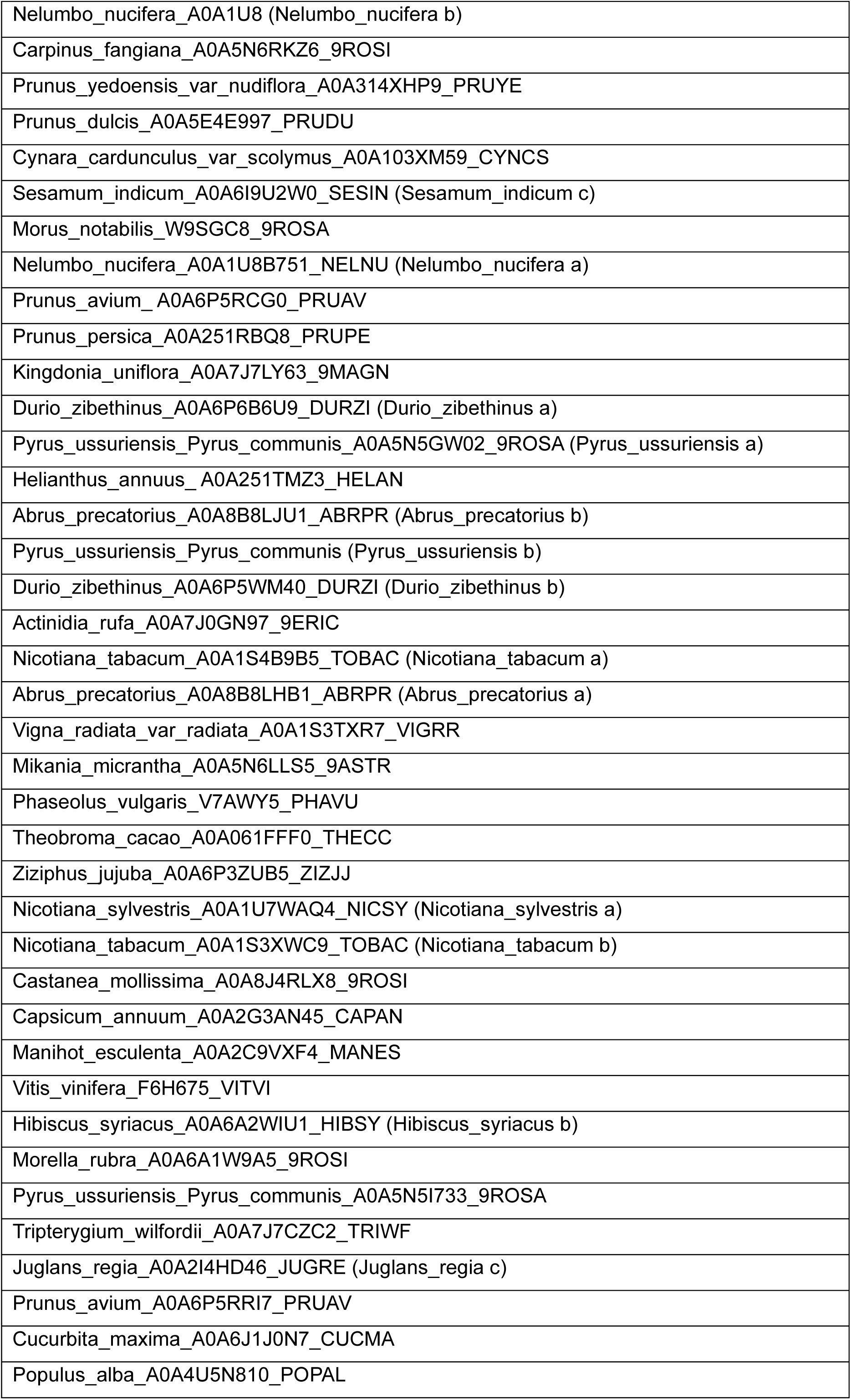
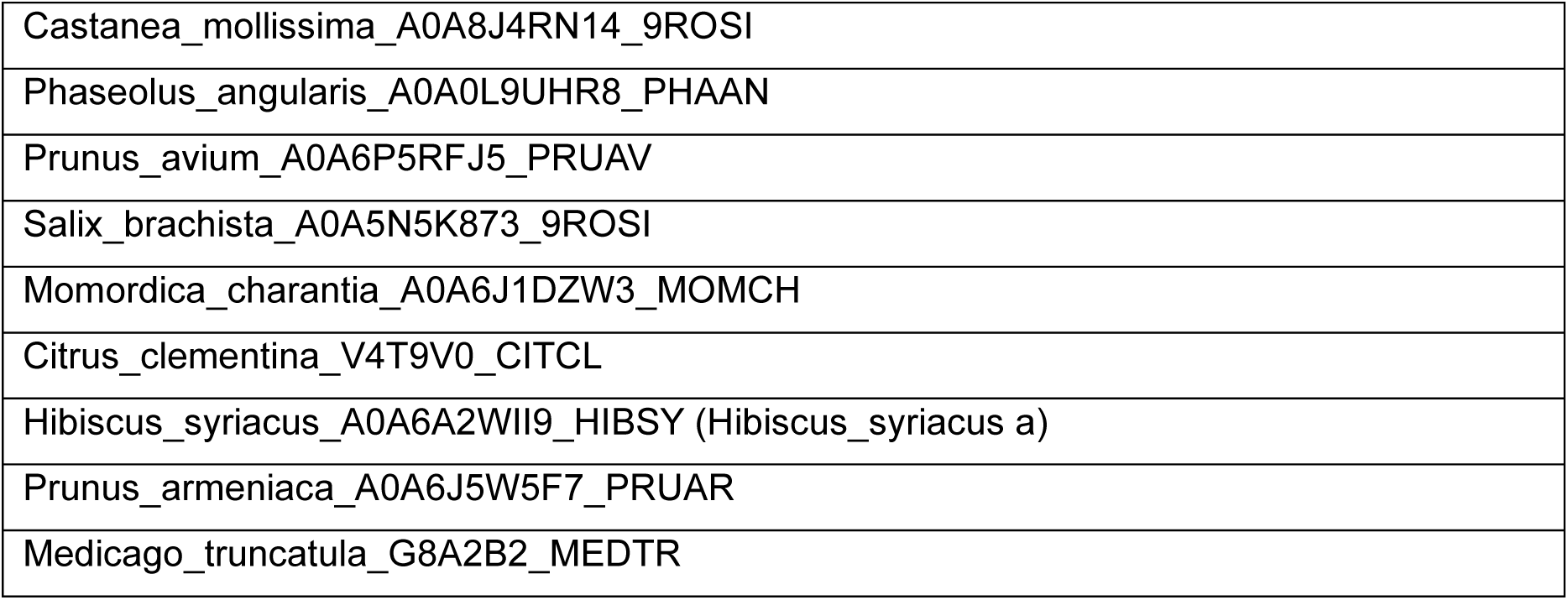
List of 5βChR plants homologs.

**Table S4.**
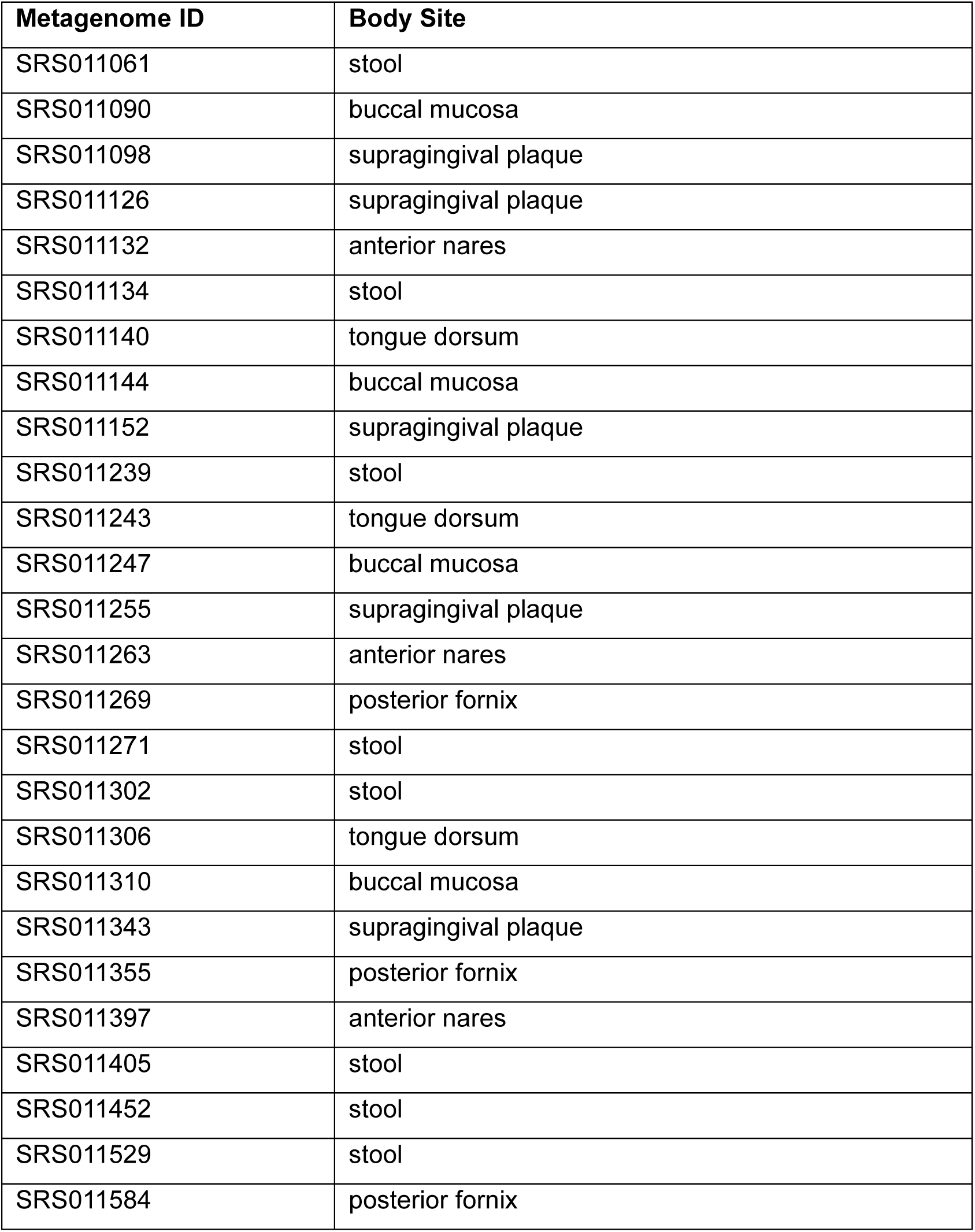
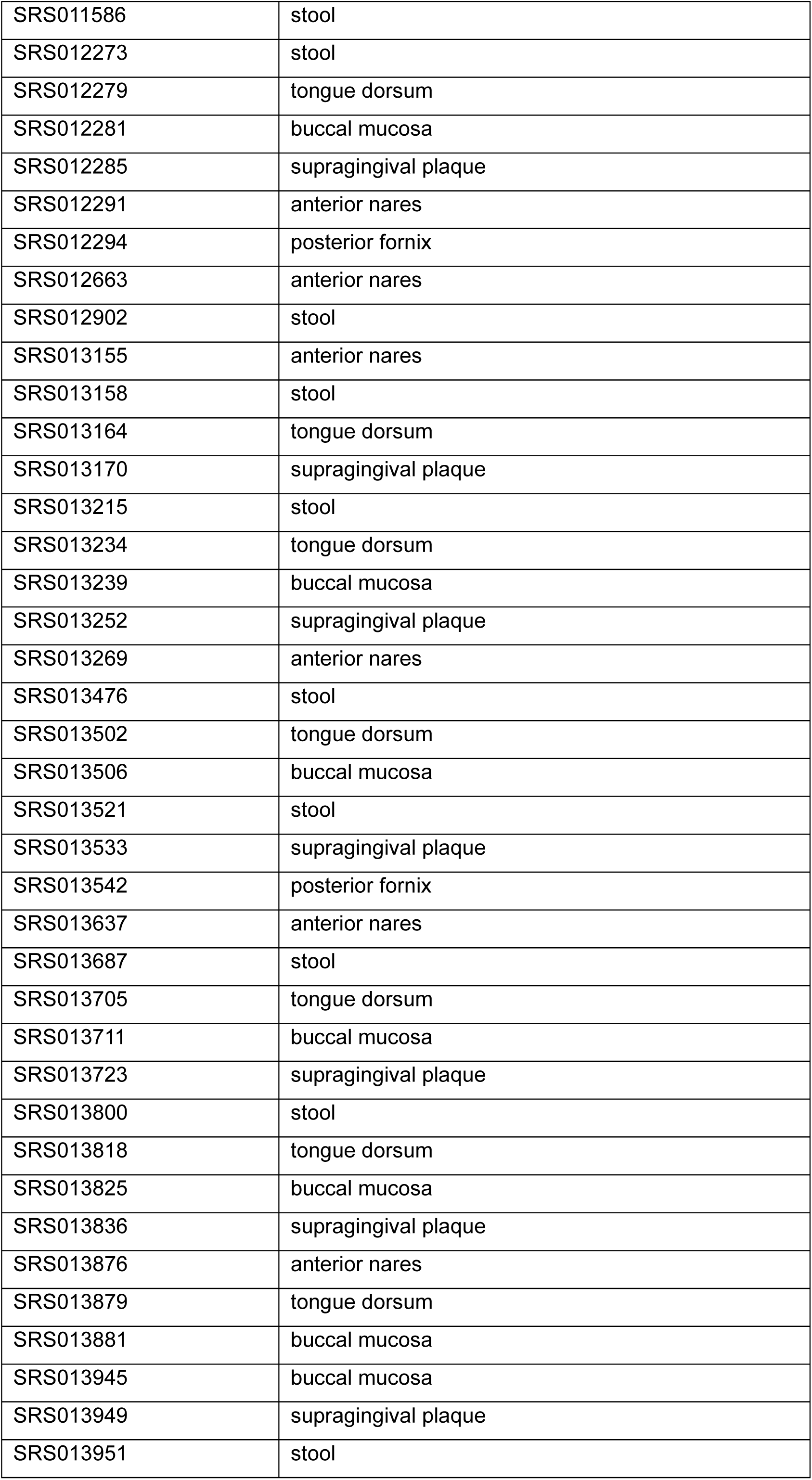
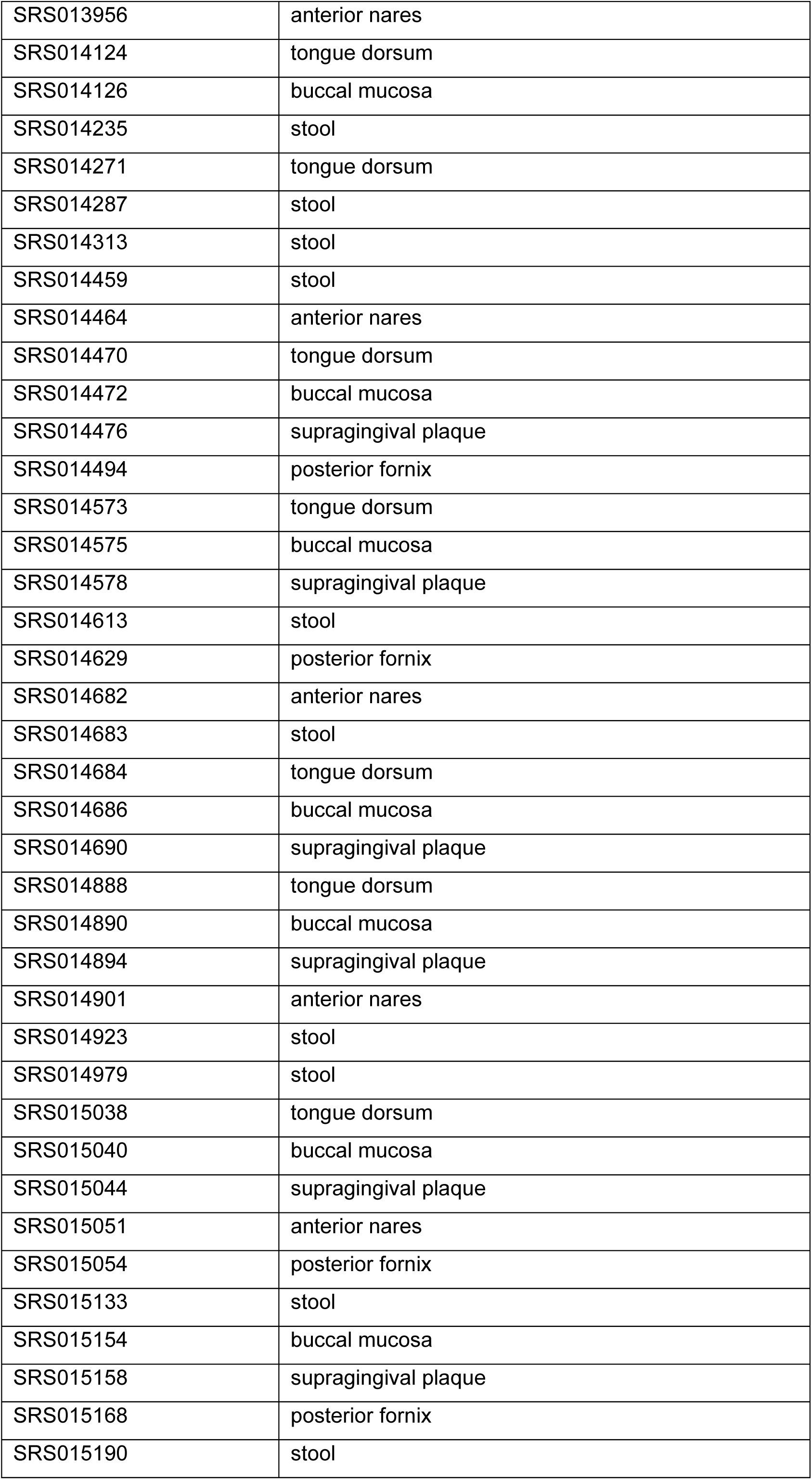
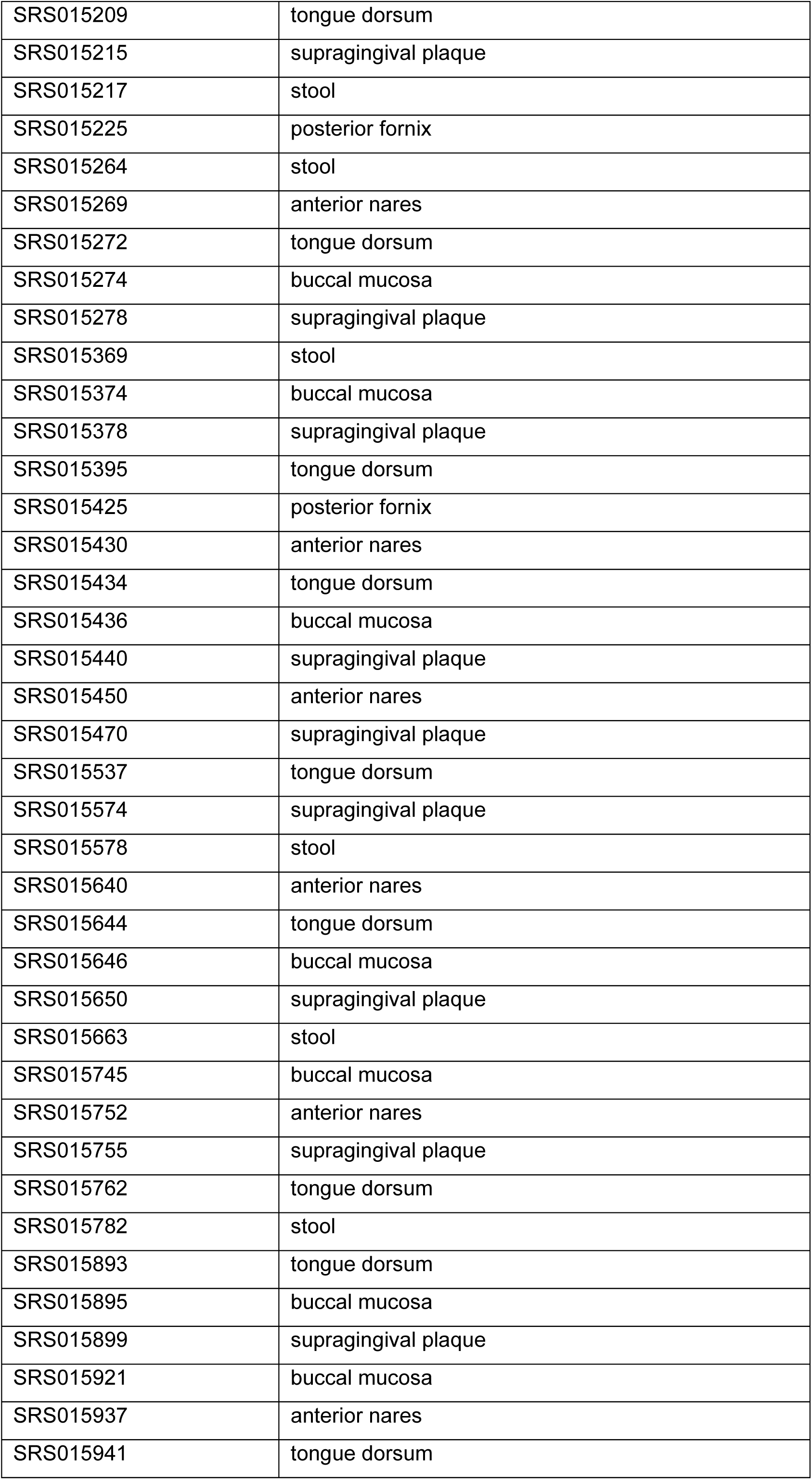
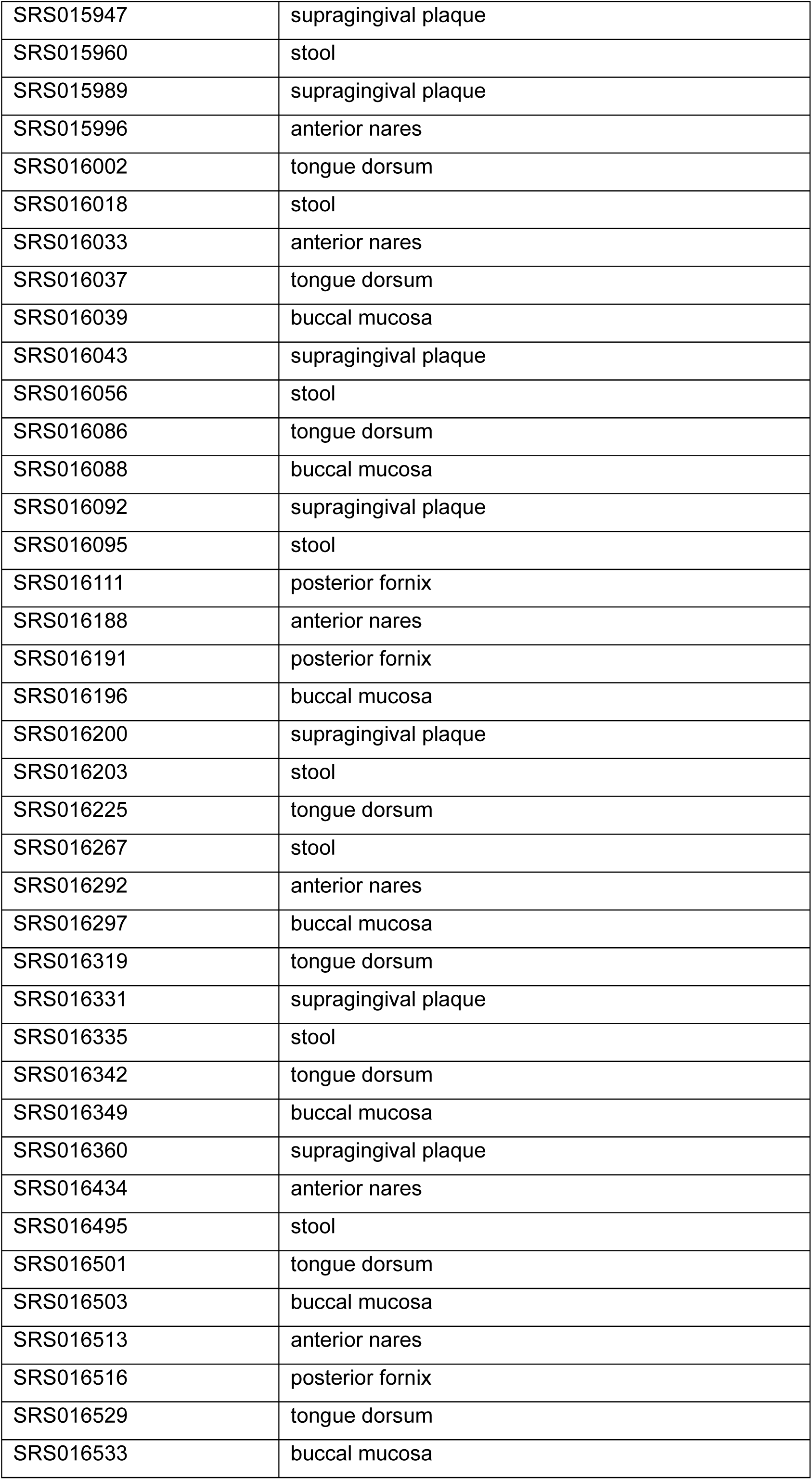
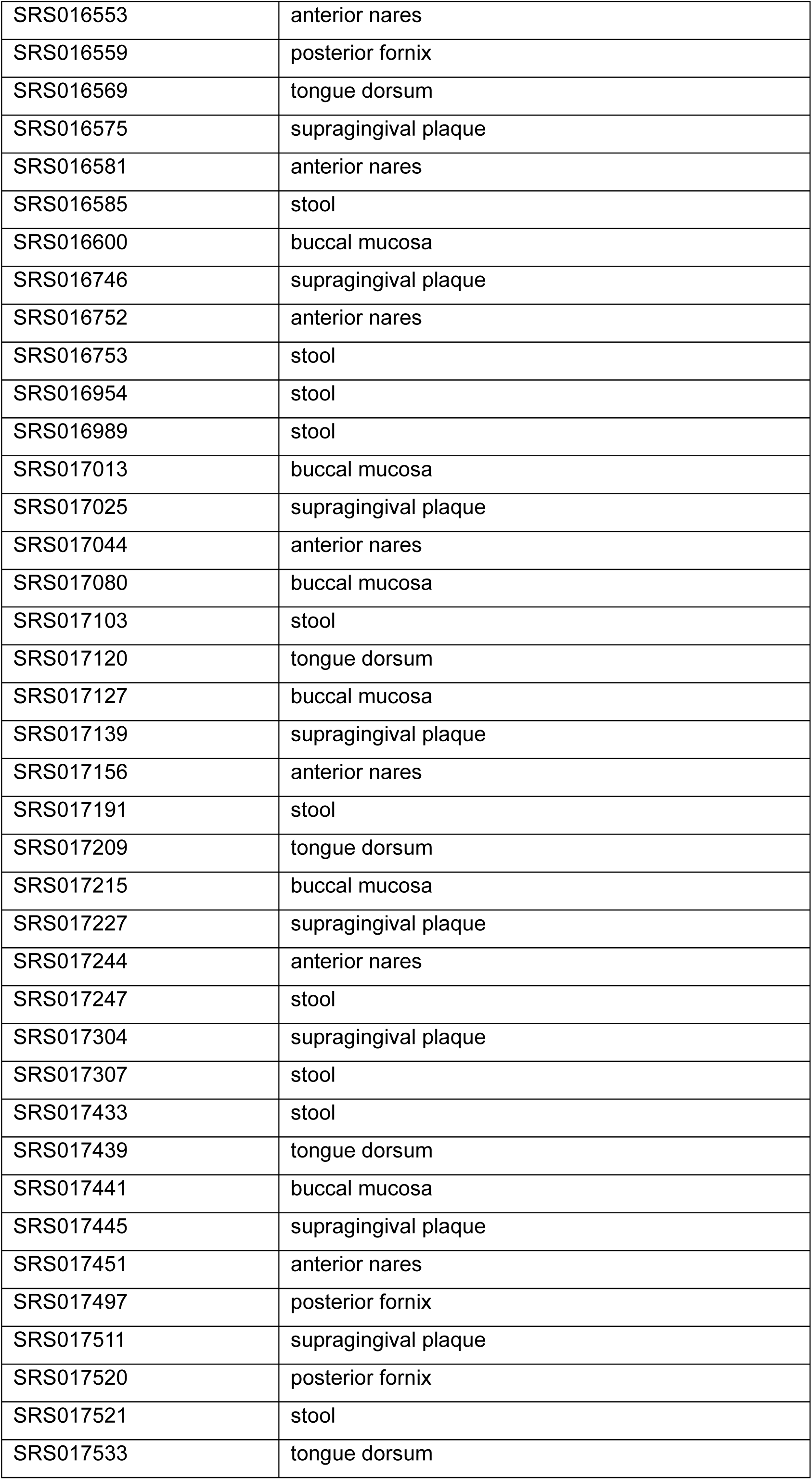
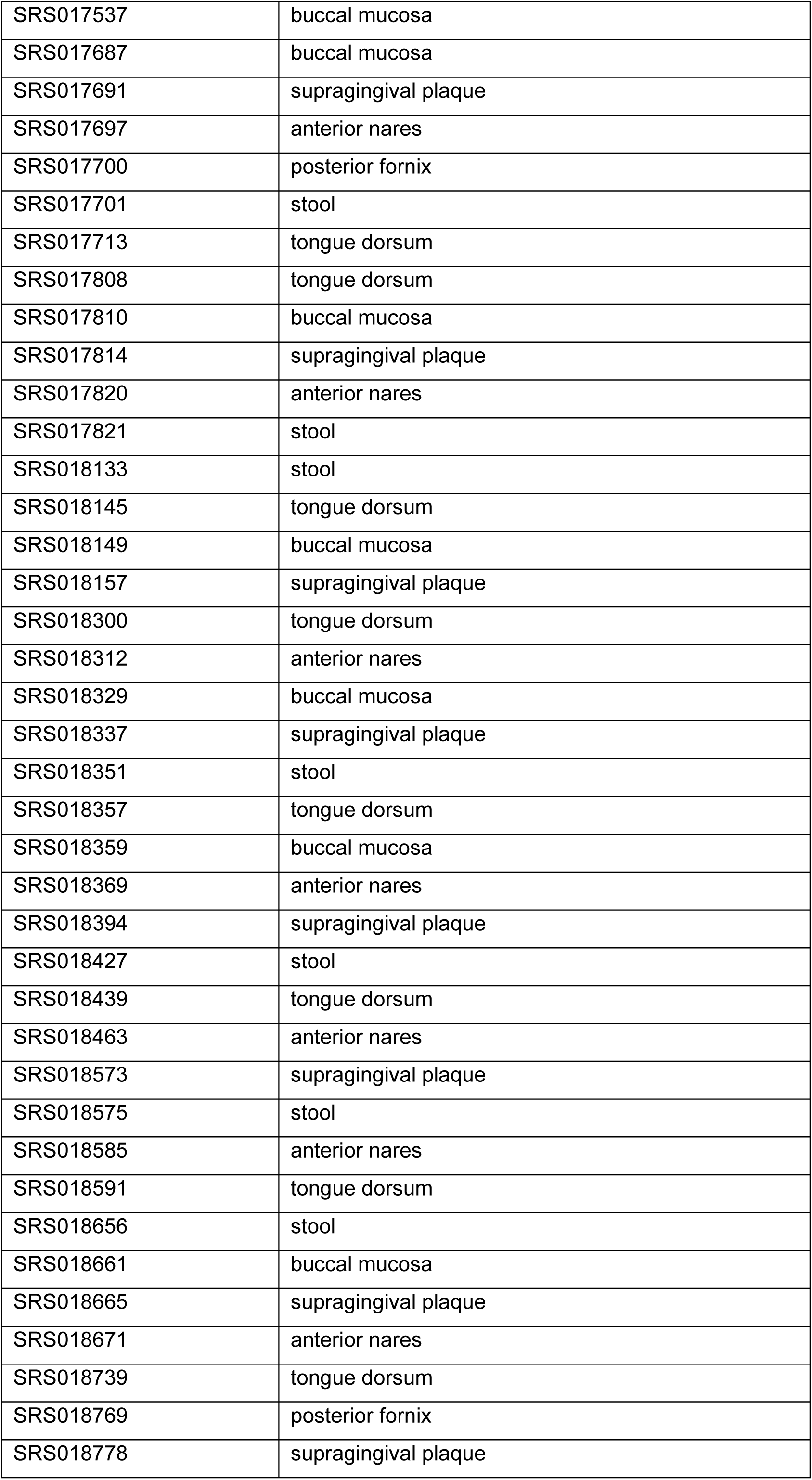
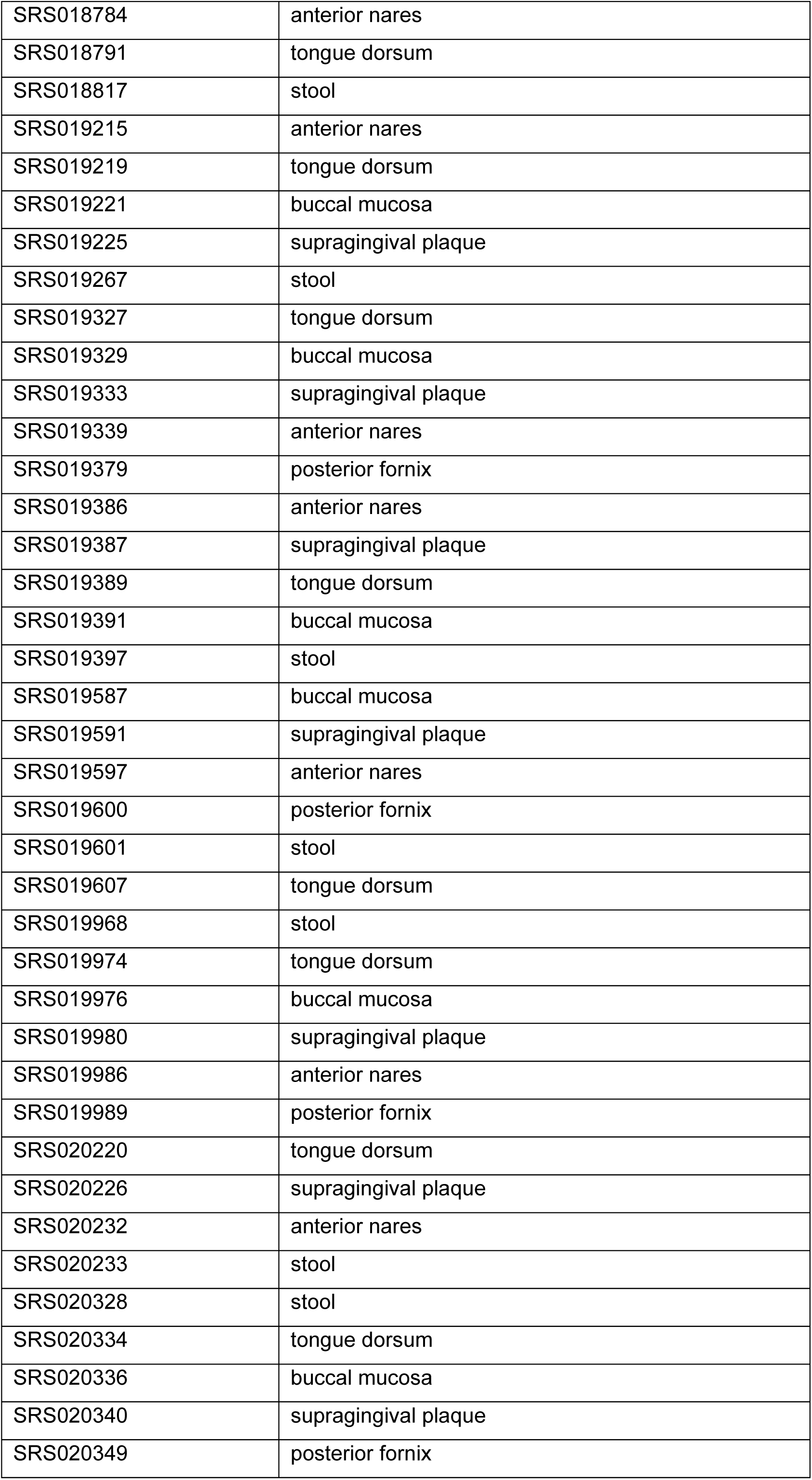
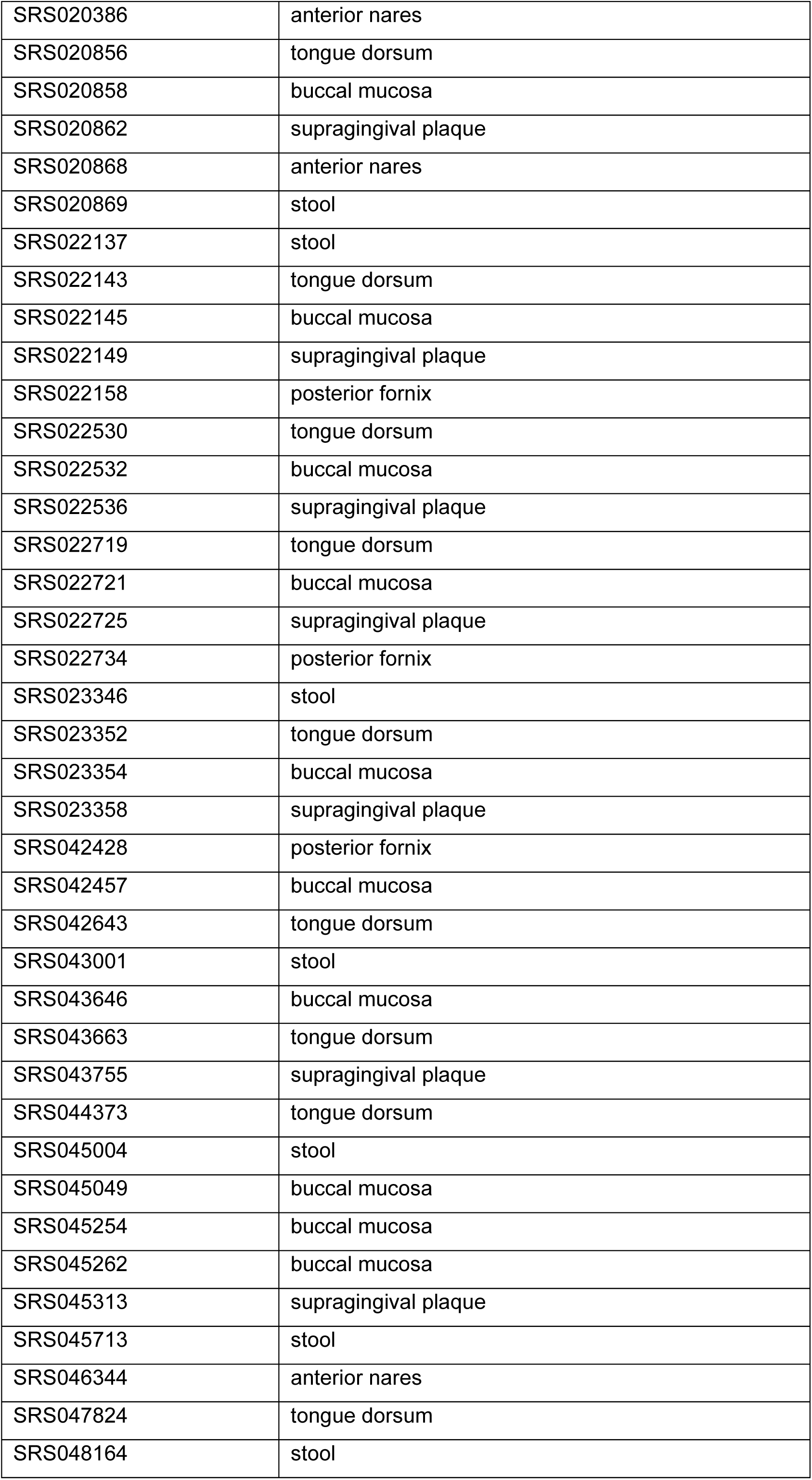
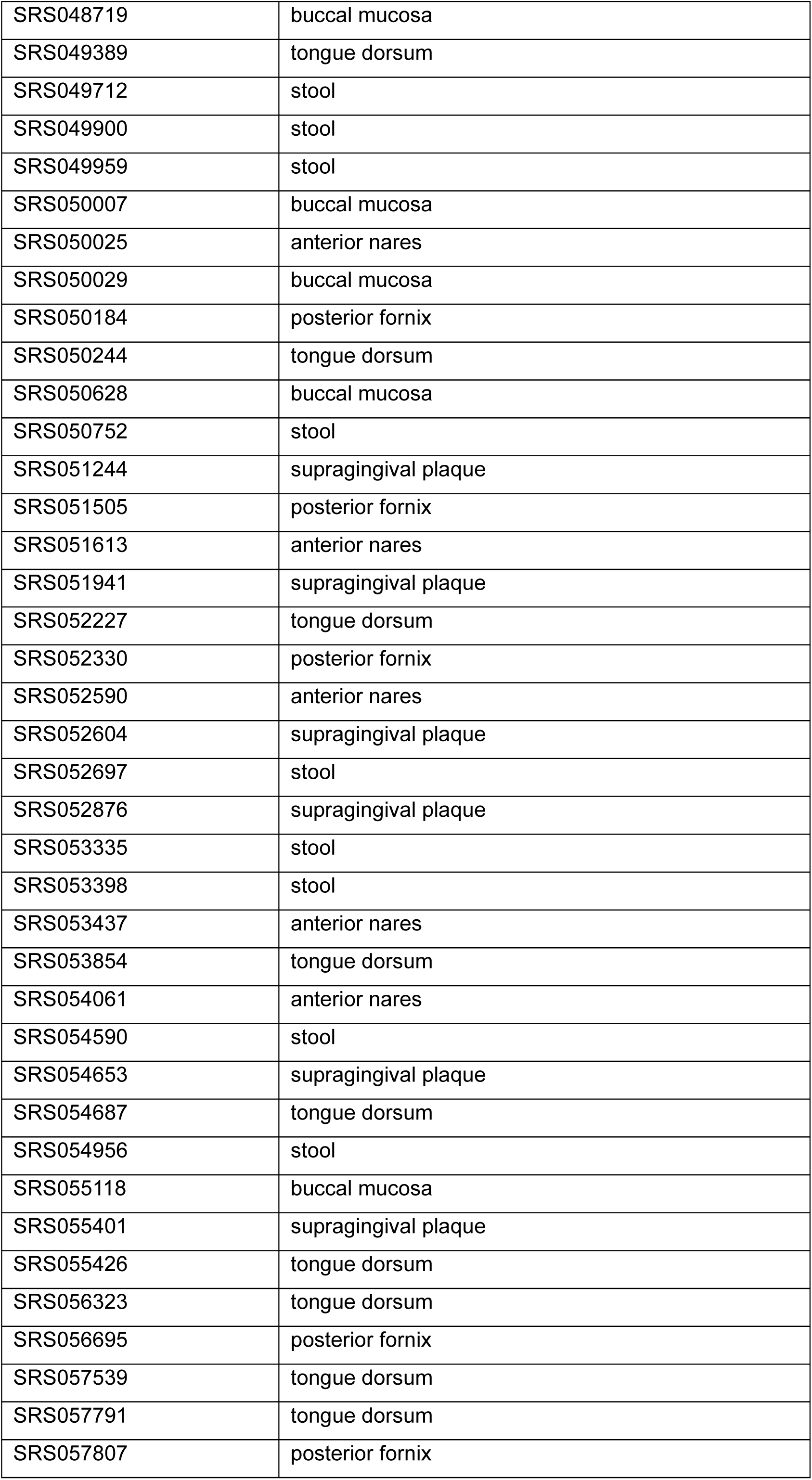
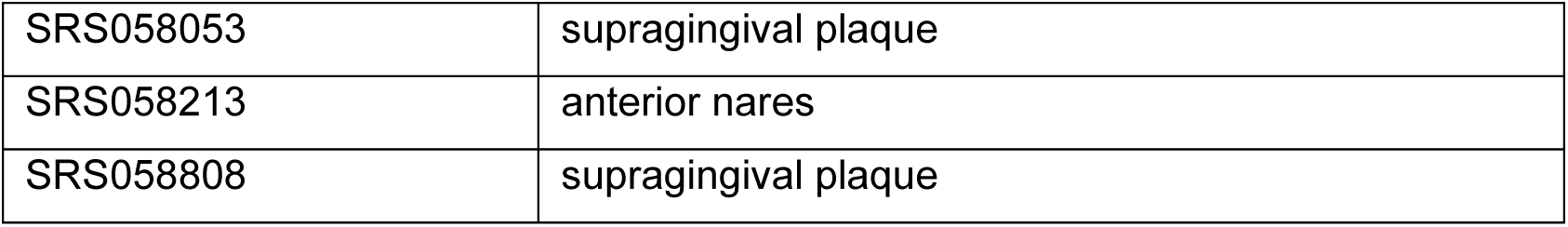
NCBI accession number of metagenome samples.

