## Supplementary information for "Discovery and mechanistic characterization of a probiotic-origin 3β-OH-Δ^5-6^-cholesterol-5β-reductase directly converting cholesterol to coprostanol"

**Table of Contents**

**1. Supplementary Figures…………………………………………………………...1-14**

**2. Supplementary Tables…………………………………………………………….15-35**

**1. Supplementary Figures**

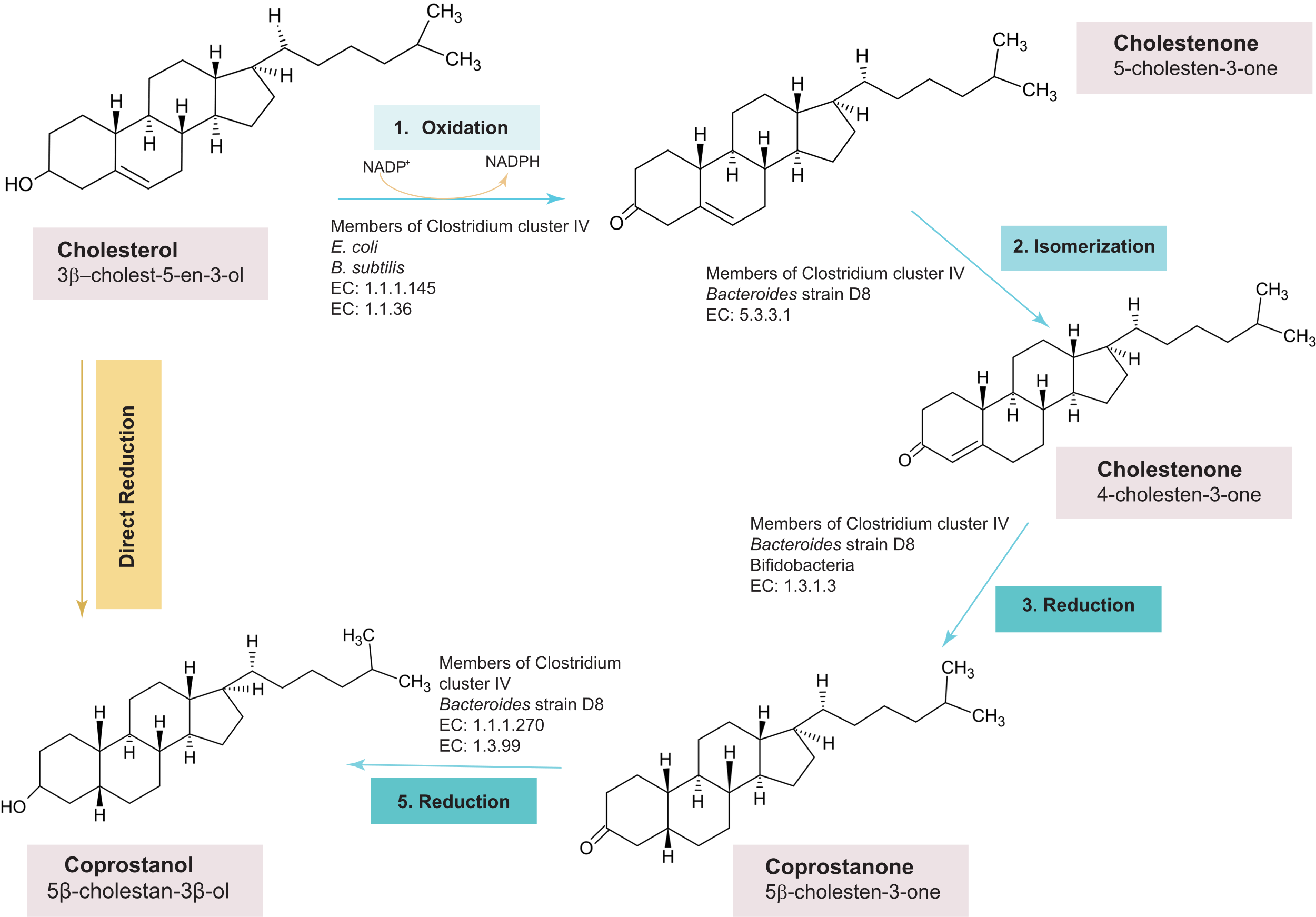

**Fig. S1** Biotransformation pathways of cholesterol to coprostanol by gut microbes.

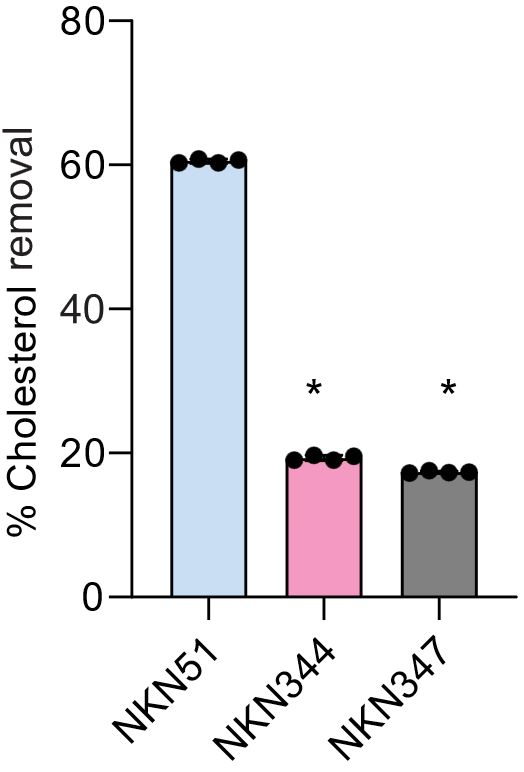

**Fig. S2**

Rudel & Morris cholesterol depletion assay representing in-vitro cholesterol removal by different *Lactobacillus* strains**.** Each point represents the biological replicate and *p- value* calculated by the Mann-Whitney test (**p*- 0.0286).

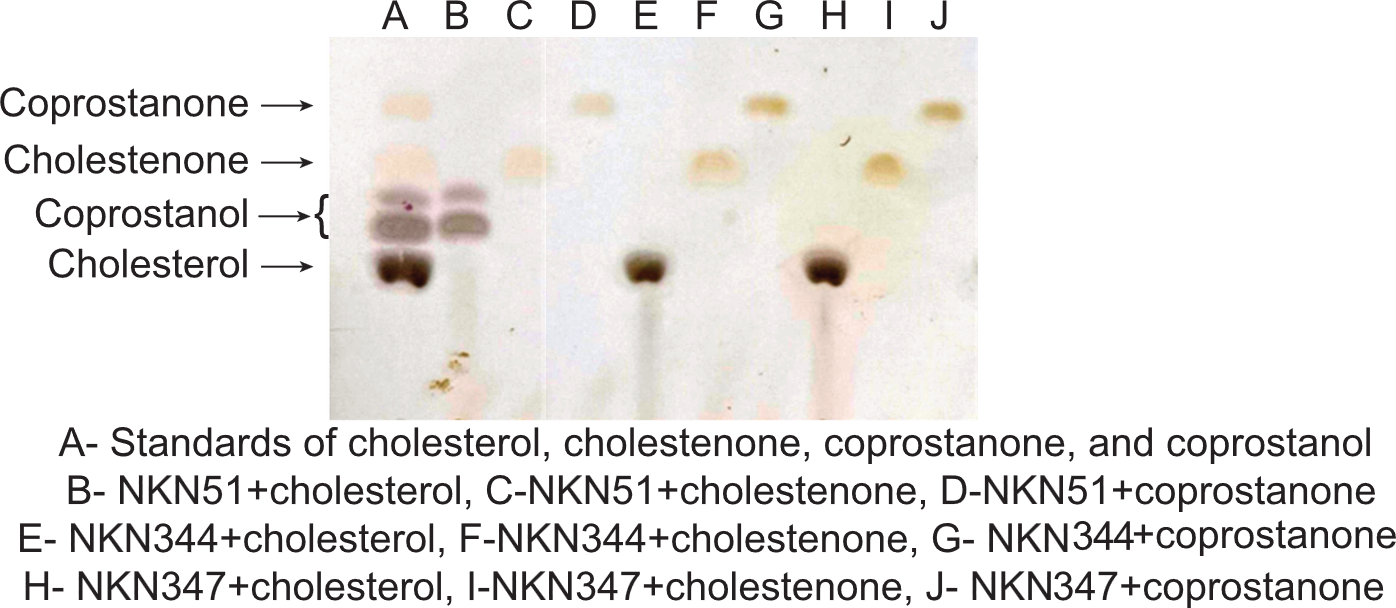

**Fig. S3**

Thin-layer chromatography (TLC) screening of cholesterol transformation to coprostanol by lactobacilli. The whole-cell lysate of different lactobacilli strains supplemented with NADP^+^ (100 µM) and either of substrates, cholesterol (100 µM), cholestenone (100 µM), or coprostanone (100 µM) for 12 h in anaerobic conditions.

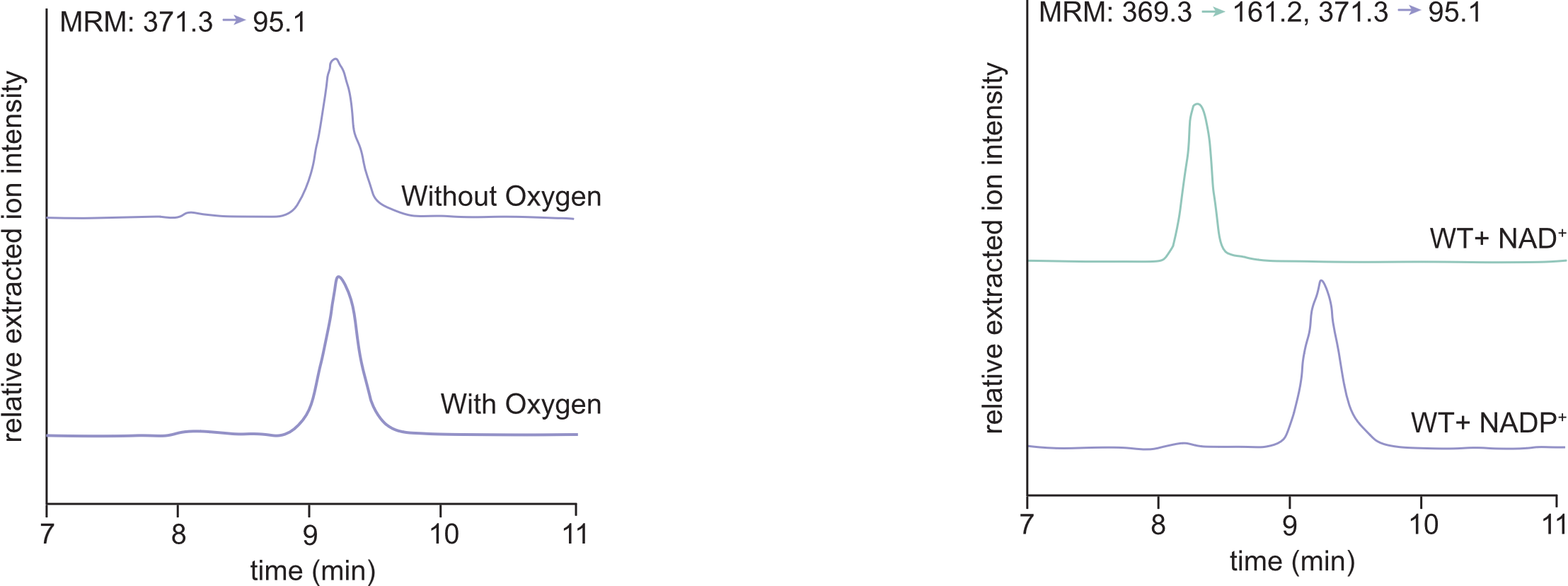

**Fig. S4**

Analysis of *L. fermentum* NKN51 lysate activity in microaerophilic conditions by Liquid chromatography Mass Spectrometry (LC-MS). The provided extracted ion chromatogram (EIC) represents the product of the whole-cell lysate assay of *L. fermentum* NKN51 supplemented with NADP^+^ (100 µM) and cholesterol (100 µM) over 12 h in microaerophilic and anaerobic conditions as a control.

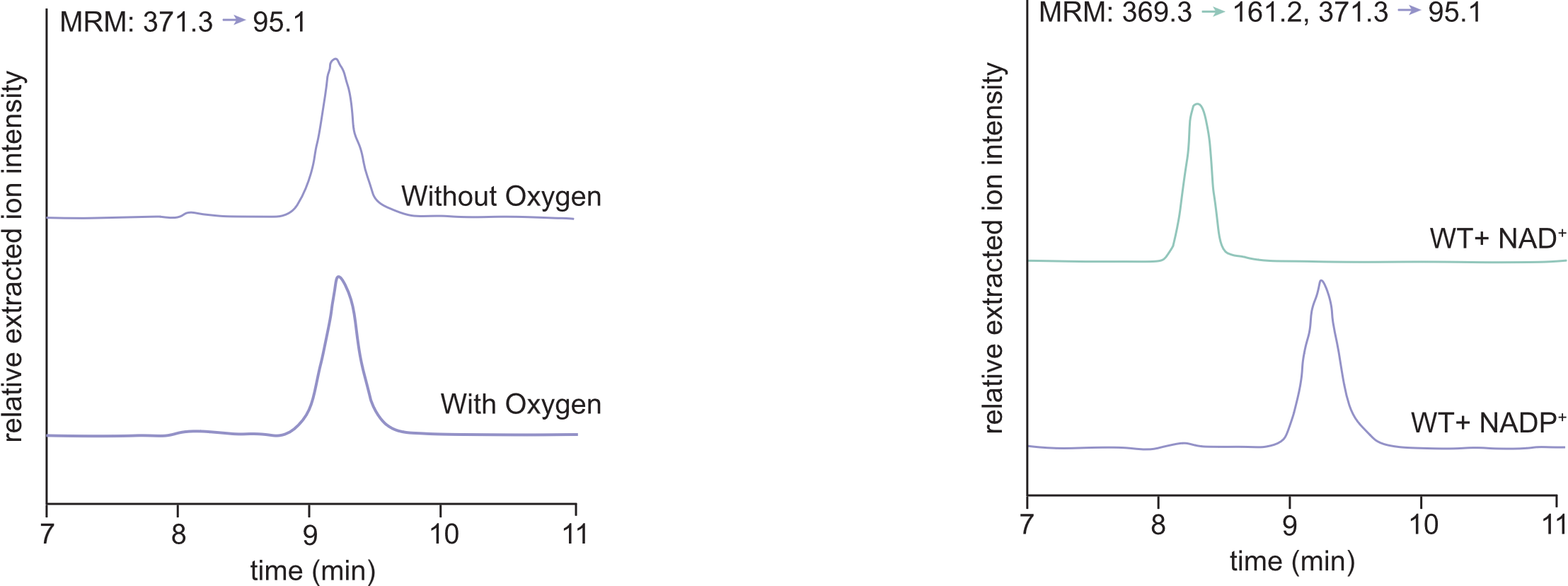

**Fig. S5**

Investigating the role of NAD^+^ as a cofactor for cholesterol metabolism by LC-MS. The reaction mixture of *L. fermentum* NKN51 whole-cell lysate with cholesterol (100 µM) and NAD^+^ (100 µM) or NADP^+^ (100 µM) for 12 h. No reaction was observed with NAD^+^ as a cofactor.

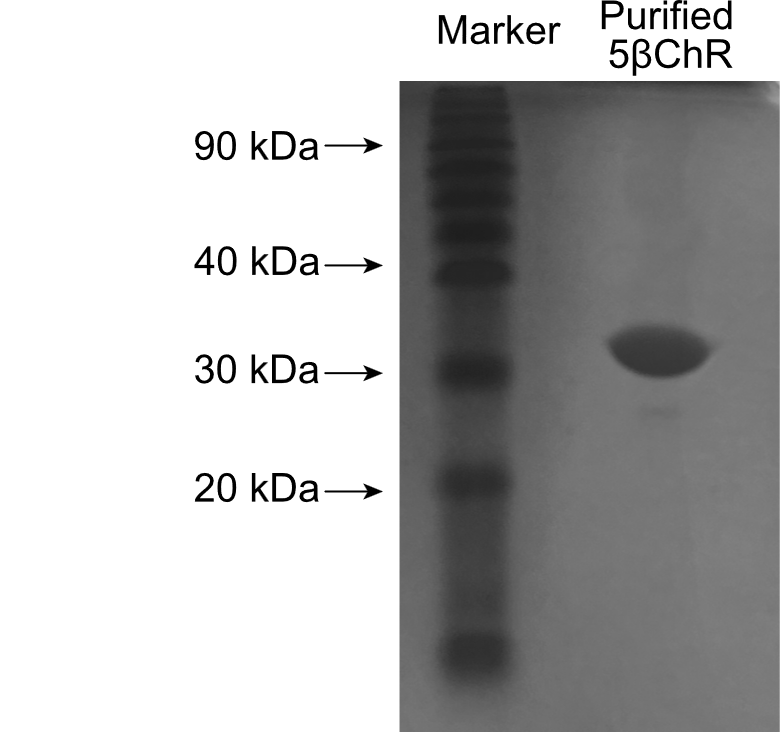

**Fig. S6**

Recombinant putative cholesterol dehydrogenase protein purification by immobilized affinity chromatography (IMAC). The SDS-PAGE depicts purified 5βChR protein by Ni-NTA-based affinity chromatography, with a theoretical molecular weight of 29.5 kDa.

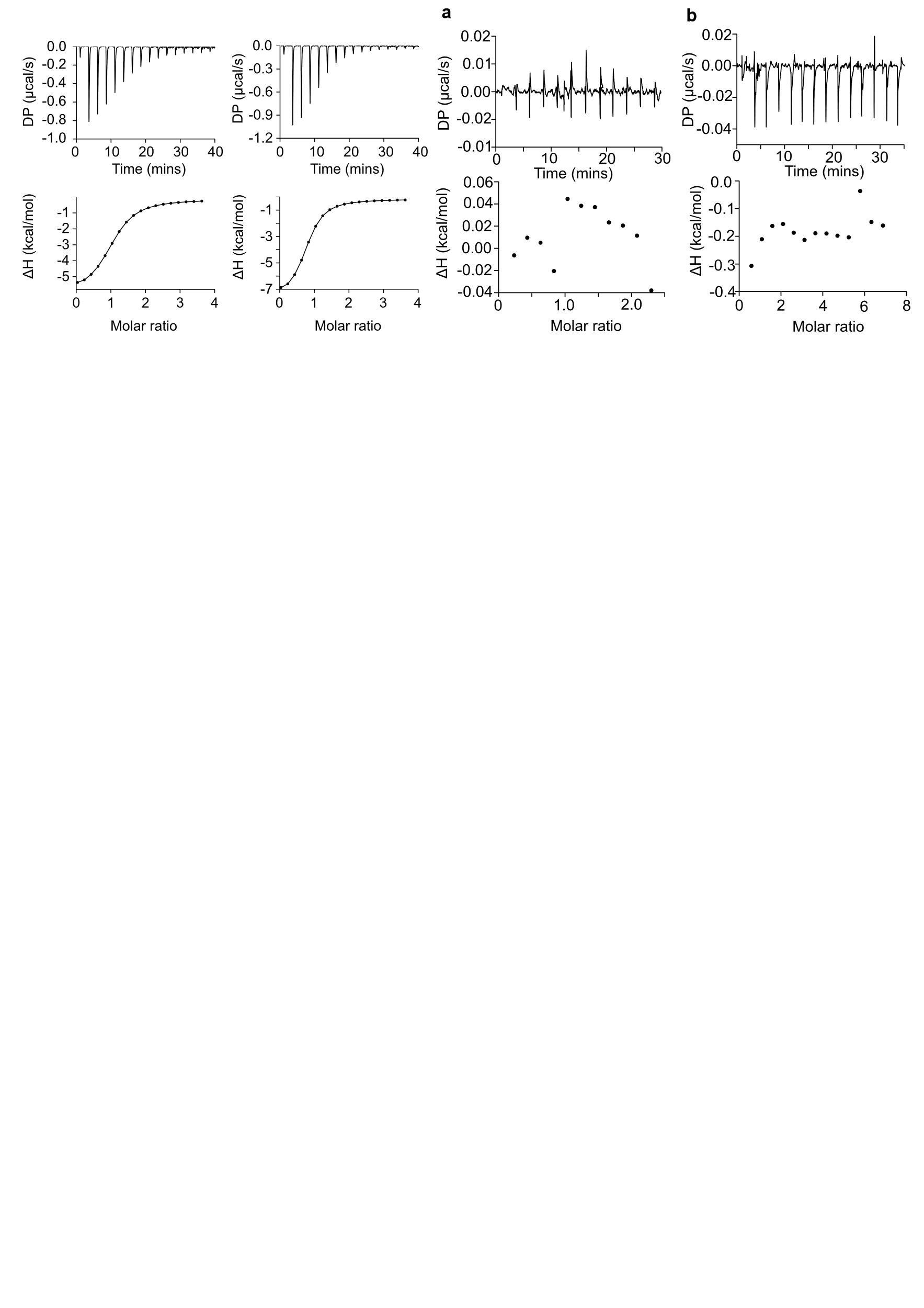

**Fig. S7**

Binding studies of 5βChR protein with substrates. The Isothermal Titration Calorimetry (ITC) of WT5βChR with (a) cholestenone and (b) coprostanone, wherein the points represent the normalized enthalpy change of the reaction at the different time intervals.

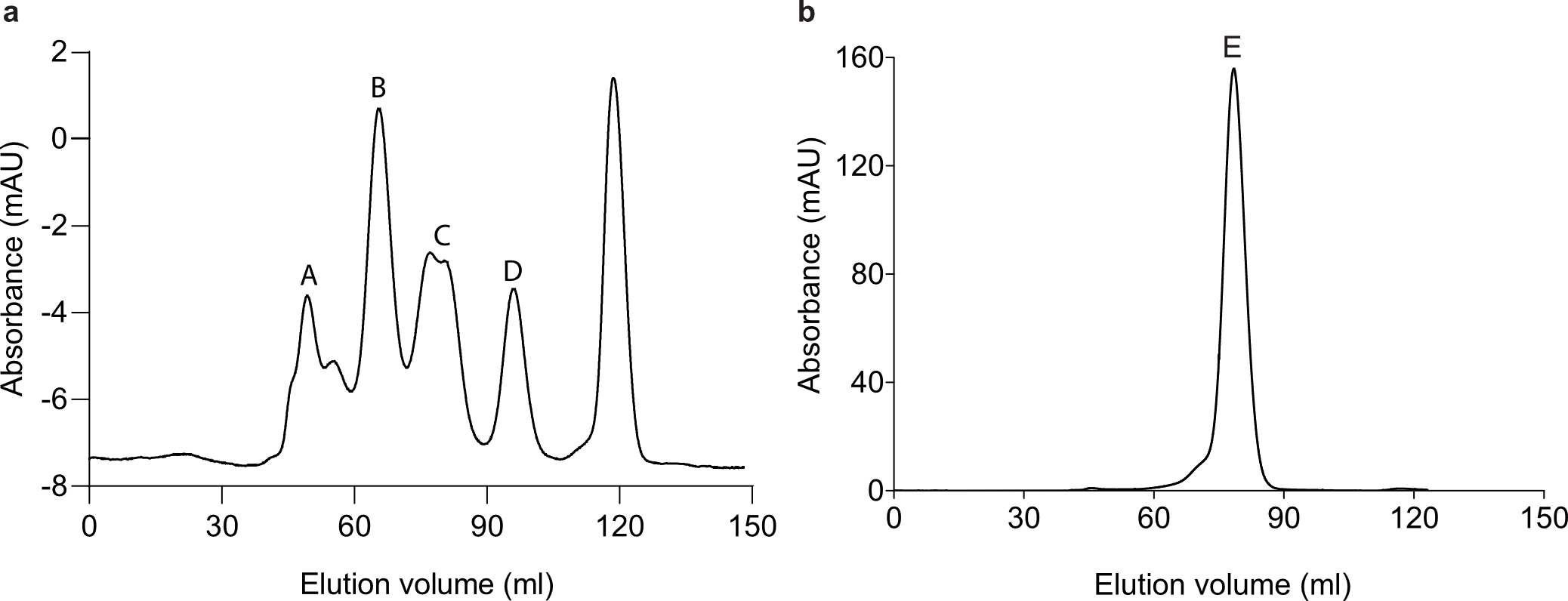

**Fig. S8**

Calculation of native molecular size of recombinant 5βChR. **(a).** The gel filtration graph of the standard mixture consists of various proteins represented by a distinct peak and their respective molecular weight along with elution volumes, which are as follows: A- Thyroglobulin (670kDa) 49.33ml, B- Gamma globulin (150 kDa) 65.63ml, C- Ovalbumin (44.5 kDa) 80.02 ml and, D- Ribonuclease A (13.7 kDa) 96.97 ml. **(b).** A 29.5 kDa recombinant cholesterol dehydrogenase (E) eluted at 78.43 ml, depicts its dimeric form.

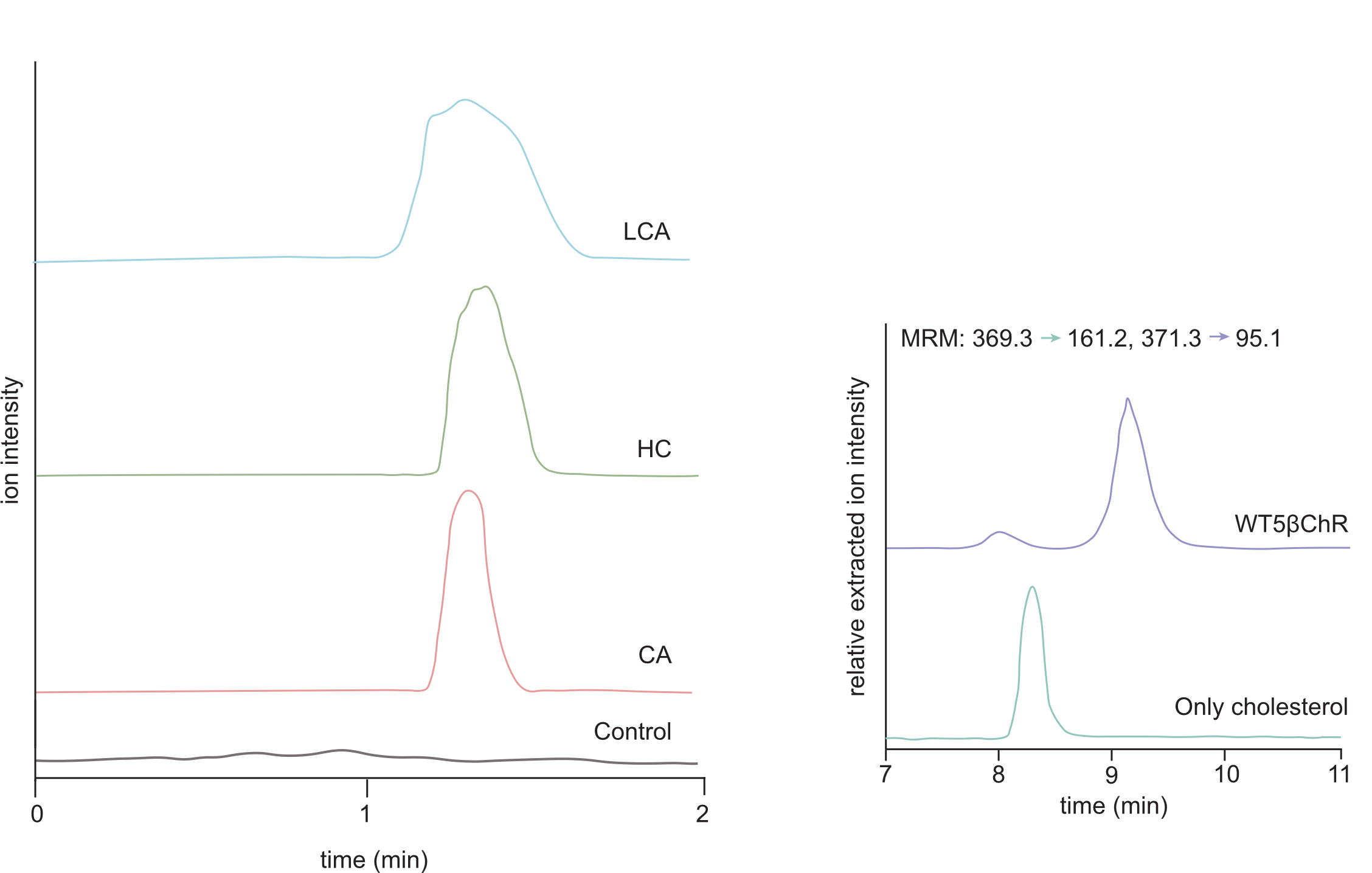

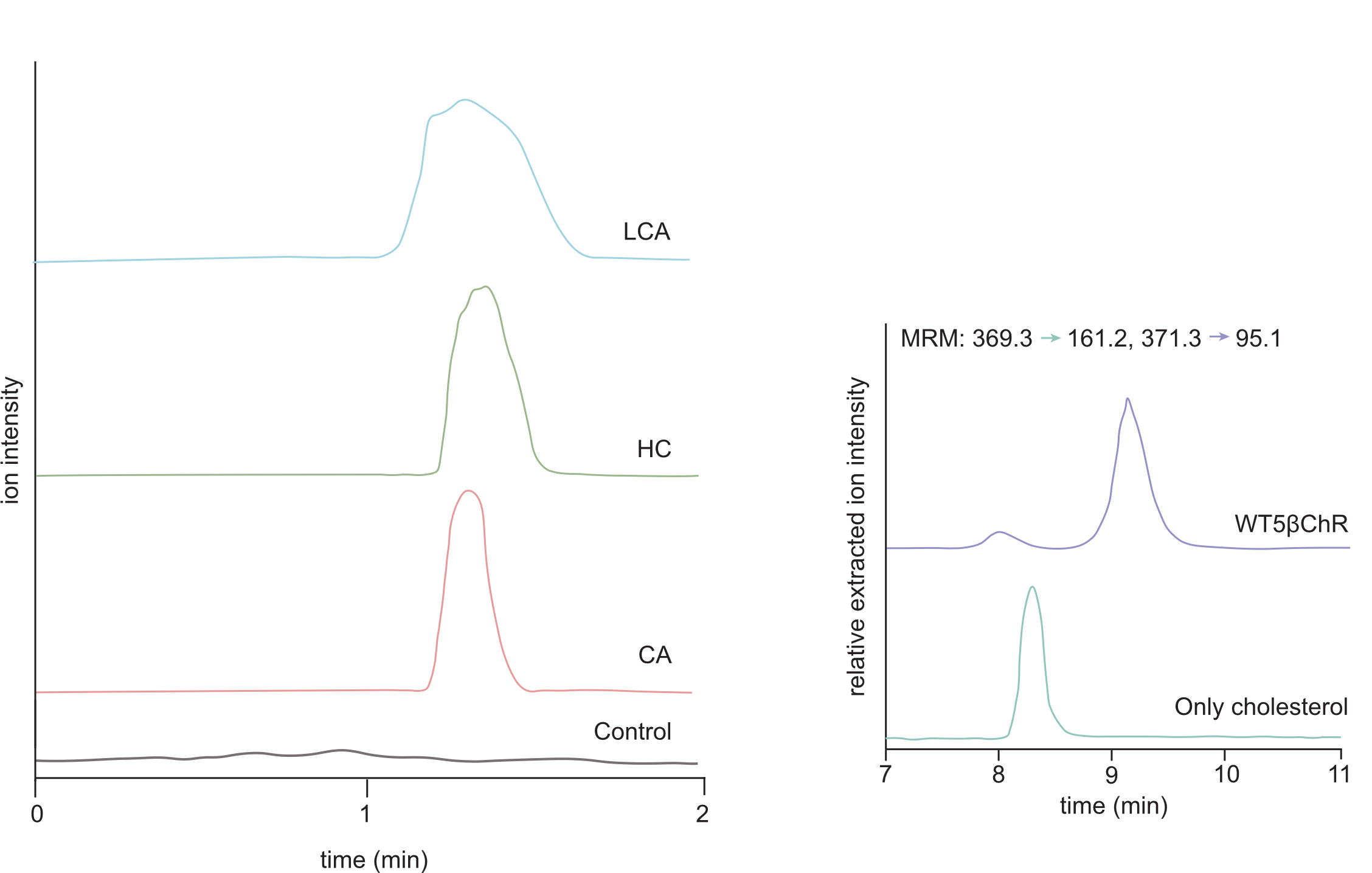

**Fig. S9**

5βChR enzyme activity with a similar structure compound like cholesterol. The total ion chromatograph demonstrates no reaction observed by 5βChR incubated with either 100 µM cholic acid (CA), hydrocortisone (HC), or lithocholic acid (LCA). EIC shows the coprostanol detection from the reaction of 5βChR incubated with cholesterol (100 µM) and NADP^+^ (100 µM).

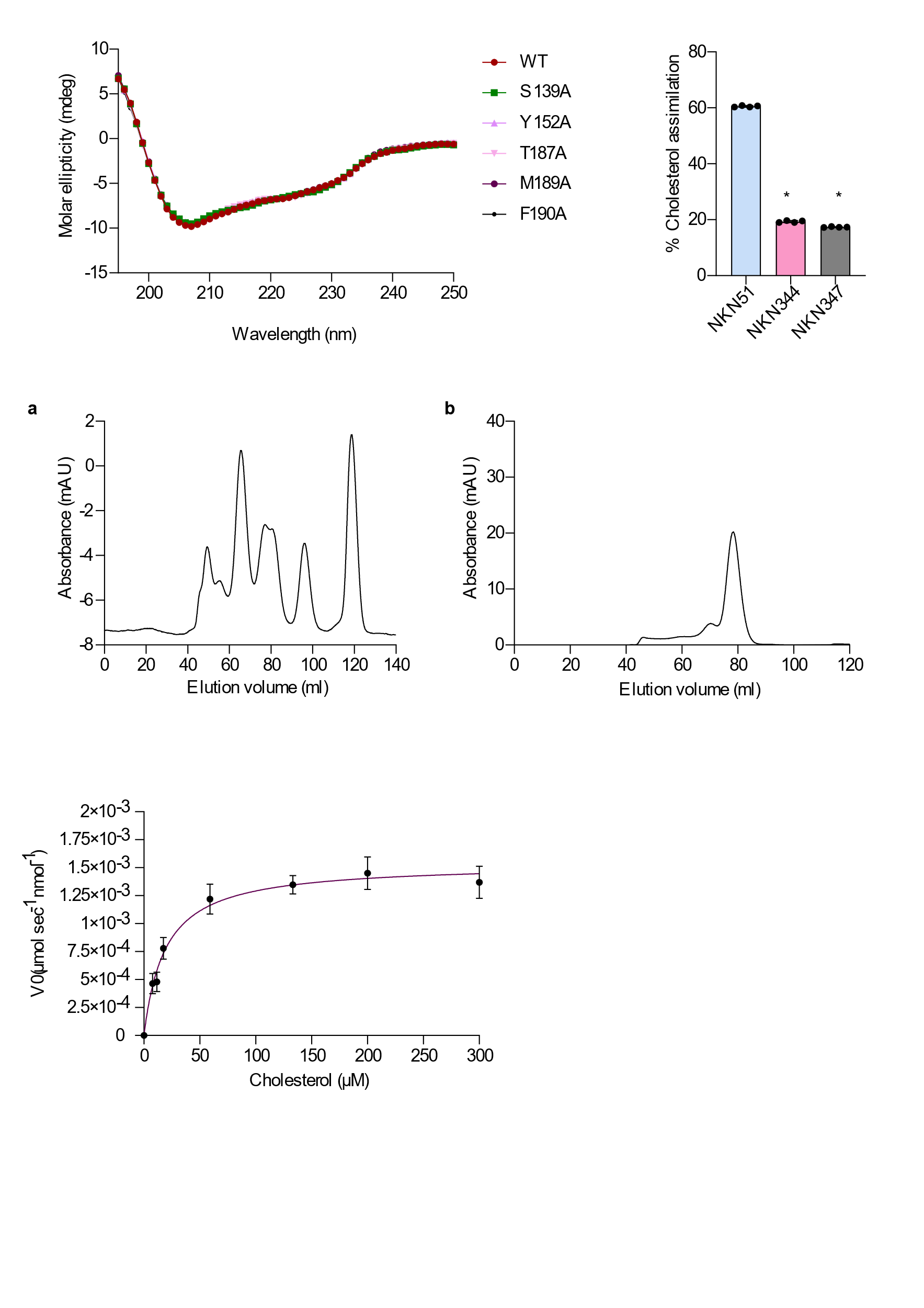

**Fig. S10**

Enzyme kinetics of WT5βChR. The Michalis-Menton curve is determined from the reaction of cholesterol with 5βChR. The saturation curve predicts the kinetics of cholesterol reduction by recombinant 5βChR enzyme with *Km*- 19.09 µM and *V_max_*- 15.35x10^-6^ µmol sec^-1^ nmol^-1^.

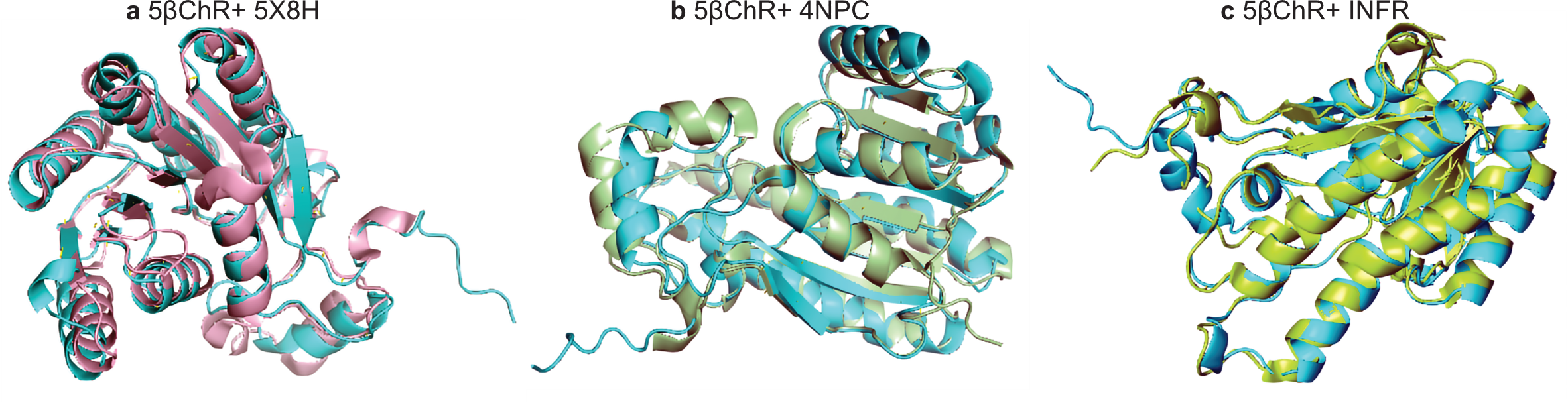

**Fig. S11**

Structural comparison of 5βChR with its predicted homologs. According to the DALI server, the closest structural homologs of 5βChR are SDR reductase (PDB ID: 5X8H), Sorbitol dehydrogenase (PDB ID: 4NPC), and Putative oxidoreductase RV2002 (PDB ID: 1NFR). These three enzymes are superimposed onto 5βChR with root square mean value (r.m.s) deviation of 1.3, 1.3, and 1.2 Å, respectively.

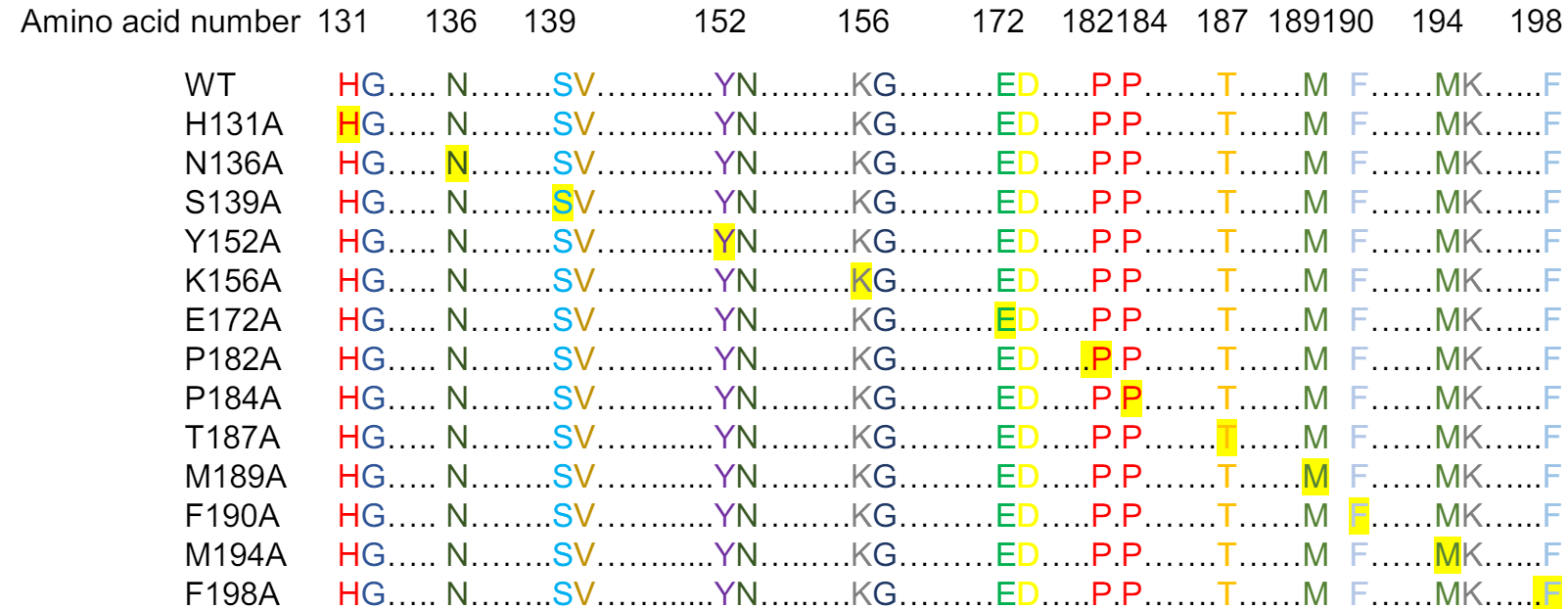

**Fig. S12**

Site-directed mutagenesis of WT5βChR. The highlighted amino acids represent the mutation in conserved residues of WT5βChR. N-terminus residues, H131, N136, S139, Y152, K156 and E172 mutated to alanine. C-terminus residues, P182, P184, T187, M189, F190, M194, and F198, are converted to alanine for cholesterol binding analysis.

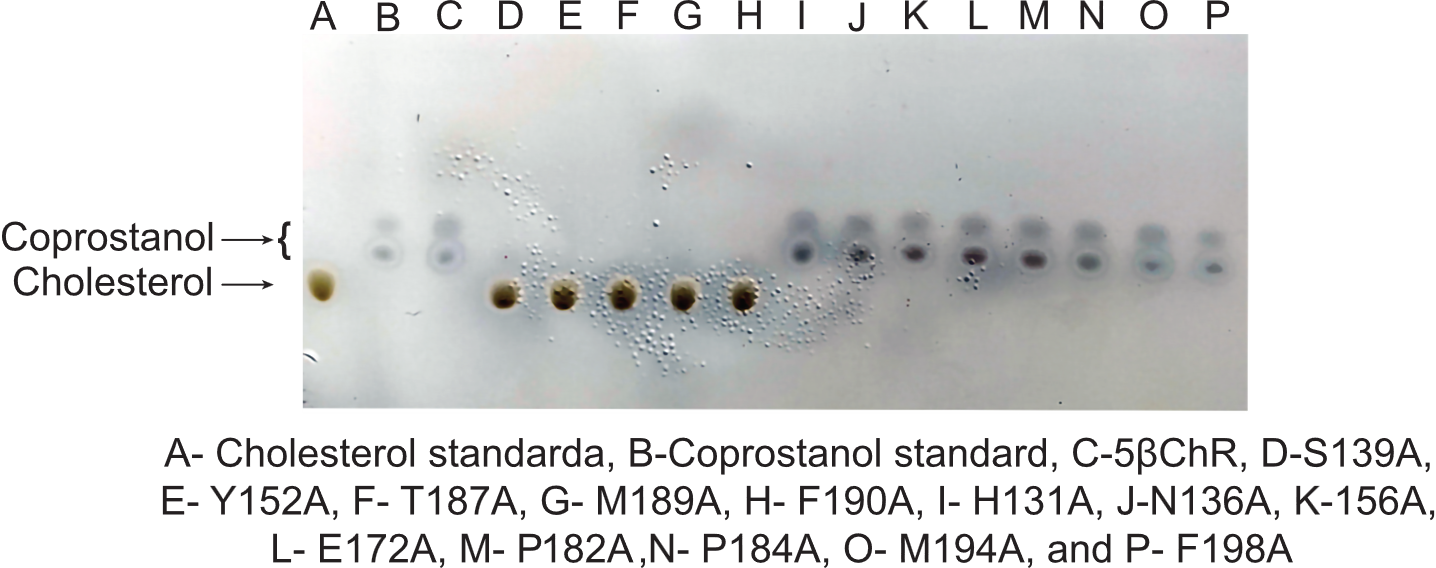

**Fig. S13**

Cholesterol transformation profile of WT and mutant 5βChR**.** The reaction mixture of 5βChR WT and mutants with cholesterol (100 µM) and NADP^+^ (100 µM) are analyzed through TLC. S139A, Y152A, T187A, M189A, and F190A show the spot equivalent to cholesterol. H131A, N136A, 156A, E172A, P182A, P184A, M194A, and F198A highlighting spots corresponding to coprostanol.

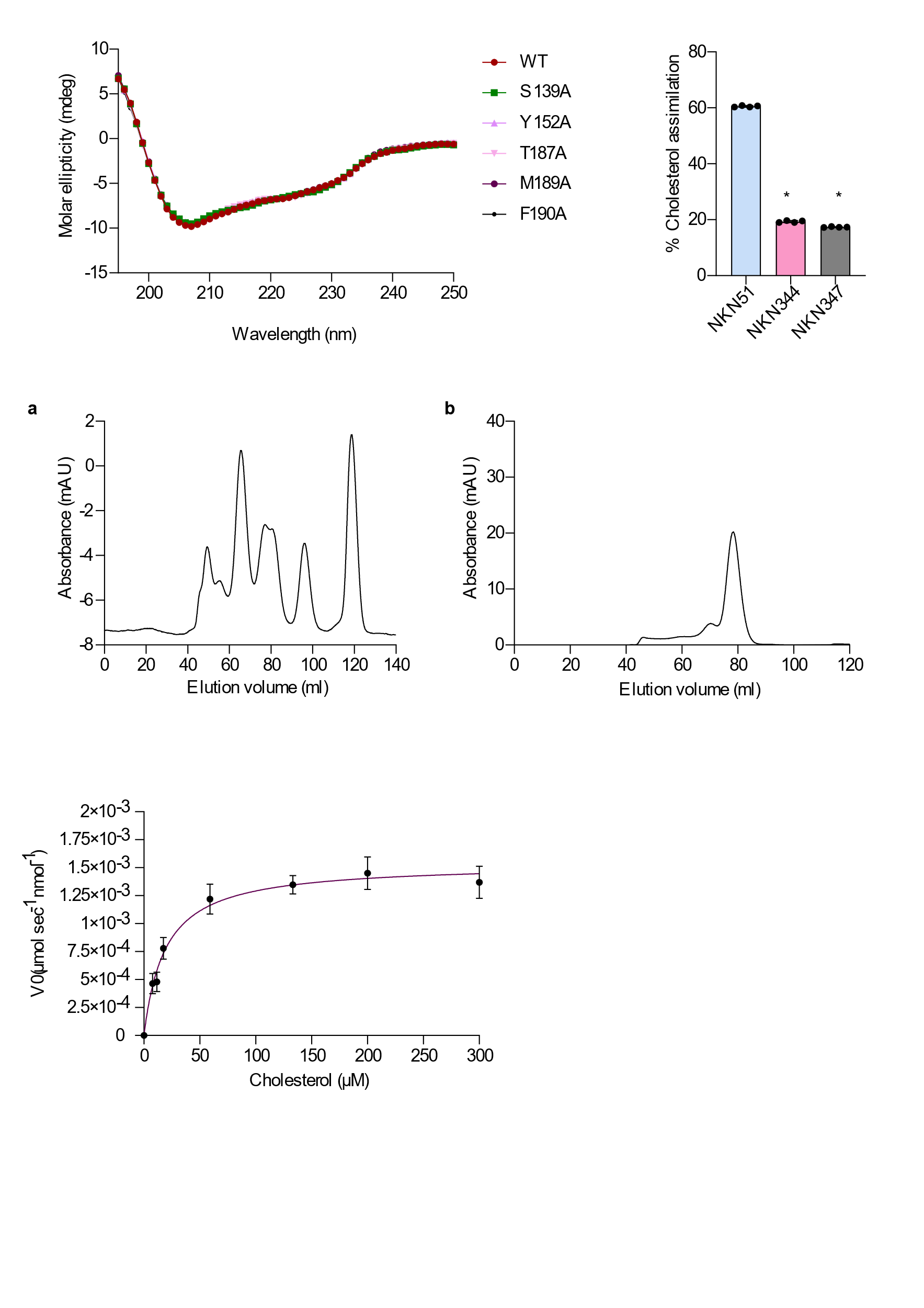

**Fig. S14**

Circular Dichroism spectra of 5βChR and its mutants. The graph represents the secondary structure profiles of WT5βChR, S139A, Y152A, T187A, M189A, and F190A.

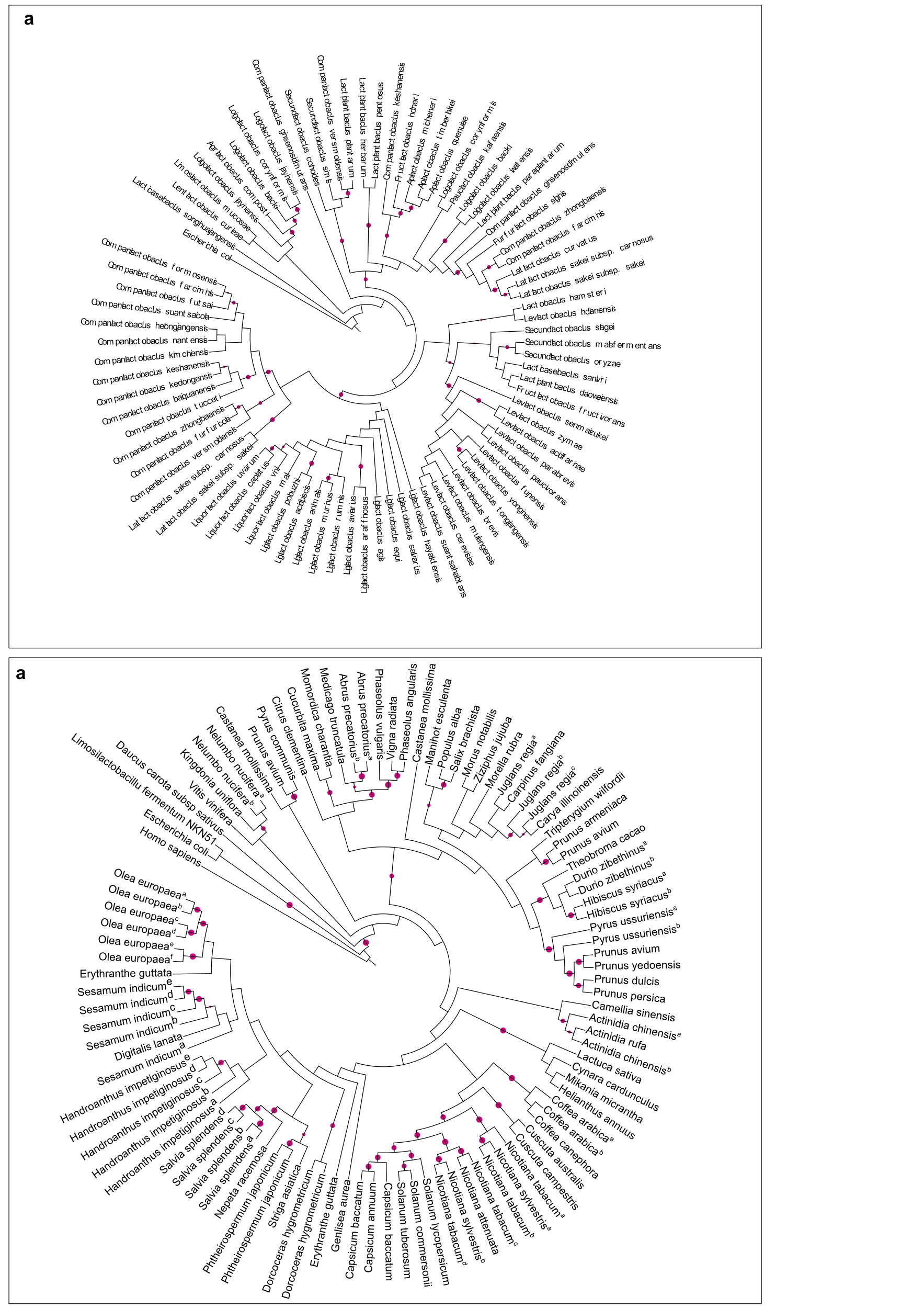

**Fig. S15**

Mapping the prevalence of 5βChR within plants by phylogenetic profiling. **(a).** The evolution analysis of 5βChR homologs in plants (~100) is acquired from UniProt in RAxML using the PROTCATWAG model with human 5β reductase as an outgroup. 5βChR and plant 5β reductase evolved from a common ancestor. A pink circle indicates a bootstrap value of more than 70.

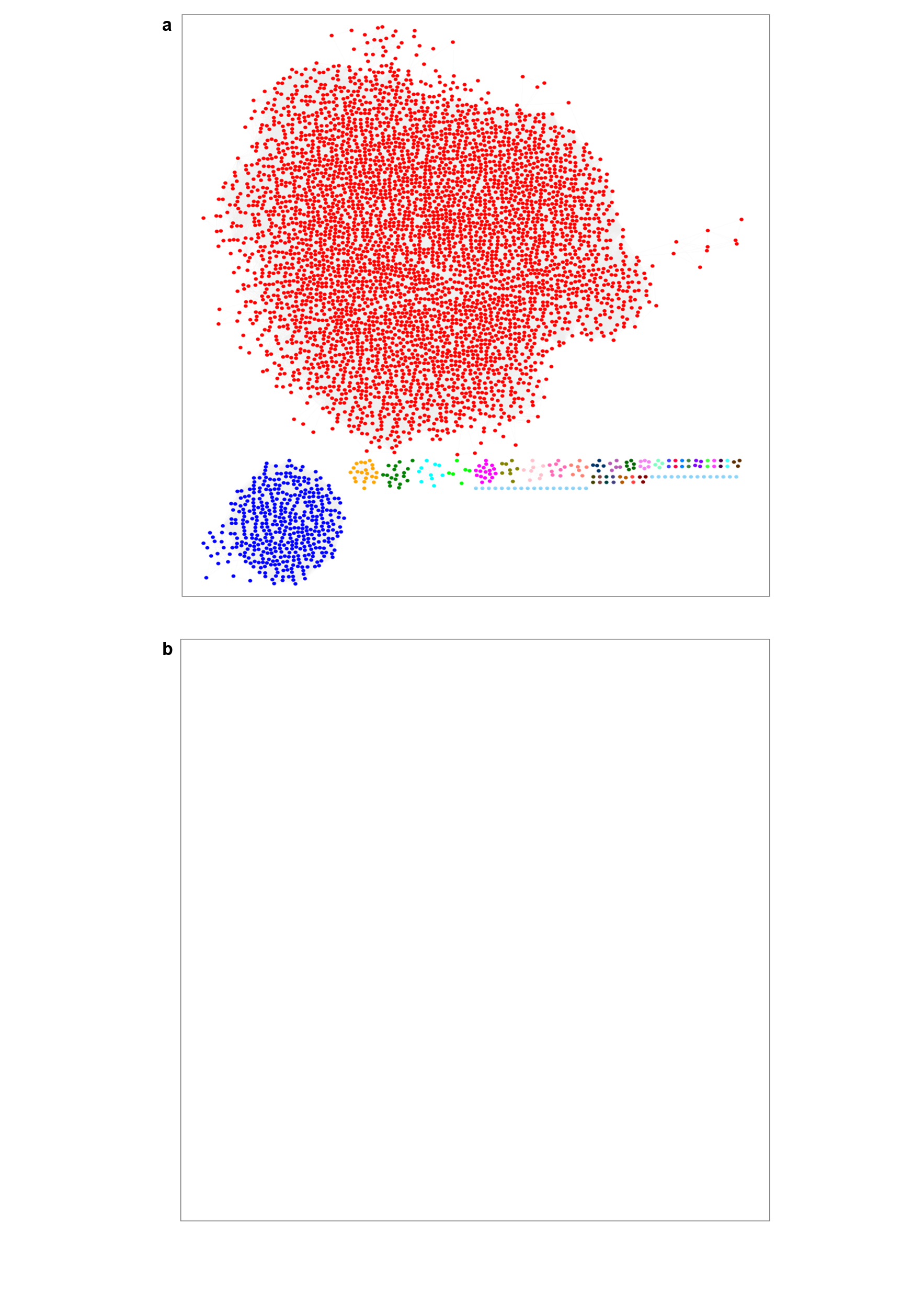

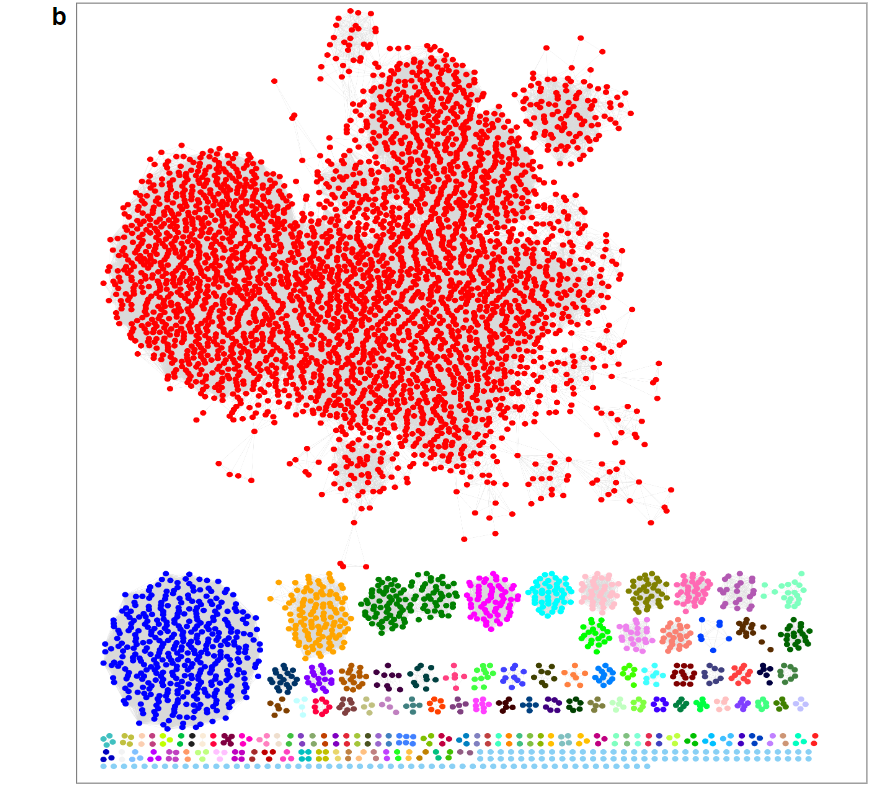

**Fig. S16**

5βChR sequence similarity network. **(a)** A sequence similarity Network (SSN) is built using 5000 proteins from the UniProt with 60% sequence coverage and >50% sequence identity to 5βChR. 4383 proteins out of 5000 are combined into a single cluster, and the rest of the protein sequences are divided into 32 clusters. **(b)** With a 70% threshold and >60% sequence identity to 5βChR, the 5000 proteins are divided into 138 clusters based on functional similarity and increasing sequence identity.

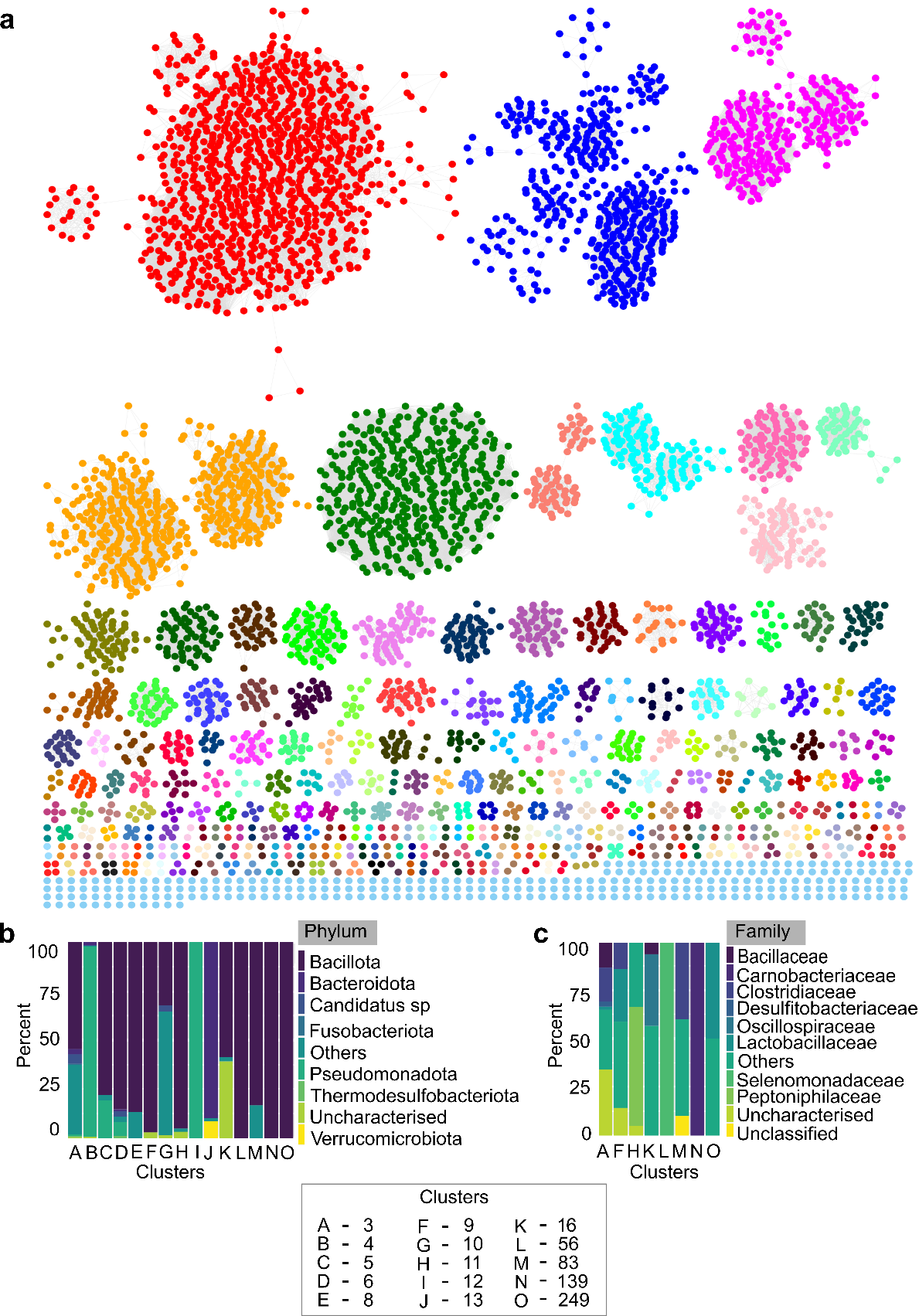

**Fig. S17**

Network maps illustrating the abundance and distribution of the 5βChR gene family across the different microbes. **(a)**. Microbial origin 5βChR protein homologs are retrieved from UniProt for sequence similarity network (SSN) analysis. These networks are visualized by the Enzyme Function Initiative-Enzyme Similarity Tool (EFI-EST) with threshold values of 10^-80^. The proteins are divided into 269 clusters based on sequence identity, structural similarities, and functional profile of 5βChR proteins, highlighting their presence in various microbes. **(b-c)** The graphs depict the taxonomy of microbial 5βChR homologs from the selected clusters, which are prevalent in human microbiome data. The representative sequences of clusters belong to different bacteria belonging to diverse phyla and families.

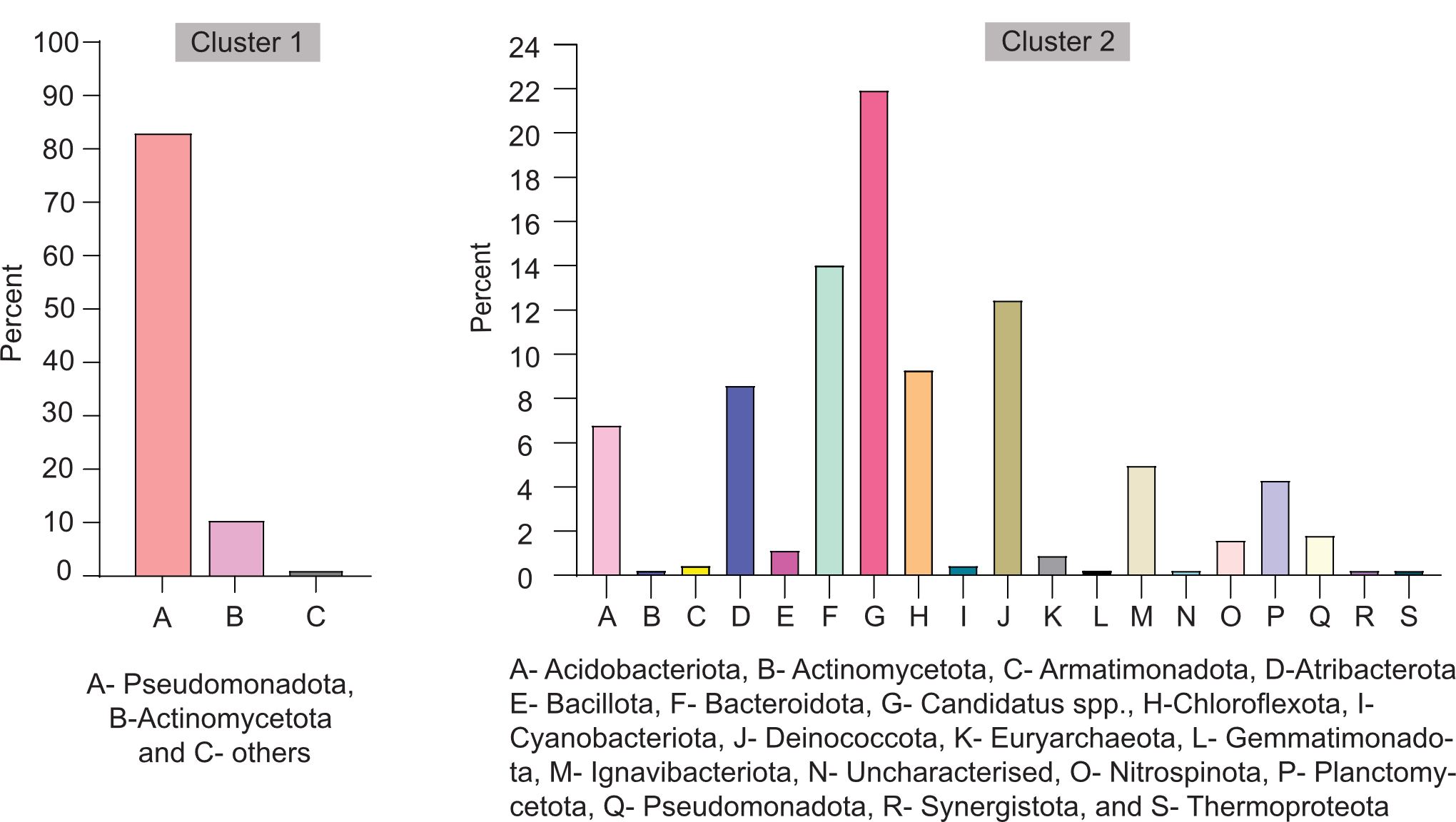

**Fig. S18**

Assessing the taxonomy of 5βChR representatives. The phyla-based classification of clusters 1 and 2 of 5βChR homologs with >80% sequence coverage and >70% sequence identity. The bar graphs show the prevalence of 5βChR protein in various phyla.

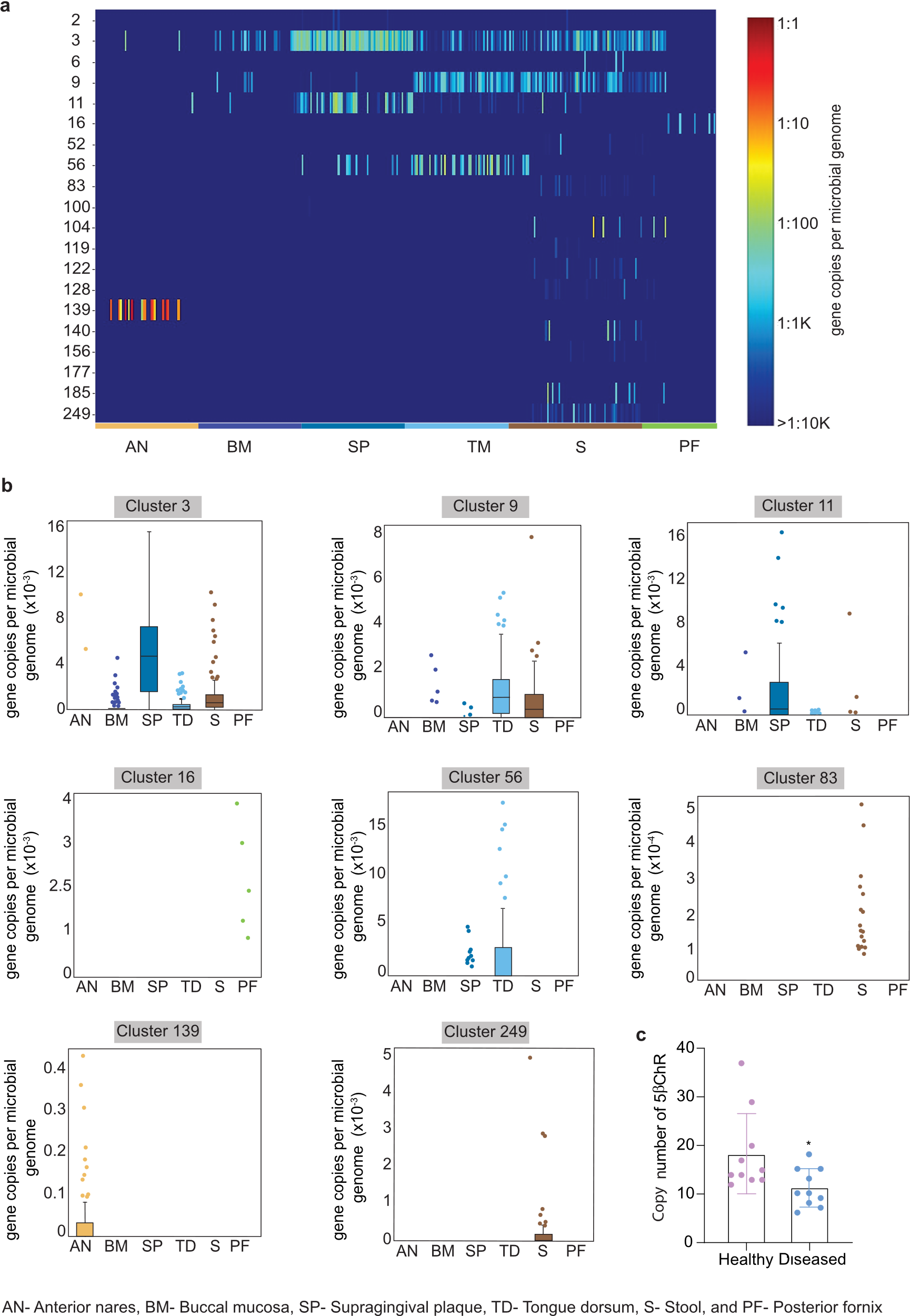

**Fig. S19**

Meta-analysis depicts the abundance of 5βChR homologs in a healthy microbiome. The heat map illustrates the prevalence of 5βChR homologs in the human metagenomic data of 380 healthy individuals. Analysis validates the presence of 20 clusters in the participants' metagenome data.

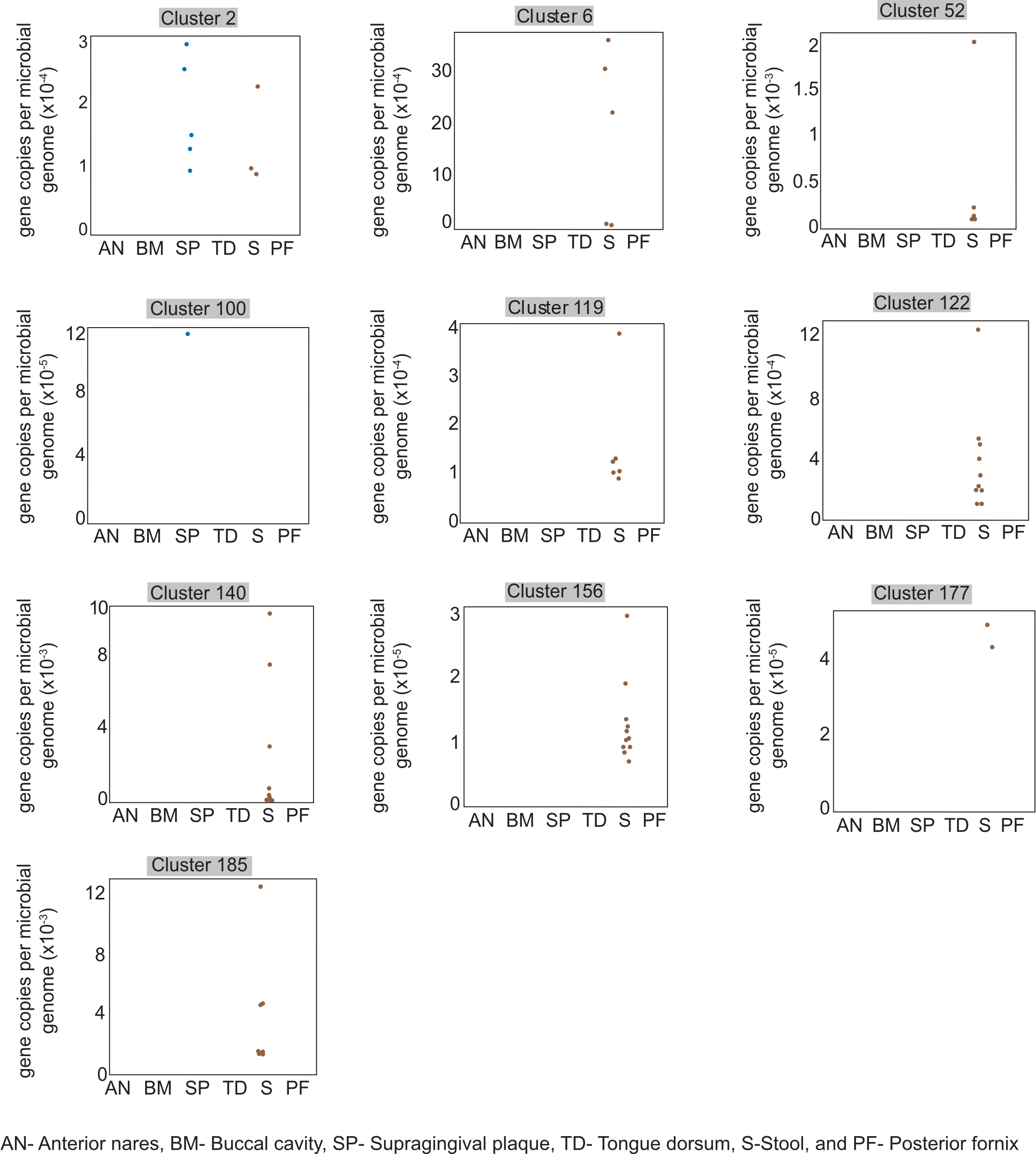

**Fig. S20**

The distribution of different 5βChR homolog clusters in the 380-microbiome data of healthy human participants. The box plot displays the various clusters of 5βChR homologs primarily present in stool samples.

**2. Supplementary Tables**

**Table S1. Primer details used in this study.**

| **Primer name** | **Sequence 5'-3'** |
| --- | --- |
| 5βChR FP | TTATCATATGATGCAAACAGTTGTTATCACGGGGGCC |
| 5βChR RP | TTTATCTCGAGTTATACCTGCTTCGGTTGGCCCGTCG |
| 5βChR RT FP | CGTGGAACCAGAGGAAGTGG |
| 5βChR RT RP | TGGCCCGTCGAAATCTCAAA |
| 16S rRNA RT FP | AGTCACGGCTAACTACGTGC |
| 16S rRNA RT RP | ACCGCCTGCACTCTCTTTAC |
| S139RP | TAACATCTCGgcgGTTTCCGGGATGG |
| S139FP | ACAATGGCGCCGTGTCGT |
| Y152RP | GATGGTGGCCgcgAACGCCACCAAG |
| Y152FP | CCGTAGTCACCACCCATC |
| K156FP | TAACGCCACCgcgGGGGCCATCATC |
| K156RP | TAGGCCACCATCCCGTAG |
| H131 FP | AGACCAACGAgcgGGCGCCATTGTTAACATCTCGTCC |
| H131 RP | AGCATGGTCGGCAGGACC |
| N136 FP | CGCCATTGTTgcgATCTCGTCCGTTTC |
| N136 RP | CCGTGTCGTTGGTCTAGC |
| E172 FP | TGACTACGGTgcgGACCAGATCCG |
| E172 RP | ATCGCCATCGCCCGGGTC |
| P182 FP | CAGCGTGGCCgcgGGTCCAACCA |
| P182 RP | TTGACGCGGATCTGGTCCTCACC |
| P184 FP | GGCCCCGGGTgcgACCAACACCC |
| P184 RP | ACGCTGTTGACGCGGATCTGG |
| T187 FP | TCCAACCAACgcgCCGATGTTCCCGC |
| T187 RP | CCCGGGGCCACGCTGTTG |
| M189 FP | CAACACCCCGgcgTTCCCGCAAGACATGAAGG |
| M189 RP | GTTGGACCCGGGGCCACG |
| F190 FP | CACCCCGATGgcgCCGCAAGACATGAAGGGG |
| F190 RP | TTGGTTGGACCCGGGGCC |
| M194 FP | CCCGCAAGACgcgAAGGGGACCT |
| M194 RP | AACATCGGGGTGTTGGTT |
| F198 FP | GAAGGGGACCgcgGCCAAAAACAG |
| F198 RP | ATGTCTTGCGGGAACATC |

**Table S2. Lactobacillus strains used for phylogenetic analysis.**

**Species**

Lactobacillus_delbrueckii_subsp._bulgaricus

Lactobacillus_delbrueckii_subsp._delbrueckii

Lactobacillus_delbrueckii_subsp._indicus

Lactobacillus_delbrueckii_subsp._Jakobsenii

Lactobacillus_delbrueckii_subsp._lactis

Lactobacillus_delbrueckii_subsp._sunkii

Lactobacillus_acetotolerans

Lactobacillus_acidophilus

Lactobacillus_amylolyticus

Lactobacillus_amylovorus

Lactobacillus_apis

Lactobacillus_bombicola

Lactobacillus_crispatus

Lactobacillus_equicursoris

Lactobacillus_gallinarum

Lactobacillus_gasseri

Lactobacillus_gigeriorum

Lactobacillus_jensenii

Lactobacillus_johnsonii

Lactobacillus_hamsteri

Lactobacillus_helsingborgensis

Lactobacillus_helveticus

Lactobacillus_hominis

Lactobacillus_iners

Lactobacillus_intestinalis

Lactobacillus_kalixensis

Lactobacillus_kefiranofaciens_subsp._kefiranofaciens

Lactobacillus_kefiranofaciens_subsp._kefirgranum

Lactobacillus_kimbladii

Lactobacillus_kitasatonis

Lactobacillus_kullabergensis

Lactobacillus_melliventris

Lactobacillus_mulieris

Lactobacillus_panisapium

Lactobacillus_paragasseri

Lactobacillus_pasteurii

Lactobacillus_psittaci

Lactobacillus_rodentium

Lactobacillus_taiwanensis

Lactobacillus_ultunensis

Lactobacillus_xujianguonis

Amylolactobacillus_amylophilus

Holzapfelia_floricola

Bombilactobacillus_mellifer

Bombilactobacillus_bombi

Bombilactobacillus_mellis

Companilactobacillus_alimentarius

Companilactobacillus_allii

Companilactobacillus_baiquanensis

Companilactobacillus_bobalius

Companilactobacillus_crustorum

Companilactobacillus_farciminis

Companilactobacillus_formosensis

Companilactobacillus_furfuricola

Companilactobacillus_futsaii

Companilactobacillus_ginsenosidimutans

Companilactobacillus_halodurans

Companilactobacillus_heilongjiangensis

Companilactobacillus_huachuanensis

Companilactobacillus_hulinensis

Companilactobacillus_insicii

Companilactobacillus_jidongensis

Companilactobacillus_kedongensis

Companilactobacillus_keshanensis

Companilactobacillus_kimchiensis

Companilactobacillus_kimchii

Companilactobacillus_metriopterae

Companilactobacillus_mindensis

Companilactobacillus_mishanensis

Companilactobacillus_musae

Companilactobacillus_nantensis

Companilactobacillus_nodensis

Companilactobacillus_nuruki

Companilactobacillus_paralimentarius

Companilactobacillus_suantsaicola

Companilactobacillus_tucceti

Companilactobacillus_versmoldensis

Companilactobacillus_zhachilii

Companilactobacillus_zhongbaensis

Lapidilactobacillus_concavus

Lapidilactobacillus_bayanensis

Lapidilactobacillus_dextrinicus

Agrilactobacillus_composti

Agrilactobacillus_yilanensis

Schleiferilactobacillus_perolens

Schleiferilactobacillus_harbinensis

Schleiferilactobacillus_shenzhenensis

Lacticaseibacillus_casei

Lacticaseibacillus_baoqingensis

Lacticaseibacillus_brantae

Lacticaseibacillus_camelliae

Lacticaseibacillus_chiayiensis

Lacticaseibacillus_hulanensis

Lacticaseibacillus_jixianensis

Lacticaseibacillus_manihotivorans

Lacticaseibacillus_nasuensis

Lacticaseibacillus_pantheris

Lacticaseibacillus_paracasei_subsp._paracasei

Lacticaseibacillus_paracasei_subsp._tolerans

Lacticaseibacillus_porcinae

Lacticaseibacillus_rhamnosus

Lacticaseibacillus_saniviri

Lacticaseibacillus_sharpeae

Lacticaseibacillus_songhuajiangensis

Lacticaseibacillus_thailandensis

Paralactobacillus_selangorensis

Latilactobacillus_sakei_subsp._carnosus

Latilactobacillus_sakei_subsp._sakei

Latilactobacillus_curvatus

Latilactobacillus_fuchuensis

Latilactobacillus_graminis

Loigolactobacillus_coryniformis_subsp._coryniformis

Loigolactobacillus_coryniformis_subsp._torquens

Loigolactobacillus_backii

Loigolactobacillus_bifermentans

Loigolactobacillus_iwatensis

Loigolactobacillus_jiayinensis

Loigolactobacillus_rennini

Loigolactobacillus_zhaoyuanensis

Dellaglioa_algida

Liquorilactobacillus_mali

Liquorilactobacillus_aquaticus

Liquorilactobacillus_cacaonum

Liquorilactobacillus_capillatus

Liquorilactobacillus_ghanensis

Liquorilactobacillus_hordei

Liquorilactobacillus_nagelii

Liquorilactobacillus_satsumensis

Liquorilactobacillus_oeni

Liquorilactobacillus_sucicola

Liquorilactobacillus_uvarum

Liquorilactobacillus_vini

Ligilactobacillus_salivarius

Ligilactobacillus_acidipiscis

Ligilactobacillus_agilis

Ligilactobacillus_animalis

Ligilactobacillus_apodemi

Ligilactobacillus_araffinosus

Ligilactobacillus_aviarius

Ligilactobacillus_ceti

Ligilactobacillus_equi

Ligilactobacillus_hayakitensis

Ligilactobacillus_murinus

Ligilactobacillus_pobuzihii

Ligilactobacillus_ruminis

Ligilactobacillus_saerimneri

Lactiplantibacillus_plantarum_subsp._plantarum

Lactiplantibacillus_plantarum_subsp._argentoratensis

Lactiplantibacillus_daoliensis

Lactiplantibacillus_daowaiensis

Lactiplantibacillus_dongliensis

Lactiplantibacillus_fabifermentans

Lactiplantibacillus_herbarum

Lactiplantibacillus_modestisalitolerans

Lactiplantibacillus_mudanjiangensis

Lactiplantibacillus_nangangensis

Lactiplantibacillus_paraplantarum

Lactiplantibacillus_pentosus

Lactiplantibacillus_pingfangensis

Lactiplantibacillus_plajomi

Lactiplantibacillus_songbeiensis

Lactiplantibacillus_xiangfangensis

Furfurilactobacillus_rossiae

Furfurilactobacillus_siliginis

Paucilactobacillus_vaccinostercus

Paucilactobacillus_hokkaidonensis

Paucilactobacillus_kaifaensis

Paucilactobacillus_nenjiangensis

Paucilactobacillus_oligofermentans

Paucilactobacillus_suebicus

Paucilactobacillus_wasatchensis

Limosilactobacillus_fermentum

Limosilactobacillus_antri

Limosilactobacillus_coleohominis

Limosilactobacillus_equigenerosi

Limosilactobacillus_frumenti

Limosilactobacillus_gastricus

Limosilactobacillus_gorillae

Limosilactobacillus_ingluviei

Limosilactobacillus_mucosae

Limosilactobacillus_oris

Limosilactobacillus_panis

Limosilactobacillus_pontis

Limosilactobacillus_reuteri

Limosilactobacillus_secaliphilus

Limosilactobacillus_vaginalis

Secundilactobacillus_malefermentans

Secundilactobacillus_collinoides

Secundilactobacillus_kimchicus

Secundilactobacillus_mixtipabuli

Secundilactobacillus_odoratitofui

Secundilactobacillus_oryzae

Secundilactobacillus_paracollinoides

Secundilactobacillus_pentosiphilus

Secundilactobacillus_silagei

Secundilactobacillus_silagincola

Secundilactobacillus_similis

Levilactobacillus_brevis

Levilactobacillus_acidifarinae

Levilactobacillus_bambusae

Levilactobacillus_cerevisiae

Levilactobacillus_fujinensis

Levilactobacillus_fuyuanensis

Levilactobacillus_hammesii

Levilactobacillus_huananensis

Levilactobacillus_koreensis

Levilactobacillus_lindianensis

Levilactobacillus_mulengensis

Levilactobacillus_namurensis

Levilactobacillus_parabrevis

Levilactobacillus_paucivorans

Levilactobacillus_senmaizukei

Levilactobacillus_spicheri

Levilactobacillus_suantsaii

Levilactobacillus_suantsaiihabitans

Levilactobacillus_tangyuanensis

Levilactobacillus_tongjiangensis

Levilactobacillus_yonginensis

Levilactobacillus_zymae

Fructilactobacillus_fructivorans

Fructilactobacillus_florum

Fructilactobacillus_lindneri

Fructilactobacillus_sanfranciscensis

Acetilactobacillus_jinshanensis

Apilactobacillus_kunkeei

Apilactobacillus_apinorum

Apilactobacillus_micheneri

Apilactobacillus_ozensis

Apilactobacillus_quenuiae

Apilactobacillus_timberlakei

Lentilactobacillus_buchneri

Lentilactobacillus_curieae

Lentilactobacillus_diolivorans

Lentilactobacillus_farraginis

Lentilactobacillus_hilgardii

Lentilactobacillus_kefiri

Lentilactobacillus_kisonensis

Lentilactobacillus_otakiensis

Lentilactobacillus_parabuchneri

Lentilactobacillus_parafarraginis

Lentilactobacillus_parakefiri

Lentilactobacillus_raoultii

Lentilactobacillus_rapi

Lentilactobacillus_senioris

Lentilactobacillus_sunkii

Lapidilactobacillus_achengensis

Lapidilactobacillus_gannanensis

Lapidilactobacillus_mulanensis

Lapidilactobacillus_wuchangensis

Lacticaseibacillus_daqingensis

Lacticaseibacillus_hegangensis

Lacticaseibacillus_suibinensis

Lacticaseibacillus_yichunensis

Loigolactobacillus_binensis

Lactiplantibacillus_garii

Levilactobacillus_angrenensis

**Table S3. List of 5βChR plants homologs**

**Name & Uniprot ID**

Digitalis_lanata_Q6PQJ

Sesamum_indicum_A0A6I9TR48_SESIN (Sesamum_indicum d)

Erythranthe_guttata_A0A022RBG3_ERYGU

Handroanthus_impetiginosus_A0A2G9HCN6_9LAMI (Handroanthus_impetiginosus c)

Sesamum_indicum_A0A6I9UEB3_SESIN

Handroanthus_impetiginosus_A0A2G9HDB4_9LAMI (Handroanthus_impetiginosus b)

Handroanthus_impetiginosus_A0A2G9FZQ9_9LAMI (Handroanthus_impetiginosus d)

Handroanthus_impetiginosus_A0A2G9G080_9LAMI (Handroanthus_impetiginosus a)

Phtheirospermum_japonicum_A0A830BWA5_9LAMI (Phtheirospermum_japonicum a)

Striga_asiatica_A0A5A7PVK4_STRAF

Olea_europaea_subspe_uropaea_A0A8S0T3M5_OLEEU (Olea_europaea f)

Handroanthus_impetiginosus_A0A2G9HCM6_9LAMI (Handroanthus_impetiginosus e)

Olea_europaea_subsp_europaea_A0A8S0UNL0_OLEEU (Olea_europaea d)

Olea_europaea_subsp_europaea_A0A8S0TDW3_OLEEU (Olea_europaea c)

Olea_europaea_subsp_europaea_A0A8S0T0U6_OLEEU (Olea_europaea e)

Dorcoceras_hygrometricum_A0A2Z7BQP4_9LAMI (Dorcoceras_hygrometricum b)

Nepeta_racemosa_ISY1_NEPRA

Sesamum_indicum_A0A6I9TWM7_SESIN (Sesamum_indicum b)

Genlisea_aurea_S8DQY0_9LAMI

Sesamum_indicum_A0A8M8V613_SESINO (Sesamum_indicum a)

Coffea_arabica_A0A6P6WBJ5_COFAR (Coffea_arabica a)

Coffea_canephora_A0A068TPH4_COFCA (Coffea_arabica b)

Coffea_arabica_A0A6P6VU68_COFAR

Phtheirospermum_japonicum_A0A830BZG9_9LAMI (Phtheirospermum_japonicum b)

Salvia_splendens_A0A8X8Y7X1_SALSN (Salvia_splendens a)

Salvia_splendens_A0A8X8Y1P7_SALSN (Salvia_splendens b)

Nicotiana_attenuata_A0A1J6KD04_NICAT

Nicotiana_tabacum_C0SUC5_TOBAC (Nicotiana_tabacum c)

Solanum_lycopersicum_A0A3Q7IFK3_SOLLCOX

Nicotiana_sylvestris_A0A1U7W281_NICSY (Nicotiana_sylvestris b)

Nicotiana_tabacum_A0A1S4C7K6_TOBAC (Nicotiana_tabacum d)

Salvia_splendens­_A0A8X8XA34_SALSN (Salvia_splendens c)

Olea_europaea_subsp_europaea_A0A8S0S199_OLEEU (Olea_europaea a)

Solanum_tuberosum_M1A3I7_SOLTU

Erythranthe_guttata_A0A022RB89_ERYGU

Salvia_splendens_A0A8X8X5Q9_SALSN (Salvia_splendens d)

Solanum_commersonii_A0A9J5WZQ5_SOLCO

Dorcoceras_hygrometricum_A0A2Z7AKE5_9LAMI (Dorcoceras_hygrometricum a)

Capsicum_baccatum_A0A2G2W4K4_CAPBA

Cuscuta_campestris_A0A484KG59_9ASTE

Camellia_sinensis_A0A7J7HHB0_CAMSI

Olea_europaea_subsp_europaea_A0A8S0SEQ7_OLEEU

Cuscuta_australis_ A0A328D8P5_9ASTE

Capsicum_baccatum_A0A2G2XN52_CAPBA

Actinidia_chinensis_var_chinensis_A0A2R6Q5W0_ACTCC (Actinidia_chinensis a)

Juglans_regia_A0A2I4FQH4_JUGRE (Juglans_regia a)

Juglans_regia_A0A2I4FWX2_JUGRE (Juglans_regia c)

Daucus_carota_subsp_sativus_A0A166GIS8_DAUCS

Lactuca_sativa_A0A166GIS8_DAUCS

Actinidia_chinensis_var_chinensis_A0A2R6RGX9_ACTCC (Actinidia_chinensis b)

Carya_illinoinensis_A0A8T1N750_CARIL

Nelumbo_nucifera_A0A1U8 (Nelumbo_nucifera b)

Carpinus_fangiana_A0A5N6RKZ6_9ROSI

Prunus_yedoensis_var_nudiflora_A0A314XHP9_PRUYE

Prunus_dulcis_A0A5E4E997_PRUDU

Cynara_cardunculus_var_scolymus_A0A103XM59_CYNCS

Sesamum_indicum_A0A6I9U2W0_SESIN (Sesamum_indicum c)

Morus_notabilis_W9SGC8_9ROSA

Nelumbo_nucifera_A0A1U8B751_NELNU (Nelumbo_nucifera a)

Prunus_avium_ A0A6P5RCG0_PRUAV

Prunus_persica_A0A251RBQ8_PRUPE

Kingdonia_uniflora_A0A7J7LY63_9MAGN

Durio_zibethinus_A0A6P6B6U9_DURZI (Durio_zibethinus a)

Pyrus_ussuriensis_Pyrus_communis_A0A5N5GW02_9ROSA (Pyrus_ussuriensis a)

Helianthus_annuus_ A0A251TMZ3_HELAN

Abrus_precatorius_A0A8B8LJU1_ABRPR (Abrus_precatorius b)

Pyrus_ussuriensis_Pyrus_communis (Pyrus_ussuriensis b)

Durio_zibethinus_A0A6P5WM40_DURZI (Durio_zibethinus b)

Actinidia_rufa_A0A7J0GN97_9ERIC

Nicotiana_tabacum_A0A1S4B9B5_TOBAC (Nicotiana_tabacum a)

Abrus_precatorius_A0A8B8LHB1_ABRPR (Abrus_precatorius a)

Vigna_radiata_var_radiata_A0A1S3TXR7_VIGRR

Mikania_micrantha_A0A5N6LLS5_9ASTR

Phaseolus_vulgaris­­_V7AWY5_PHAVU

Theobroma_cacao_A0A061FFF0_THECC

Ziziphus_jujuba_A0A6P3ZUB5_ZIZJJ

Nicotiana_sylvestris_A0A1U7WAQ4_NICSY (Nicotiana_sylvestris a)

Nicotiana_tabacum_A0A1S3XWC9_TOBAC (Nicotiana_tabacum b)

Castanea_mollissima_A0A8J4RLX8_9ROSI

Capsicum_annuum_A0A2G3AN45_CAPAN

Manihot_esculenta_A0A2C9VXF4_MANES

Vitis_vinifera_F6H675_VITVI

Hibiscus_syriacus_A0A6A2WIU1_HIBSY (Hibiscus_syriacus b)

Morella_rubra_A0A6A1W9A5_9ROSI

Pyrus_ussuriensis_Pyrus_communis_A0A5N5I733_9ROSA

Tripterygium_wilfordii_A0A7J7CZC2_TRIWF

Juglans_regia_A0A2I4HD46_JUGRE (Juglans_regia c)

Prunus_avium_A0A6P5RRI7_PRUAV

Cucurbita_maxima_A0A6J1J0N7_CUCMA

Populus_alba_A0A4U5N810_POPAL

Castanea_mollissima_A0A8J4RN14_9ROSI

Phaseolus_angularis_A0A0L9UHR8_PHAAN

Prunus_avium_A0A6P5RFJ5_PRUAV

Salix_brachista_A0A5N5K873_9ROSI

Momordica_charantia_A0A6J1DZW3_MOMCH

Citrus_clementina_V4T9V0_CITCL

Hibiscus_syriacus_A0A6A2WII9_HIBSY (Hibiscus_syriacus a)

Prunus_armeniaca_A0A6J5W5F7_PRUAR

Medicago_truncatula_G8A2B2_MEDTR

**Table S4. NCBI accession number of metagenome samples.**

| **Metagenome ID** | **Body Site** |
| --- | --- |
| SRS011061 | stool |
| SRS011090 | buccal mucosa |
| SRS011098 | supragingival plaque |
| SRS011126 | supragingival plaque |
| SRS011132 | anterior nares |
| SRS011134 | stool |
| SRS011140 | tongue dorsum |
| SRS011144 | buccal mucosa |
| SRS011152 | supragingival plaque |
| SRS011239 | stool |
| SRS011243 | tongue dorsum |
| SRS011247 | buccal mucosa |
| SRS011255 | supragingival plaque |
| SRS011263 | anterior nares |
| SRS011269 | posterior fornix |
| SRS011271 | stool |
| SRS011302 | stool |
| SRS011306 | tongue dorsum |
| SRS011310 | buccal mucosa |
| SRS011343 | supragingival plaque |
| SRS011355 | posterior fornix |
| SRS011397 | anterior nares |
| SRS011405 | stool |
| SRS011452 | stool |
| SRS011529 | stool |
| SRS011584 | posterior fornix |
| SRS011586 | stool |
| SRS012273 | stool |
| SRS012279 | tongue dorsum |
| SRS012281 | buccal mucosa |
| SRS012285 | supragingival plaque |
| SRS012291 | anterior nares |
| SRS012294 | posterior fornix |
| SRS012663 | anterior nares |
| SRS012902 | stool |
| SRS013155 | anterior nares |
| SRS013158 | stool |
| SRS013164 | tongue dorsum |
| SRS013170 | supragingival plaque |
| SRS013215 | stool |
| SRS013234 | tongue dorsum |
| SRS013239 | buccal mucosa |
| SRS013252 | supragingival plaque |
| SRS013269 | anterior nares |
| SRS013476 | stool |
| SRS013502 | tongue dorsum |
| SRS013506 | buccal mucosa |
| SRS013521 | stool |
| SRS013533 | supragingival plaque |
| SRS013542 | posterior fornix |
| SRS013637 | anterior nares |
| SRS013687 | stool |
| SRS013705 | tongue dorsum |
| SRS013711 | buccal mucosa |
| SRS013723 | supragingival plaque |
| SRS013800 | stool |
| SRS013818 | tongue dorsum |
| SRS013825 | buccal mucosa |
| SRS013836 | supragingival plaque |
| SRS013876 | anterior nares |
| SRS013879 | tongue dorsum |
| SRS013881 | buccal mucosa |
| SRS013945 | buccal mucosa |
| SRS013949 | supragingival plaque |
| SRS013951 | stool |
| SRS013956 | anterior nares |
| SRS014124 | tongue dorsum |
| SRS014126 | buccal mucosa |
| SRS014235 | stool |
| SRS014271 | tongue dorsum |
| SRS014287 | stool |
| SRS014313 | stool |
| SRS014459 | stool |
| SRS014464 | anterior nares |
| SRS014470 | tongue dorsum |
| SRS014472 | buccal mucosa |
| SRS014476 | supragingival plaque |
| SRS014494 | posterior fornix |
| SRS014573 | tongue dorsum |
| SRS014575 | buccal mucosa |
| SRS014578 | supragingival plaque |
| SRS014613 | stool |
| SRS014629 | posterior fornix |
| SRS014682 | anterior nares |
| SRS014683 | stool |
| SRS014684 | tongue dorsum |
| SRS014686 | buccal mucosa |
| SRS014690 | supragingival plaque |
| SRS014888 | tongue dorsum |
| SRS014890 | buccal mucosa |
| SRS014894 | supragingival plaque |
| SRS014901 | anterior nares |
| SRS014923 | stool |
| SRS014979 | stool |
| SRS015038 | tongue dorsum |
| SRS015040 | buccal mucosa |
| SRS015044 | supragingival plaque |
| SRS015051 | anterior nares |
| SRS015054 | posterior fornix |
| SRS015133 | stool |
| SRS015154 | buccal mucosa |
| SRS015158 | supragingival plaque |
| SRS015168 | posterior fornix |
| SRS015190 | stool |
| SRS015209 | tongue dorsum |
| SRS015215 | supragingival plaque |
| SRS015217 | stool |
| SRS015225 | posterior fornix |
| SRS015264 | stool |
| SRS015269 | anterior nares |
| SRS015272 | tongue dorsum |
| SRS015274 | buccal mucosa |
| SRS015278 | supragingival plaque |
| SRS015369 | stool |
| SRS015374 | buccal mucosa |
| SRS015378 | supragingival plaque |
| SRS015395 | tongue dorsum |
| SRS015425 | posterior fornix |
| SRS015430 | anterior nares |
| SRS015434 | tongue dorsum |
| SRS015436 | buccal mucosa |
| SRS015440 | supragingival plaque |
| SRS015450 | anterior nares |
| SRS015470 | supragingival plaque |
| SRS015537 | tongue dorsum |
| SRS015574 | supragingival plaque |
| SRS015578 | stool |
| SRS015640 | anterior nares |
| SRS015644 | tongue dorsum |
| SRS015646 | buccal mucosa |
| SRS015650 | supragingival plaque |
| SRS015663 | stool |
| SRS015745 | buccal mucosa |
| SRS015752 | anterior nares |
| SRS015755 | supragingival plaque |
| SRS015762 | tongue dorsum |
| SRS015782 | stool |
| SRS015893 | tongue dorsum |
| SRS015895 | buccal mucosa |
| SRS015899 | supragingival plaque |
| SRS015921 | buccal mucosa |
| SRS015937 | anterior nares |
| SRS015941 | tongue dorsum |
| SRS015947 | supragingival plaque |
| SRS015960 | stool |
| SRS015989 | supragingival plaque |
| SRS015996 | anterior nares |
| SRS016002 | tongue dorsum |
| SRS016018 | stool |
| SRS016033 | anterior nares |
| SRS016037 | tongue dorsum |
| SRS016039 | buccal mucosa |
| SRS016043 | supragingival plaque |
| SRS016056 | stool |
| SRS016086 | tongue dorsum |
| SRS016088 | buccal mucosa |
| SRS016092 | supragingival plaque |
| SRS016095 | stool |
| SRS016111 | posterior fornix |
| SRS016188 | anterior nares |
| SRS016191 | posterior fornix |
| SRS016196 | buccal mucosa |
| SRS016200 | supragingival plaque |
| SRS016203 | stool |
| SRS016225 | tongue dorsum |
| SRS016267 | stool |
| SRS016292 | anterior nares |
| SRS016297 | buccal mucosa |
| SRS016319 | tongue dorsum |
| SRS016331 | supragingival plaque |
| SRS016335 | stool |
| SRS016342 | tongue dorsum |
| SRS016349 | buccal mucosa |
| SRS016360 | supragingival plaque |
| SRS016434 | anterior nares |
| SRS016495 | stool |
| SRS016501 | tongue dorsum |
| SRS016503 | buccal mucosa |
| SRS016513 | anterior nares |
| SRS016516 | posterior fornix |
| SRS016529 | tongue dorsum |
| SRS016533 | buccal mucosa |
| SRS016553 | anterior nares |
| SRS016559 | posterior fornix |
| SRS016569 | tongue dorsum |
| SRS016575 | supragingival plaque |
| SRS016581 | anterior nares |
| SRS016585 | stool |
| SRS016600 | buccal mucosa |
| SRS016746 | supragingival plaque |
| SRS016752 | anterior nares |
| SRS016753 | stool |
| SRS016954 | stool |
| SRS016989 | stool |
| SRS017013 | buccal mucosa |
| SRS017025 | supragingival plaque |
| SRS017044 | anterior nares |
| SRS017080 | buccal mucosa |
| SRS017103 | stool |
| SRS017120 | tongue dorsum |
| SRS017127 | buccal mucosa |
| SRS017139 | supragingival plaque |
| SRS017156 | anterior nares |
| SRS017191 | stool |
| SRS017209 | tongue dorsum |
| SRS017215 | buccal mucosa |
| SRS017227 | supragingival plaque |
| SRS017244 | anterior nares |
| SRS017247 | stool |
| SRS017304 | supragingival plaque |
| SRS017307 | stool |
| SRS017433 | stool |
| SRS017439 | tongue dorsum |
| SRS017441 | buccal mucosa |
| SRS017445 | supragingival plaque |
| SRS017451 | anterior nares |
| SRS017497 | posterior fornix |
| SRS017511 | supragingival plaque |
| SRS017520 | posterior fornix |
| SRS017521 | stool |
| SRS017533 | tongue dorsum |
| SRS017537 | buccal mucosa |
| SRS017687 | buccal mucosa |
| SRS017691 | supragingival plaque |
| SRS017697 | anterior nares |
| SRS017700 | posterior fornix |
| SRS017701 | stool |
| SRS017713 | tongue dorsum |
| SRS017808 | tongue dorsum |
| SRS017810 | buccal mucosa |
| SRS017814 | supragingival plaque |
| SRS017820 | anterior nares |
| SRS017821 | stool |
| SRS018133 | stool |
| SRS018145 | tongue dorsum |
| SRS018149 | buccal mucosa |
| SRS018157 | supragingival plaque |
| SRS018300 | tongue dorsum |
| SRS018312 | anterior nares |
| SRS018329 | buccal mucosa |
| SRS018337 | supragingival plaque |
| SRS018351 | stool |
| SRS018357 | tongue dorsum |
| SRS018359 | buccal mucosa |
| SRS018369 | anterior nares |
| SRS018394 | supragingival plaque |
| SRS018427 | stool |
| SRS018439 | tongue dorsum |
| SRS018463 | anterior nares |
| SRS018573 | supragingival plaque |
| SRS018575 | stool |
| SRS018585 | anterior nares |
| SRS018591 | tongue dorsum |
| SRS018656 | stool |
| SRS018661 | buccal mucosa |
| SRS018665 | supragingival plaque |
| SRS018671 | anterior nares |
| SRS018739 | tongue dorsum |
| SRS018769 | posterior fornix |
| SRS018778 | supragingival plaque |
| SRS018784 | anterior nares |
| SRS018791 | tongue dorsum |
| SRS018817 | stool |
| SRS019215 | anterior nares |
| SRS019219 | tongue dorsum |
| SRS019221 | buccal mucosa |
| SRS019225 | supragingival plaque |
| SRS019267 | stool |
| SRS019327 | tongue dorsum |
| SRS019329 | buccal mucosa |
| SRS019333 | supragingival plaque |
| SRS019339 | anterior nares |
| SRS019379 | posterior fornix |
| SRS019386 | anterior nares |
| SRS019387 | supragingival plaque |
| SRS019389 | tongue dorsum |
| SRS019391 | buccal mucosa |
| SRS019397 | stool |
| SRS019587 | buccal mucosa |
| SRS019591 | supragingival plaque |
| SRS019597 | anterior nares |
| SRS019600 | posterior fornix |
| SRS019601 | stool |
| SRS019607 | tongue dorsum |
| SRS019968 | stool |
| SRS019974 | tongue dorsum |
| SRS019976 | buccal mucosa |
| SRS019980 | supragingival plaque |
| SRS019986 | anterior nares |
| SRS019989 | posterior fornix |
| SRS020220 | tongue dorsum |
| SRS020226 | supragingival plaque |
| SRS020232 | anterior nares |
| SRS020233 | stool |
| SRS020328 | stool |
| SRS020334 | tongue dorsum |
| SRS020336 | buccal mucosa |
| SRS020340 | supragingival plaque |
| SRS020349 | posterior fornix |
| SRS020386 | anterior nares |
| SRS020856 | tongue dorsum |
| SRS020858 | buccal mucosa |
| SRS020862 | supragingival plaque |
| SRS020868 | anterior nares |
| SRS020869 | stool |
| SRS022137 | stool |
| SRS022143 | tongue dorsum |
| SRS022145 | buccal mucosa |
| SRS022149 | supragingival plaque |
| SRS022158 | posterior fornix |
| SRS022530 | tongue dorsum |
| SRS022532 | buccal mucosa |
| SRS022536 | supragingival plaque |
| SRS022719 | tongue dorsum |
| SRS022721 | buccal mucosa |
| SRS022725 | supragingival plaque |
| SRS022734 | posterior fornix |
| SRS023346 | stool |
| SRS023352 | tongue dorsum |
| SRS023354 | buccal mucosa |
| SRS023358 | supragingival plaque |
| SRS042428 | posterior fornix |
| SRS042457 | buccal mucosa |
| SRS042643 | tongue dorsum |
| SRS043001 | stool |
| SRS043646 | buccal mucosa |
| SRS043663 | tongue dorsum |
| SRS043755 | supragingival plaque |
| SRS044373 | tongue dorsum |
| SRS045004 | stool |
| SRS045049 | buccal mucosa |
| SRS045254 | buccal mucosa |
| SRS045262 | buccal mucosa |
| SRS045313 | supragingival plaque |
| SRS045713 | stool |
| SRS046344 | anterior nares |
| SRS047824 | tongue dorsum |
| SRS048164 | stool |
| SRS048719 | buccal mucosa |
| SRS049389 | tongue dorsum |
| SRS049712 | stool |
| SRS049900 | stool |
| SRS049959 | stool |
| SRS050007 | buccal mucosa |
| SRS050025 | anterior nares |
| SRS050029 | buccal mucosa |
| SRS050184 | posterior fornix |
| SRS050244 | tongue dorsum |
| SRS050628 | buccal mucosa |
| SRS050752 | stool |
| SRS051244 | supragingival plaque |
| SRS051505 | posterior fornix |
| SRS051613 | anterior nares |
| SRS051941 | supragingival plaque |
| SRS052227 | tongue dorsum |
| SRS052330 | posterior fornix |
| SRS052590 | anterior nares |
| SRS052604 | supragingival plaque |
| SRS052697 | stool |
| SRS052876 | supragingival plaque |
| SRS053335 | stool |
| SRS053398 | stool |
| SRS053437 | anterior nares |
| SRS053854 | tongue dorsum |
| SRS054061 | anterior nares |
| SRS054590 | stool |
| SRS054653 | supragingival plaque |
| SRS054687 | tongue dorsum |
| SRS054956 | stool |
| SRS055118 | buccal mucosa |
| SRS055401 | supragingival plaque |
| SRS055426 | tongue dorsum |
| SRS056323 | tongue dorsum |
| SRS056695 | posterior fornix |
| SRS057539 | tongue dorsum |
| SRS057791 | tongue dorsum |
| SRS057807 | posterior fornix |
| SRS058053 | supragingival plaque |
| SRS058213 | anterior nares |
| SRS058808 | supragingival plaque |
